# The role of aging on endothelial cell-cell junctions and pulmonary microvascular permeability

**DOI:** 10.1101/2025.07.29.667420

**Authors:** Aminmohamed Manji, Lefeng Wang, Cynthia Pape, Sanjay Mehta, Preya Patel, Samuel-Caleb Yeung, Eric K. Patterson, Antoine Dufour, Daniel Young, Ruud A.W. Veldhuizen, Sean E. Gill

## Abstract

Lung injury leads to pulmonary microvascular endothelial cell (PMVEC) damage, disruption of cell-cell junctions, and increased permeability. Previously, we demonstrated in a mechanical ventilation-induced model of lung injury that aging exacerbated pulmonary microvascular permeability. Based on this, we hypothesized that aging was associated with increased PMVEC barrier dysfunction due to impaired cell-cell junction integrity. PMVEC were isolated from young and aged mice and cultured to confluence *in vitro*. Barrier function and cell-cell junction integrity were assessed through electric cell-substrate impedance sensing, XPerT permeability assay, immunofluorescence, and western blot analysis. Further studies were conducted to examine alterations in the proteome, markers of inflammation, and actin cytoskeleton organization. To model injurious conditions, PMVEC were stimulated with inflammatory cytokines; permeability and actin cytoskeletal alterations were subsequently assessed. We observed increased basal permeability in PMVEC from aged mice, which was associated with disrupted cell-surface localization of the adherens junction protein, vascular endothelial (VE)-cadherin. Protein abundance of VE-cadherin was significantly increased with age; however, levels of the adapter protein, *γ*-catenin, and the tight junction protein, claudin-5, were decreased. Measures of inflammation, including cytokine expression and cell surface abundance of adhesion molecules, did not differ with age. Alterations in actin cytoskeleton organization, characterized by augmented presence of actin stress fibers, were observed in aged PMVEC. Under inflammatory conditions, permeability and actin stress fiber formation were exacerbated with age. It is concluded that aging predisposes PMVEC to elevated injury, due to inherent deficiencies in cell-cell junctions and barrier function, potentially mediated through altered actin cytoskeleton organization.

**New and Noteworthy:** Compared with pulmonary microvascular endothelial cells (PMVEC) from young mice, PMVEC isolated from aged mice had higher permeability, which was directly associated with impairments in cell-cell junctions. The higher permeability in aged PMVEC was not associated with augmented inflammatory signaling but was associated with actin cytoskeletal alterations. Following an inflammatory insult, PMVEC from aged mice had further exacerbated permeability. These findings may begin to highlight why older patients exhibit higher mortality during lung injury.

## Introduction

Under healthy conditions, the pulmonary microvasculature serves as a selectively permeable barrier that prevents the leak of fluids and large macromolecules from circulation into the lung interstitium (1). However, during lung injury, such as with mechanical ventilation-induced lung injury or acute respiratory distress syndrome (ARDS), widespread inflammation occurs within the lungs, leading to activation and injury of the pulmonary microvascular endothelial cells (PMVEC) (2–5). This results in microvascular barrier dysfunction and the accumulation of protein-rich edema fluid within the alveolar spaces, leading to impaired gas exchange and respiratory failure (6).

A critical component of the normal pulmonary microvascular barrier is the presence of inter-PMVEC cell-cell junctions (7). This includes adherens junctions, which provide mechanical anchorage between adjacent cells, and tight junctions, which seal paracellular gaps (1, 8). Both junctions are necessary to limit permeability and maintain PMVEC barrier function (7). The junctions are composed of transmembrane proteins, including vascular endothelial (VE)-cadherin for the adherens junctions, and claudin-5 for the tight junctions (8). Additionally, the transmembrane proteins are linked with the underlying actin cytoskeleton via several adapter proteins, such as *α*-catenin, *β*-catenin, and *γ*-catenin for the adherens junctions (8). Binding of the junctional proteins to the cytoskeleton, particularly to thick cortical actin bundles, is crucial for maintaining junctional integrity (9–11). Damage to the microvasculature, due to the inflammatory processes taking place during lung injury, results in disruption of endothelial cell-cell junctions, formation of actin stress fibers that lead to cell retraction, and cell apoptosis (2–4, 12). This subsequently leads to endothelial barrier dysfunction, gap formation within the microvascular walls, and the characteristic augmentation in microvascular macromolecular permeability that is a hallmark of ARDS.

While pulmonary microvascular hyperpermeability is a common feature during lung injury and occurs in all ARDS patients, the elderly population is disproportionately impacted. In fact, the incidence of ARDS rises almost exponentially for every decade of age, and both severity and mortality are known to be exacerbated in elderly ARDS patients (13–19). Enhancing our understanding of the role that aging plays in contributing to the increased morbidity and mortality during lung injury will be paramount in improving outcomes in older patients.

Our group has previously demonstrated an age-associated predisposition to increased pulmonary microvascular permeability in a model of direct lung injury, induced by mechanical ventilation, which was associated with altered endothelial cell-cell junction transcriptional signaling (20). Based on this, our current study aimed to further assess the role of aging on pulmonary microvascular barrier function in an *in vitro* model to directly examine mechanisms at the level of PMVEC. While other studies have examined endothelial barrier function and mechanisms associated with cell-cell junction disruption *in vitro*, they have relied on the induction of senescence as their model of aging (21–28). Examples of induction of senescence have included repeated cell passaging (23–26, 28), activation of certain signaling pathways (21), and cell-mediated injury (22, 27), and while useful, these models may not accurately capture the biomolecular processes taking place with natural aging. Notably, the current study was performed in microvascular endothelial cells that were isolated directly from the lungs of young and aged mice, with subsequent analyses performed to assess barrier function. We hypothesized that aging would be associated with increased PMVEC barrier dysfunction due to impaired cell-cell junction integrity.

## Methods

### Model of aging and pulmonary microvascular endothelial cell (PMVEC) isolation

All animal procedures were performed in accordance with the Canadian Council on Animal Care guidelines and approved by the Western University Animal Care Committee (Approval #2020-054). C57BL/6 male mice from Charles River laboratories were aged to 22 months under controlled conditions (standard light-dark cycles, temperature, humidity, and group-housing) within our institutional animal facility. Obesity has been shown to be associated with augmented proinflammatory responses, changes in respiratory mechanics, elevated risk of ARDS, and vascular damage (29–32). Based on this, the mice were fed a food-restricted diet with a gradual reduction to 70% ad libitum to maintain body weight, as described previously (33). For these studies, PMVEC were isolated from the lungs of young (3-month-old) and aged (18-22-month-old) mice as previously described (34, 35). Briefly, excised lungs were finely minced, digested with collagenase, and the resulting suspension incubated with magnetic microbeads (Dynal Biotech Inc., Lake Success, NY) coated with anti-CD31 (platelet endothelial cell adhesion molecule) antibodies (BD Pharmingen, Franklin Lakes, NJ). Bead-bound PMVEC were isolated, washed, and resuspended in complete Dulbecco’s Modified Eagle Medium (DMEM; Gibco 11885084, 1 g/L D-glucose, L-glutamine, 110 mg/L sodium pyruvate, phenol red) supplemented with 20% fetal bovine serum (FBS; Gibco 12483-020), Penicillin-Streptomycin (Gibco 15140122), and 4-(2-hydroxyethyl)piperazine-1-ethane-sulfonic acid (HEPES; Gibco 15630-080). Cells were seeded onto gelatin-coated flasks and expanded to ∼90 % confluence. Purity was confirmed by staining with antibodies against CD31, CD34, CD146, and CD202b, conjugated to Pacific Blue, Phycoerythrin, Fluorescein Isothiocyanate, or Allophycocyanin, respectively (VWR Scientific Inc., Radnor, PA), and analyzed on an easyCyte Guava 12HT flow cytometer (Millipore, Billerica, MA, USA). This protocol consistently yielded 99% homogeneous PMVEC populations. Cells from passages 4–10 were used for experiments, with growth media being changed every two days. All experiments were performed with independent PMVEC isolations from 2–3 young and aged mice, with cells given 72 hours to grow to confluence. Cells were isolated exclusively from male mice for practical reasons related to maintenance of our aging mouse colony during the corona virus disease 2019 (COVID-19) pandemic. The limitation of using only male mice is addressed in the limitations of the study.

### *In vitro* stimulation of PMVEC

These studies aimed to assess the impact of aging on PMVEC barrier function under basal conditions as well as inflammatory conditions that model ARDS *in vitro*. An equimolar mixture of pro-inflammatory cytokines (cytomix), including tumor necrosis factor (TNF)*α*, interleukin (IL)1*β*, and interferon (IFN)*γ*, diluted in phosphate-buffered saline (PBS), was used to model the inflammatory conditions. These cytokines are relevant to inflammatory diseases such as sepsis or COVID-19, which are known to induce ARDS (36, 37). Cells were treated by adding cytomix to the growth media at a final concentration of 30 ng/mL for each cytokine, as has been established previously in our lab using mouse PMVEC (35). Following 72 hours of growth to confluence, stimulation occurred for 4 hours to represent an acute time-point that our lab has previously demonstrated to cause increases in PMVEC permeability, as well as a more long-term time-point of 24 hours, which has been shown to induce PMVEC cell death (35). For control conditions, an equal volume of PBS was added to the cell culture media.

### Real-time assessment of PMVEC monolayer barrier integrity

PMVEC were seeded in duplicates at 5 x 10^4^ cells per well in eight-well arrays (Applied Biophysics, 8W10E+ PET) coated with 1% gelatin. Cells were grown in complete DMEM, and resistance was continuously monitored at 4000 Hz with an electric cell-substrate impedance sensing (ECIS) instrument (Model Zθ, Applied Biophysics).

### PMVEC monolayer paracellular permeability visualization

To directly assess the relationship between PMVEC monolayer permeability and junctional disruption, a fluorescence technique was employed involving visualization of localized leak based on a modification of the published XPerT permeability assay (38). For these experiments, PMVEC were seeded at a density of 2.5 - 3 x 10^4^ cells in 48-well plates that were coated with biotin-conjugated gelatin overnight at 4℃. Oregon green 488-conjugated NeutrAvidin (Invitrogen A6374) was added to the cell culture media at a final concentration of 25 µg/mL for 3 minutes. The cells were then fixed with 200 µL of ice-cold methanol for 15 minutes, followed by three washes with PBS to remove residual methanol and remaining unbound avidin. Unlike the initial study describing the XPerT assay (38), the three PBS washes were performed following fixation rather than after the addition of NeutrAvidin to prevent disturbances to the PMVEC and induction of artificial junctional disruption. Our group has previously shown in human PMVEC that almost all paracellular leak was associated with VE-cadherin disruption (39). Furthermore, disruption of the adherens junction induced by compromised VE-cadherin adhesion has been shown to be the leading cause of tissue edema in many pathological conditions (1, 40–42). Based on this, examination of NeutrAvidin leak was performed concurrently with VE-cadherin staining. Immunocytochemistry for VE-cadherin staining was carried out as described below. Quantification of NeutrAvidin leak was performed by measuring fluorescence in ImageJ; values were normalized to average baseline levels in the PMVEC isolated from young mice.

### Immunocytochemistry of junctional proteins

Cells were fixed as described above, permeabilized with 0.1% Triton X-100 (VWR), and blocked using 3% bovine serum albumin (BSA) in PBS. The cells were then incubated with antibodies against VE-cadherin (rabbit polyclonal, 1:200 dilution, Abcam 33168) or claudin-5 (mouse monoclonal, 1:100 dilution, Invitrogen 35-2500) in 1% BSA in PBS, followed by incubation with Alexa Fluor 594-conjugated IgG secondary antibody (donkey anti-rabbit polyclonal, 1:500 dilution, Invitrogen A11037) or Alexa Fluor 488-conjugated IgG1 secondary antibody (goat anti-mouse polyclonal, 1:500 dilution, Invitrogen A21121). Nuclei were visualized by counterstaining with Hoechst 33342 in PBS (1:5000 dilution, Invitrogen H3570). Cells were then imaged at 200x magnification by fluorescent microscopy (Zeiss Axiovert 200M Inverted Microscope; Carl Zeiss Canada, Toronto, Canada) with exposure times and contrast levels kept consistent between the young and aged PMVEC. Four to five images were taken at random locations within each experimental well, excluding the very edges of the well. Image amalgamation and analysis were performed using ImageJ.

### PMVEC protein isolation and western blot analysis

PMVEC from young and aged mice were cultured at a density of 2.5 x 10^5^ cells on 6-well plates coated with 1% gelatin. On the experimental day, media from the wells was aspirated and the cells were washed with ice-cold PBS. A 200 µL mixture of ice-cold Cell Lytic M lysis buffer (Sigma-Aldrich C2978), protease inhibitor cocktail (Sigma-Aldrich P8340), and 2mM sodium orthovanadate (phosphatase inhibitor) was added to each well for 20 minutes on ice, with rocking every 5 minutes. Cell lysates were then sonicated for 10 seconds (power output of 15% on the Cole Parmer 4710 Series Ultrasonic Homogenizer). Samples were centrifuged at 20,000 x g for 15 minutes at 4℃, the supernatant transferred to a new chilled tube, and protein concentrations assessed using a detergent-compatible protein assay (Bio-Rad 5000119) and BSA standards in Cell Lytic M. Protein concentrations were normalized across the samples and gel loading buffer (60mM Tris pH 6.8, 25% glycerol, 2% sodium dodecyl sulfate [SDS], 14.4*µ*M *β*-mercaptoethanol, 0.1% BromoPhenol Blue, 0.1% H_2_O) was added. Samples were stored at −20℃ until ready to use for western blotting.

For western blot analysis, samples were thawed on ice, heated to 95℃ for 5 minutes, then loaded onto a 10% or 12% polyacrylamide gel for electrophoresis for approximately 1-2 hours (depending on protein of interest). Proteins were then transferred for 45 minutes to 1 hour (depending on protein of interest) to a polyvinylidene fluoride membrane (Bio-Rad 162-0177). Membranes were blocked using 3% BSA or 5% skim milk, both diluted in tris-buffered saline with 0.05% tween-20 (TBS-T) for 1 hour. Following blocking, the membranes were incubated overnight at 4℃ with primary antibodies targeting VE-cadherin (IgG rabbit polyclonal, 1:2,000, Abcam 33168), *α*-catenin (IgG1 mouse monoclonal, 1:1,000, BD Transduction Laboratories 610194), *β*-catenin (IgG1 mouse monoclonal, 1:4,000, BD Transduction Laboratories Biosciences 610154), *γ*-catenin (IgG2a mouse monoclonal, 1:6,000, BD Transduction Laboratories Biosciences 610254), claudin-5 (IgG1 mouse monoclonal, 1:1,000, Invitrogen 35-2500), and glyceraldehyde-3-phosphate dehydrogenase (GAPDH) (IgG1 mouse monoclonal, 1:5,000, Thermo Fisher Scientific MA5-15738), which was used as a loading control. All antibodies were diluted in 1% BSA in TBS-T, with the exception of claudin-5, which was diluted in 5% skim milk in TBS-T. Following antibody incubation, membranes were washed with TBS-T, then incubated with their appropriate secondary antibodies, including goat anti-mouse polyclonal IgG1 horseradish peroxidase (HRP) (1:10,000, Abcam Ab98693), goat anti-mouse polyclonal IgG2a HRP (1:10,000, Abcam Ab98698), goat anti-rabbit polyclonal IgG (H+L) HRP (1:10,000, Invitrogen 65-6120), and goat anti-mouse recombinant polyclonal IgG (H+L) HRP (1:5,000 for claudin-5; 1:10,000 for GAPDH, Invitrogen A28177). All secondary antibodies were diluted in 1% BSA in TBS-T. Membranes were subsequently washed with TBS-T and then visualized using enhanced chemiluminescence reagent (ECL) (Bio-Rad Clarity Western ECL Substrate 170-5060 or Bio-Rad Clarity Max Western ECL Substrate 1705062) on a MicroChemi camera system (FroggaBio). Densitometry was then performed using ImageJ to calculate protein abundance relative to GAPDH.

### PMVEC protein isolation for proteomics analysis

PMVEC from young and aged mice were cultured at a density of 2.5 x 10^5^ cells and grown to confluence on 6-well plates coated with 1% gelatin. Media from the wells were aspirated, and the cells washed three times with sterile PBS. Cells were lysed with 200 *µ*L of ice-cold sterile lysis buffer (2% SDS, 100mM Ammonium bicarbonate, 1mM EDTA, and protease inhibitor cocktail [Sigma-Aldrich P8340]). Cell lysates were then immediately scraped and collected into ice-cold Eppendorf tubes. Samples were sonicated for 10 seconds, put back on ice, then sonicated for another 10 seconds. Centrifugation was then performed at 14,000 x g for 10 minutes at 4℃ and the supernatant transferred to a new chilled tube, which was stored at −80℃. Samples were sent to a core facility at the University of Calgary (Canada) for processing. A BCA assay was used to calculate protein concentrations for each sample, according to the manufacturer’s instructions (ThermoFisher Scientific 23227). Total protein (500 µg) from each sample was removed for analysis. Samples were topped up to 1 mL with lysis buffer, followed by the addition of 500 µL of 6M GuHCl containing dithiothreitol (DTT; 5 mM) and incubation for 1 hour at 37°C to reduce disulphide bonds. Free thiols on cysteines were blocked by adding 15 mM of iodoacetamide for 25 minutes in the dark at room temperature. The thiol blocking reaction was quenched by adding an additional 5mM of DTT.

Samples were labeled with either light (control) or isotopically heavy formaldehyde (40 mM; VWD; Cambridge Isotope Laboratories, Inc. DLM-805-PK). This was followed by addition of sodium cyanoborohydride to a final concentration of 20 mM. The pH was adjusted to 6.5 before incubation overnight at 37°C. The next day, an additional 10 mM of light or heavy formaldehyde was added, followed by 10 mM of sodium cyanoborohydride. Samples were incubated at 37°C for 1 hour. After incubation, samples were transferred into a 50 ml Falcon tube with 50 ml of 8:1 ice-cold acetone methanol. Samples were incubated at −20°C for 4 hours. Samples were centrifuged at 9,000 x g for 15 minutes, then washed in 100% methanol 3 times. Following the last wash, the supernatant was discarded. Samples were air-dried in a fume hood for 5 minutes to remove residual methanol. Samples were then resuspended in 200 µl of 200 µM NaOH, and transferred to a 1.5 ml low-binding protein tube before vortexing for 5 minutes. After the protein pellet was completely resuspended, 300 µl of 200 mM HEPES buffer was added. This was followed by the addition of 50 µg of mass spec-grade trypsin. The pH was adjusted to ∼8 before incubation overnight at 37°C. Samples were then acidified to a pH of less than 3 with trifluoroacetic acid (TFA) and stored at 4°C overnight for SEP PAK (c18) clean up (Waters WAT020515). Conditioning solution (3mL; 90% v/v methanol, 0.1% TFA v/v 10% v/v high performance liquid chromatography [HPLC] water) was passed through the solid phase extraction (SPE), followed by 2 mL of load solution (100% HPLC water, 0.1% v/v TFA). The sample was loaded into the SPE column followed by 1 mL of additional load solution. Samples were desalted by passing 3 mL of desalt solution (5% v/v. methanol, 95% v/v HPLC water, 0.15 TFA) across the SPE, followed by sample collection into a new tube by passing 1 mL of elution solution (50% v/v HPLC water, 50% v/v acetonitrile, 0.1% TFA). Samples were frozen at −80 before submission for LCMS.

### High performance liquid chromatography (HPLC) and mass spectrometry (MS)

As we have described previously, tryptic peptides were analyzed on an Orbitrap Fusion Lumos Tribrid mass spectrometer (Thermo Scientific) operated with Xcalibur (version 4.4.16.14) and coupled to a Thermo Scientific Easy-nLC (nanoflow Liquid Chromatography) 1200 system (43–46). A total mass of 2 μg tryptic peptides were loaded onto a C18 trap (75 um x 2 cm; Acclaim PepMap 100, P/N 164946; ThermoScientific) at a flow rate of 2 µL/min of solvent A (0.1% formic acid in LC-MS grade water). Peptides were eluted using a 120 min gradient from 5 to 40% (5% to 28% in 105 min followed by an increase to 40% B in 15 min) of solvent B (0.1% formic acid in 80% LC-MS grade acetonitrile) at a flow rate of 0.3 μL/min and separated on a C18 analytical column (75 um x 50 cm; PepMap RSLC C18; P/N ES803; ThermoScientific). Peptides were then electrosprayed using 2.1 kV into the ion transfer tube (300°C) of the Orbitrap Lumos operating in positive mode. The Orbitrap first performed a full MS scan at a resolution of 120,000 FWHM to detect the precursor ion having a *m*/*z* between 375 and 1575 and a +2 to +7 charge. The Orbitrap AGC (Auto Gain Control) and the maximum injection time were set at 4e5 and 50 ms, respectively. The Orbitrap was operated using the top speed mode with a 3 second cycle time for precursor selection. The most intense precursor ions presenting a peptidic isotopic profile and having an intensity threshold of at least 5,000 were isolated using the quadrupole and fragmented with HCD (30% collision energy) in the ion routing multipole. The fragment ions (MS^2^) were analyzed in the ion trap at a rapid scan rate. The AGC and the maximum injection time were set at 1e4 and 35 ms, respectively, for the ion trap. Dynamic exclusion was enabled for 45 seconds to avoid acquisition of the same precursor ion having a similar *m/z* (plus or minus 10 ppm).

### Proteomic data and bioinformatics analysis

Spectra data obtained during mass spectrometry were matched to peptide sequences from a mouse FASTA reference file obtained from Uniprot on February 24^th^, 2022, using MaxQuant (v1.6.0.1). MaxQuant settings were set to default except for the following: variable modifications included oxidation (M), acetyl (N-term), deamidation (NQ); label-free quantification was turned on; first search peptide tolerance was set to 10 under the Orbitrap settings; the identifier rule for the FASTA file was set to the Uniprot; maximum peptide mass was set to 6,600 Daltons; minimum peptide length was set to 5 amino acids; match between runs was turned on. Lists containing the identified proteins and peptides were returned and used for subsequent analysis. Gene Ontology pathway analysis was performed using Metascape (47). Only proteins with a Q-value less than 0.05 between groups were included. Analyses were performed with a fold change cut-off of 2, as well as a less stringent cut-off of 1.1.

### PMVEC gene expression analysis

PMVEC from young and aged mice were grown at a density of 2.5 x 10^5^ cells on 6-well plates coated with 1% gelatin. Media from the wells were aspirated, and cells lysed for RNA collection using the RNeasy Mini Kit (Qiagen 74104), as per the manufacturer’s instructions. An on-column DNA digestion was performed during RNA purification using the RNAse-Free DNase set, according to the manufacturer’s protocol (Qiagen 79254). The RNA was reverse transcribed to cDNA using a high-capacity cDNA reverse transcription kit (Applied Biosystems 4368814), as per the manufacturer’s instructions. Quantitative polymerase chain reaction (qPCR) was then performed on the samples using TaqMan Master Mix (ThermoFisher Scientific 4369016) and probes (all from ThermoFisher Scientific) for interleukin (*Il*)*6* (Mm00446190_m1), CXC motif chemokine ligand (*Cxcl*)*1* (Mm04207460_m1), *Il1β* (Mm00434228_m1), C-C motif chemokine ligand (*Ccl*)*2* (Mm00441242_m1), *Cxcl2* (Mm00436450_m1), colony stimulating factor (*Csf*)*2* (Mm01290062_m1), intercellular adhesion molecule-1 (*Icam1*) (Mm00516023_m1), vascular cell adhesion molecule-1 (*Vcam1*) (Mm01320970_m1), E-selectin (*Sele*) (Mm01310197_m1), and hypoxanthine-guanine phosphoribosyltransferase (*Hprt*)*1* (Mm03024075_m1). Subsequent analysis was then performed using the ΔΔCt method, with *Hprt1* used as the housekeeping gene and specific samples normalized to average young PMVEC values.

### Assessment of cell adhesion molecule cell surface abundance

The cell surface abundance of ICAM1, VCAM1, and E-selectin in PMVEC from young and aged mice was assessed using the Guava easyCyte HT Flow Cytometer (EMD Millipore, Billerica, MA). PMVEC were seeded at a density of 2.5 x 10^5^ cells and grown to confluence on 6-well cell culture plates coated with 1% gelatin. Growth media was aspirated from each well and the cells were gently detached using Accutase (Gibco A1110501) for 30 minutes at 37℃. The cells were then centrifuged at 300 x g for 5 minutes at room temperature and resuspended in 3 mL of 0.1% BSA/PBS to a concentration of 500,000 cells/mL. Next, 500 µL of the cell suspension was added into three Eppendorf tubes, along with 1 µL of fluorescently labelled antibodies targeted against ICAM1 (rat monoclonal, 0.5 mg/mL, BioLegend 116112), VCAM1 (rat monoclonal, 0.5 mg/mL, BioLegend 105712), and E-selectin (rat monoclonal, 0.2 mg/mL, BD Biosciences 553751). An additional 500 µL cell aliquot was also taken for an unstained control where no antibody was added. Each tube was incubated with rotation for 1 hour at room temperature, followed by three washes with 0.1% BSA/PBS. Cells were then resuspended in 250 µL of 0.1% BSA/PBS and transferred to a clear-bottom 96-well round bottom plate to run in the Guava easyCyte HT Flow Cytometer. Live cell gates were set based on forward and side scatter, with the first 10,000 events collected and analyzed per sample. Mean fluorescent intensity was calculated for each sample.

### Assessment of PMVEC actin cytoskeleton arrangement

Following growth in 48-well plates coated with 1% gelatin, PMVEC were fixed with 200 µL of 4% paraformaldehyde for 20 minutes, washed three times with PBS, permeabilized with 0.1% Triton X-100 (VWR), and blocked using 3% BSA in PBS. Cells were then incubated in the dark with 200 µL of Phalloidin Tetramethylrhodamine B isothiocyanate (Sigma-Aldrich P1951; Excitation: 540-545nm; Emission: 570-573 nm) at a final concentration of 2 µg/mL in PBS for 1 hour at room temperature. Counterstaining and fluorescent microscopy imaging were carried out as described above (see “Immunocytochemistry of junctional proteins”).

### Statistical Analysis

Data are presented as mean ± standard error of the mean (SEM). Statistical analyses were carried out using the Student unpaired t-test and two-way ANOVA. Results were considered significant at p < 0.05. For the proteomics analysis, the MaxQuant software was used with a peptide FDR of 0.01. Only proteins with a Q-value < 0.05 were used for Metascape analysis.

## Results

### Assessment of the role of aging on PMVEC barrier function under basal conditions

PMVEC isolated from young and aged mice were grown to confluence, and barrier function was assessed by ECIS and the XPerT permeability assay. Following 26 hours after plating, the ECIS data revealed that PMVEC from aged animals exhibited a significantly lower overall resistance compared to PMVEC from young animals (Fig. 1A). In order to directly examine the relationship between junctional disruption and PMVEC permeability, the XPerT permeability assay was employed concomitantly with VE-cadherin staining. PMVEC from young animals exhibited minimal avidin leak staining along with continuous cell-surface VE-cadherin localization (Fig. 1B). In contrast, PMVEC from aged animals exhibited significantly greater avidin staining, which was all co-localized to paracellular regions with discontinuous VE-cadherin staining (Fig. 1B).

**Figure 1:**
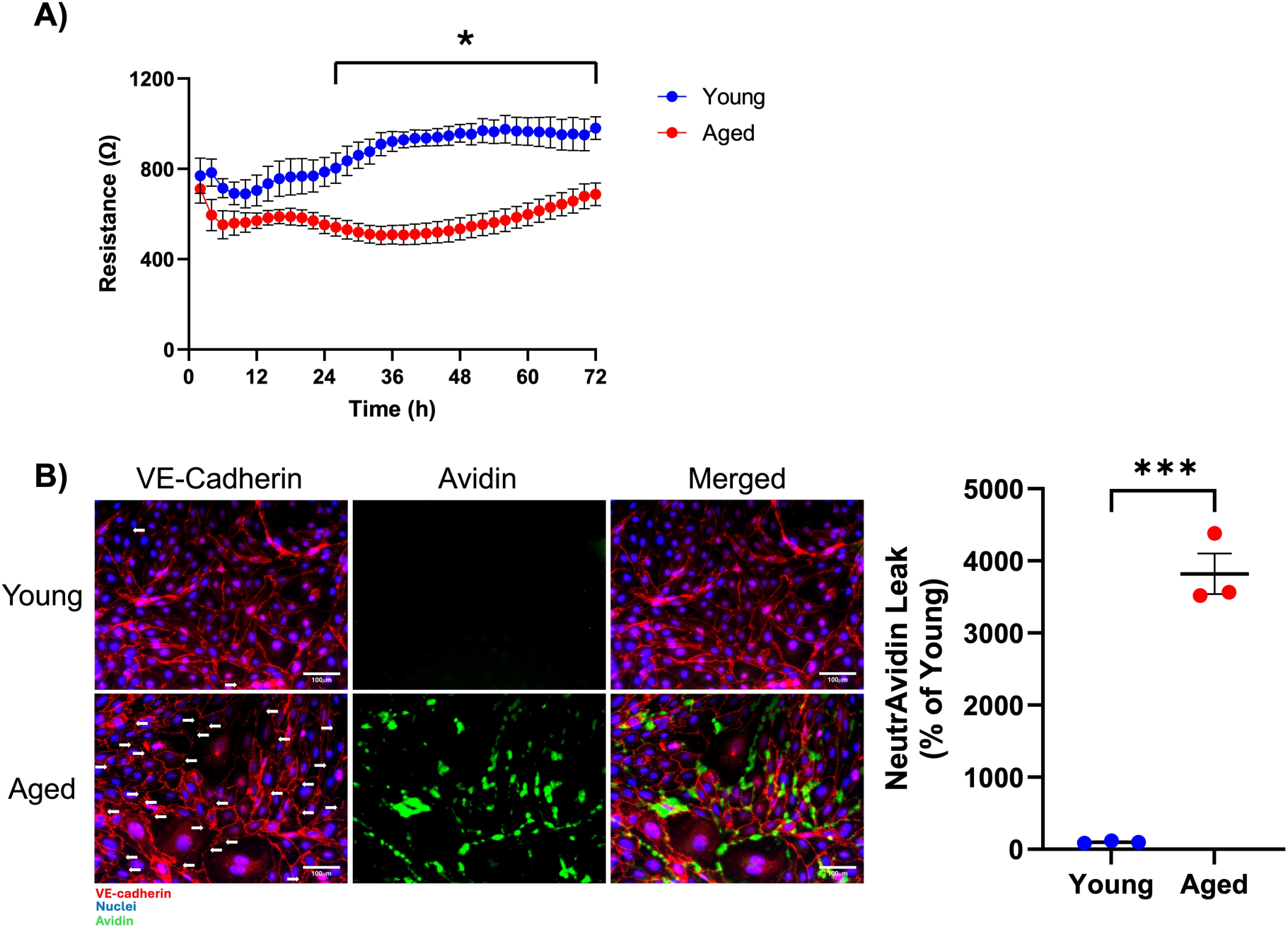
Effect of aging on pulmonary microvascular endothelial cell (PMVEC) permeability under basal conditions. **A)** Monolayer resistance was assessed in PMVEC from young and aged mice by electric cell-substrate impedance sensing (ECIS). Compared to young, PMVEC from aged mice exhibited significantly decreased monolayer resistance 26 hours after seeding. **B)** Immunofluorescent (IF) staining of vascular endothelial (VE)-cadherin (RED) and local leak of NeutrAvidin (GREEN) was carried out in PMVEC monolayers from young and aged animals. PMVEC from young mice exhibited continuous VE-cadherin IF staining around the periphery of the cells, and this was associated with minimal leak. In contrast, PMVEC from aged mice exhibited discontinuous VE-cadherin IF staining (white arrows) and significantly increased avidin leak, with areas of leak colocalized directly at paracellular regions of VE-cadherin discontinuity. For ECIS experiments, n = 4; *p<0.05; Repeated measures two-way ANOVA. For NeutrAvidin Leak experiments, n = 3; ***p<0.001; Unpaired t-test. Scale bar = 100 μm.

### Quantification of junctional proteins in PMVEC from young and aged mice

To continue to explore the impact of age on PMVEC cell-cell junctional proteins, the abundance of the transmembrane proteins, as well as the adherens junction adapter proteins, was assessed using western blot analysis under basal conditions. Compared with young, PMVEC from aged mice exhibited significantly greater VE-cadherin abundance, but lower claudin-5 and *γ*-catenin abundance (Fig. 2). No differences were detected in the other adherens junction adapter proteins, *α*-catenin and *β*-catenin (Fig. 2).

**Figure 2:**
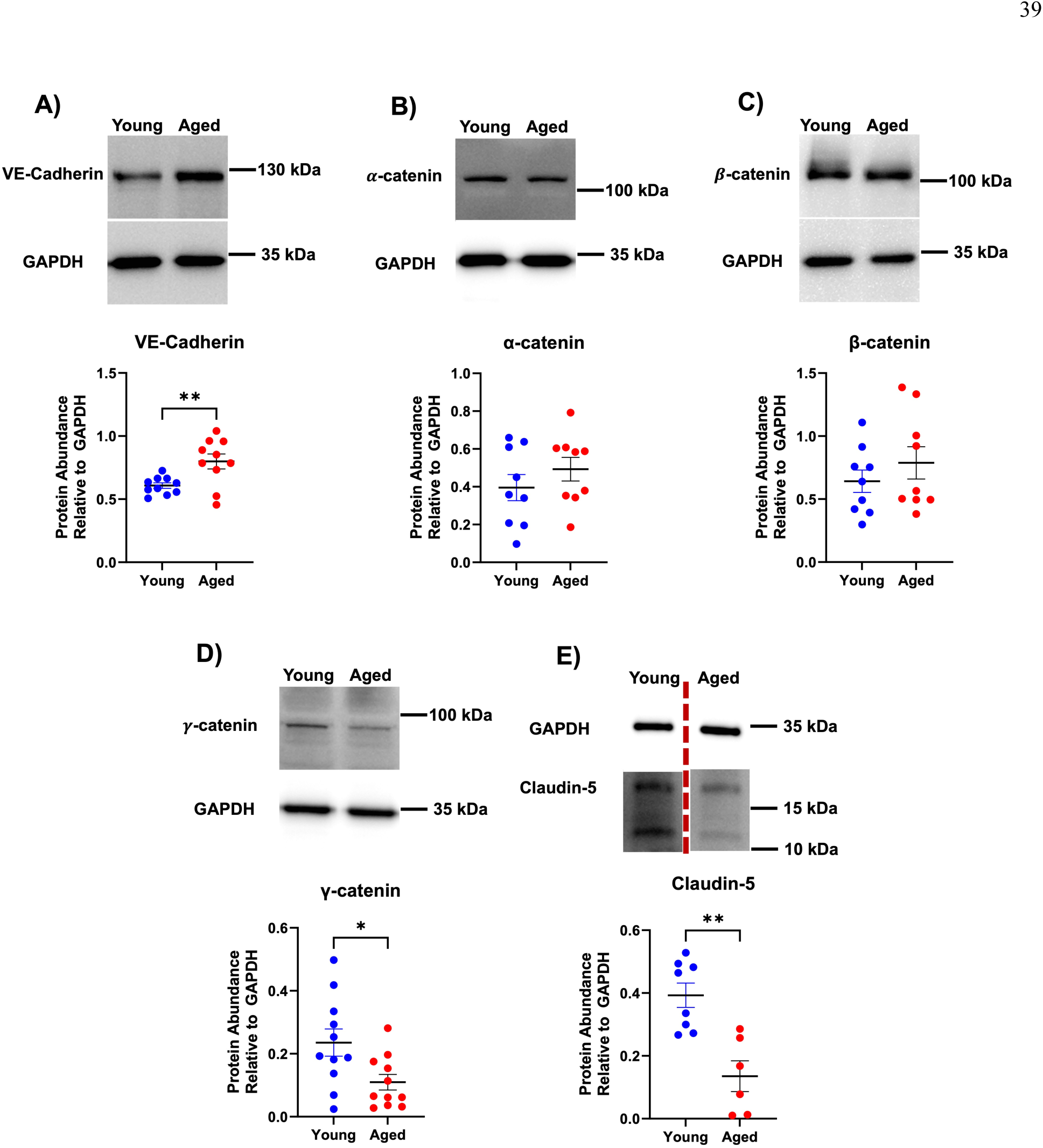
Effect of aging on cell-cell junction protein abundance in pulmonary microvascular endothelial cells (PMVEC). Western blot analyses were performed to quantify the abundance of vascular endothelial (VE)-cadherin (**A**), α-catenin (**B**), β-catenin (**C**), γ-catenin (**D**), and claudin-5 (**E**). Compared with PMVEC from young mice, PMVEC from aged mice exhibited a significant increase in VE-cadherin, but a significant reduction in γ-catenin and claudin-5. n = 6-11; *p<0.05, **p<0.01; Unpaired t-test. The claudin-5 western blot (**E**) has been spliced to remove an experimental group that was not pertinent to the comparison presented.

### Proteomics analysis of PMVEC from young and aged mice

Shotgun proteomics analysis was employed to examine global changes in the proteome between PMVEC isolated from young and aged mice (Fig. 3A). Using a fold change cut-off of 2, a total of 29 proteins were enriched in PMVEC from young animals, while 17 proteins were enriched in the aged (Supplementary Table 1). STRING-db analysis (48) revealed an enrichment of collagen-containing extracellular matrix and vesicle pathways in the young PMVEC (Fig. 3B), and extrinsic component of plasma membrane, leukocyte transendothelial migration, and fibroblast pathways in the aged PMVEC (Fig. 3C). Gene ontology analysis revealed many pathways that were significantly differentially enriched in both groups; these included pathways such as endocytosis, regulation of PI3K/protein kinase B signal transduction, regulation of smooth muscle cell proliferation, cell surface interactions at the vascular wall, and cell activation pathways associated with inflammation, such as neutrophil degranulation (Fig. 3D). A common feature of each of these pathways is actin cytoskeleton modification or rearrangement (49–54). Additional analysis was conducted on the proteomics dataset, with a less stringent fold change cut-off of 10% variance, in order to extend the search to other proteins that may be relevant to the above pathways. Examination of the significantly differentially enriched proteins revealed 218 proteins enriched in young and 114 proteins enriched in aged; this led to pathways associated with the actin cytoskeleton that appeared in both groups, driven by different proteins enriched in each group (Supplementary Fig. 1). Specifically, these pathways were driven by 23 proteins from the young PMVEC, such as beta actin, actin related protein 2/3 complex subunit 3, and actin related protein 2 (Supplementary Table 2), and 28 proteins from the aged PMVEC, including actinin alpha 4, destrin, and adducin 1 (Supplementary Table 3).

**Figure 3:**
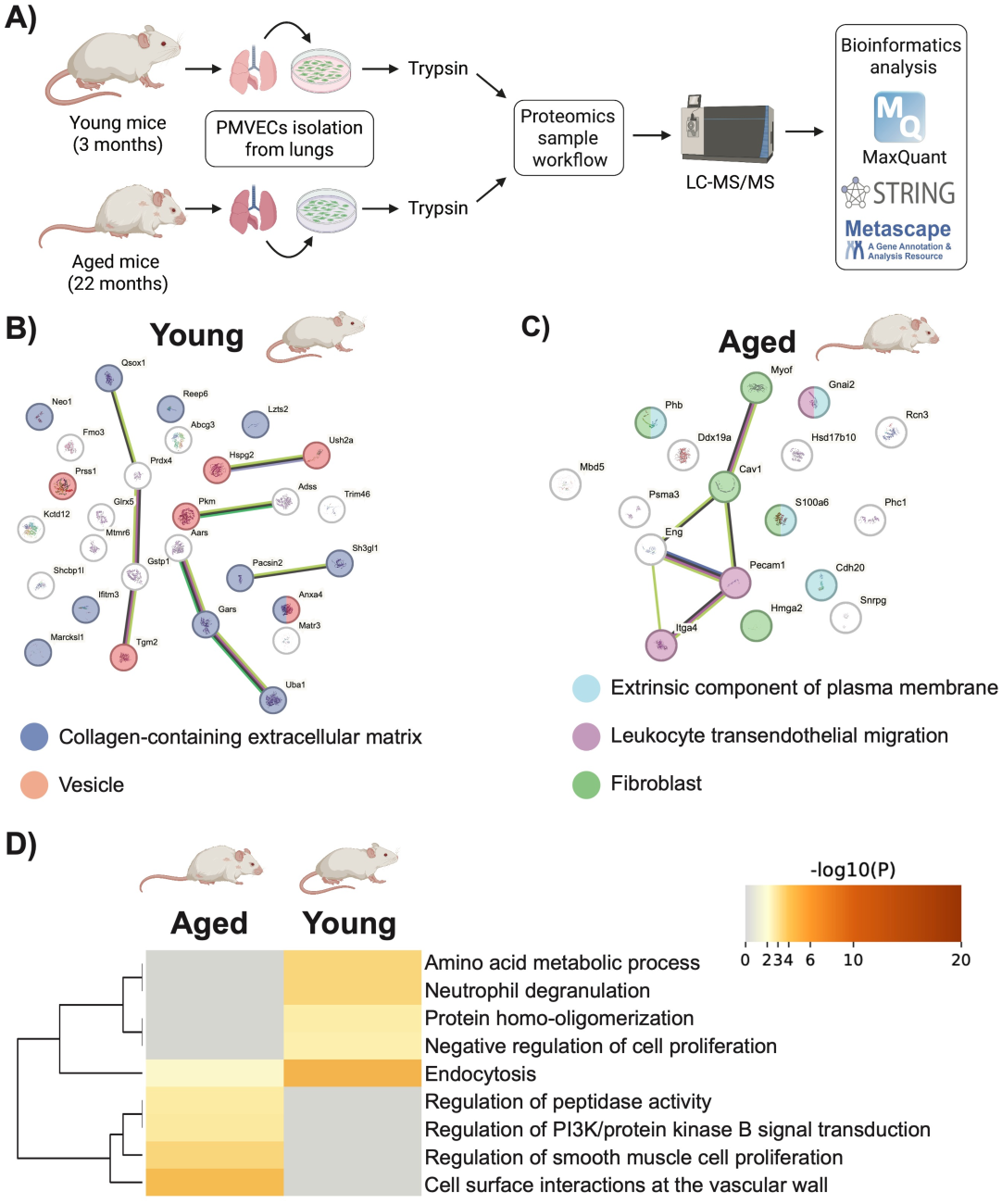
Proteomics analysis of pulmonary microvascular endothelial cells (PMVEC) from young and aged mice. **A)** Workflow schematic of proteomics experimental design. PMVEC were isolated from young and aged mice, cultured to confluency *in vitro*, and lysed for protein collection. Proteins were digested with trypsin, labeled with light or heavy formaldehyde, subjected to LC-MS/MS, and analyzed using MaxQuant. **B&C)** STRING-db analysis (48) of enriched proteins in PMVEC from young and aged mice. Enrichment of collagen-containing extracellular matrix (DARK BLUE) and vesicle (ORANGE) in young, and extrinsic component of plasma membrane (LIGHT BLUE), leukocyte transendothelial migration (PURPLE) and fibroblasts (GREEN) in aged, are shown in colored spheres. **D)** Metascape analysis (47) revealed a number of biological pathways that were altered between PMVEC from young and aged animals, including endocytosis, regulation of peptidase activity, regulation of PI3K/protein kinase B signal transduction, and cell surface interactions at the vascular wall. Schematic graphics used in this figure were created using BioRender. Datasets were derived from n = 4 per age group.

### Assessment of actin cytoskeleton arrangement in PMVEC from young and aged mice

To follow up on the pathways identified in the proteomics analysis associated with alterations in the actin cytoskeleton (Fig. 3 & Supplemental Fig. 1), further analysis was conducted to assess the arrangement of the actin cytoskeleton in PMVEC from young and aged mice (Fig. 4). This analysis revealed minimal stress fiber formation with strong cortical actin staining in the PMVEC from young animals. In contrast, PMVEC from aged animals exhibited stress fiber formation under basal conditions (Fig. 4).

**Figure 4:**
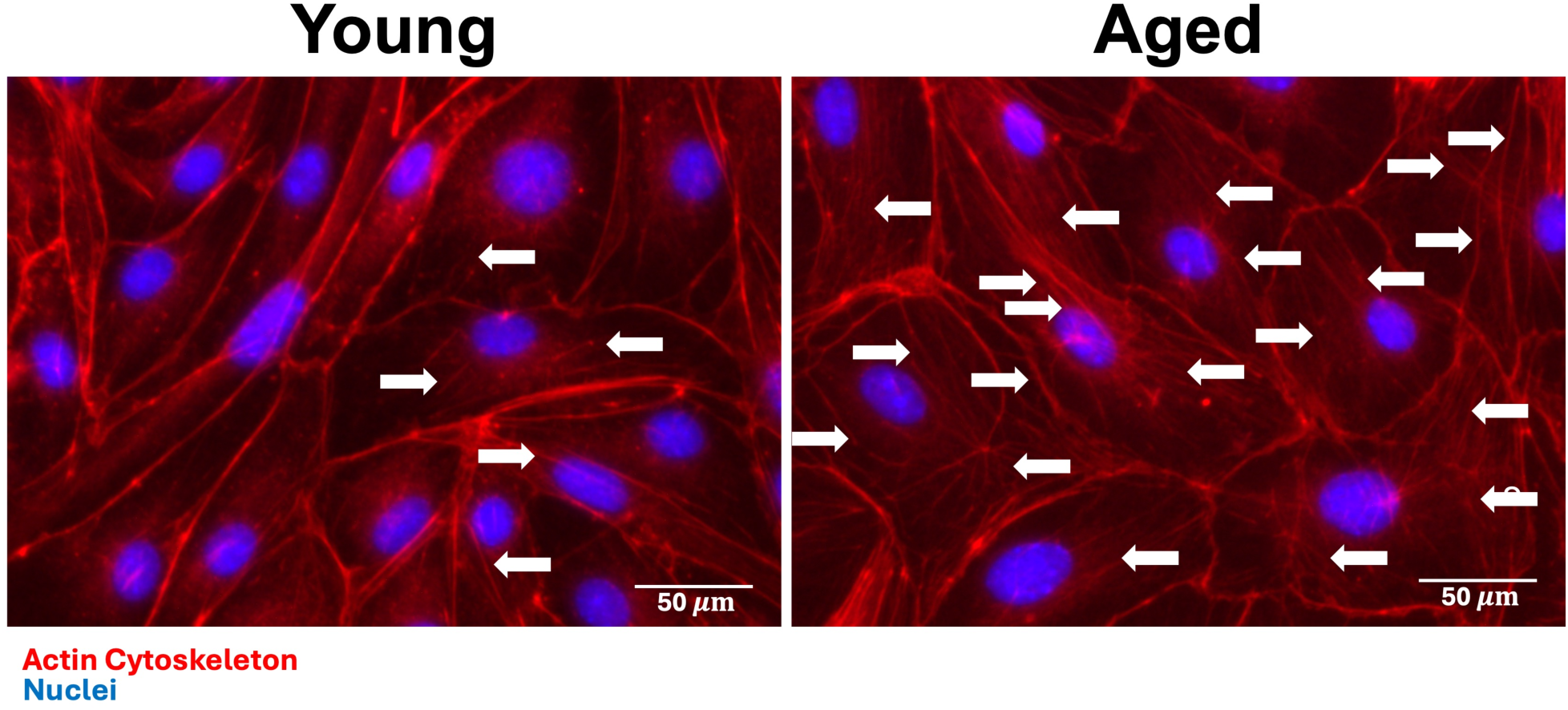
Effect of aging on actin cytoskeleton organization in pulmonary microvascular endothelial cells (PMVEC). Phalloidin staining was performed to examine the actin cytoskeleton in PMVEC from young and aged mice. Compared to PMVEC from young animals, which exhibit primarily cortical actin staining, PMVEC from aged animals have stress fiber formation under basal conditions. Above are representative images from n = 3 experiments. Arrows indicate actin stress fibers. Scale bar = 50 μm.

### Impact of aging on inflammatory mediator expression in PMVEC from young and aged mice

To follow up on other key findings from the proteomics analysis associated with inflammatory signaling, assessment of inflammation was conducted in the PMVEC. To examine the potential impact of age on the expression of pro-inflammatory mediators under basal conditions, qPCR analysis was employed to quantify expression of a variety of cytokines and chemokines, including *Il6*, *Cxcl1*, *Il1β*, *Ccl2*, *Cxcl2*, and *Csf2*, in PMVEC isolated from young and aged mice. Across all cytokines, no significant differences were observed between young and aged PMVEC under basal conditions (Fig. 5).

**Figure 5:**
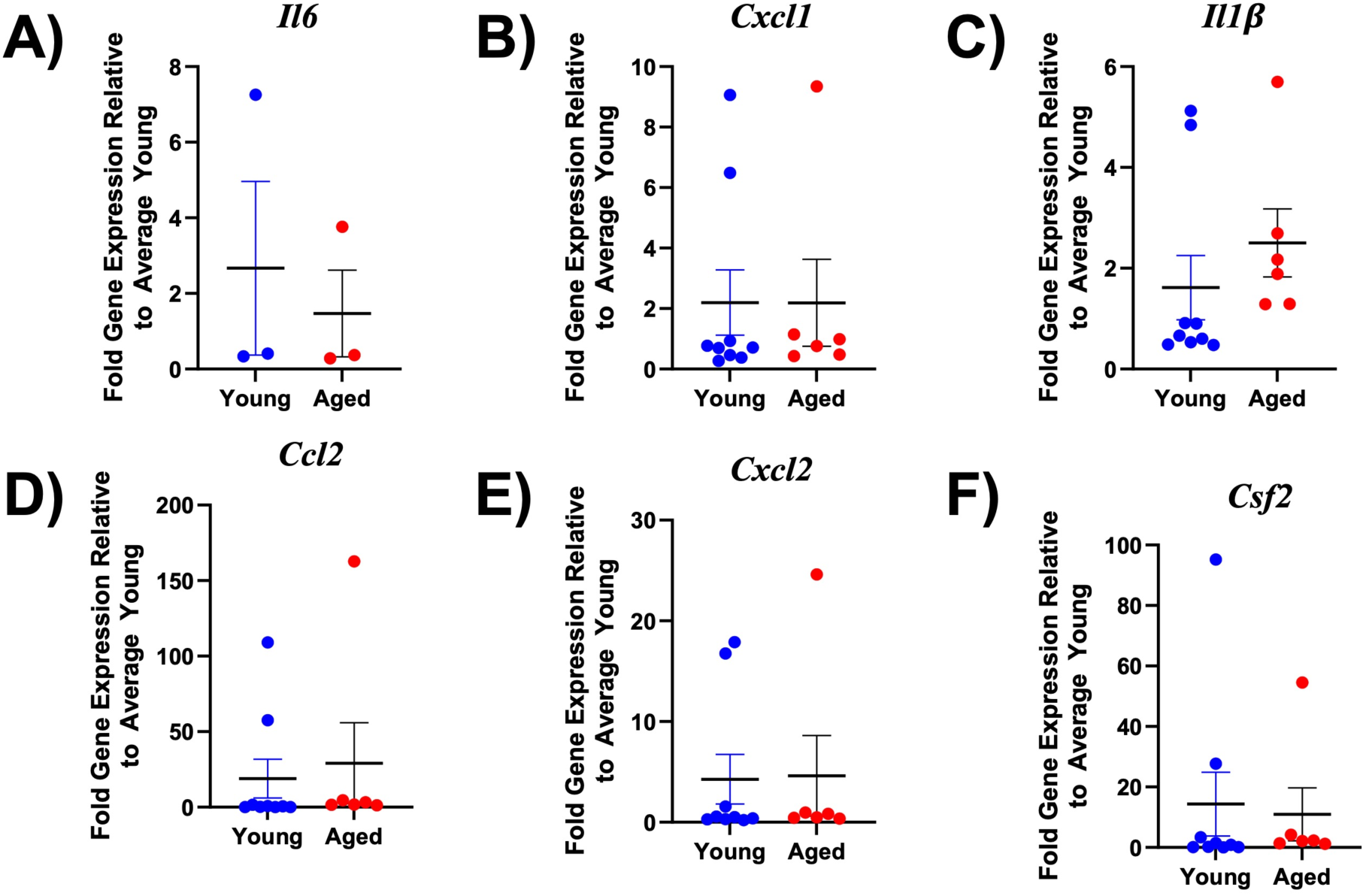
Analysis of proinflammatory cytokine expression in pulmonary microvascular endothelial cells (PMVEC) from young and aged animals under basal conditions. Expression levels of interleukin (*Il*)*6* (**A**), CXC motif chemokine ligand (*Cxcl*)*1* (**B**), Il1β (**C**), C-C motif chemokine ligand (*Ccl*)*2* (**D**), *Cxcl2* (**E**), and colony stimulating factor (*Csf*)*2* (**F**) were assessed by qPCR. No significant differences were observed between young and aged PMVEC across all cytokines. n = 3-9; Unpaired t-test.

### Effect of aging on PMVEC cell adhesion molecules

Additional assessment of PMVEC activation was performed through analysis of cell adhesion molecule expression and cell surface abundance. qPCR analysis revealed significantly decreased expression of *Icam1*, *Vcam1*, and *Sele* in PMVEC from aged mice compared to young (Fig. 6A). While cell surface presence of ICAM1 and VCAM1 was detected with flow cytometry under basal conditions in both young and aged PMVEC, there was no discernable presence of E-selectin, as indicated by the overlap of the basal signal with the unstained control cells (Fig. 6B). Further, analysis of mean fluorescence intensity revealed that cell surface abundance of all the adhesion molecules was not significantly different between young and aged; notably, there was a trend towards a decrease in ICAM1 and VCAM1 (Fig. 6B).

**Figure 6:**
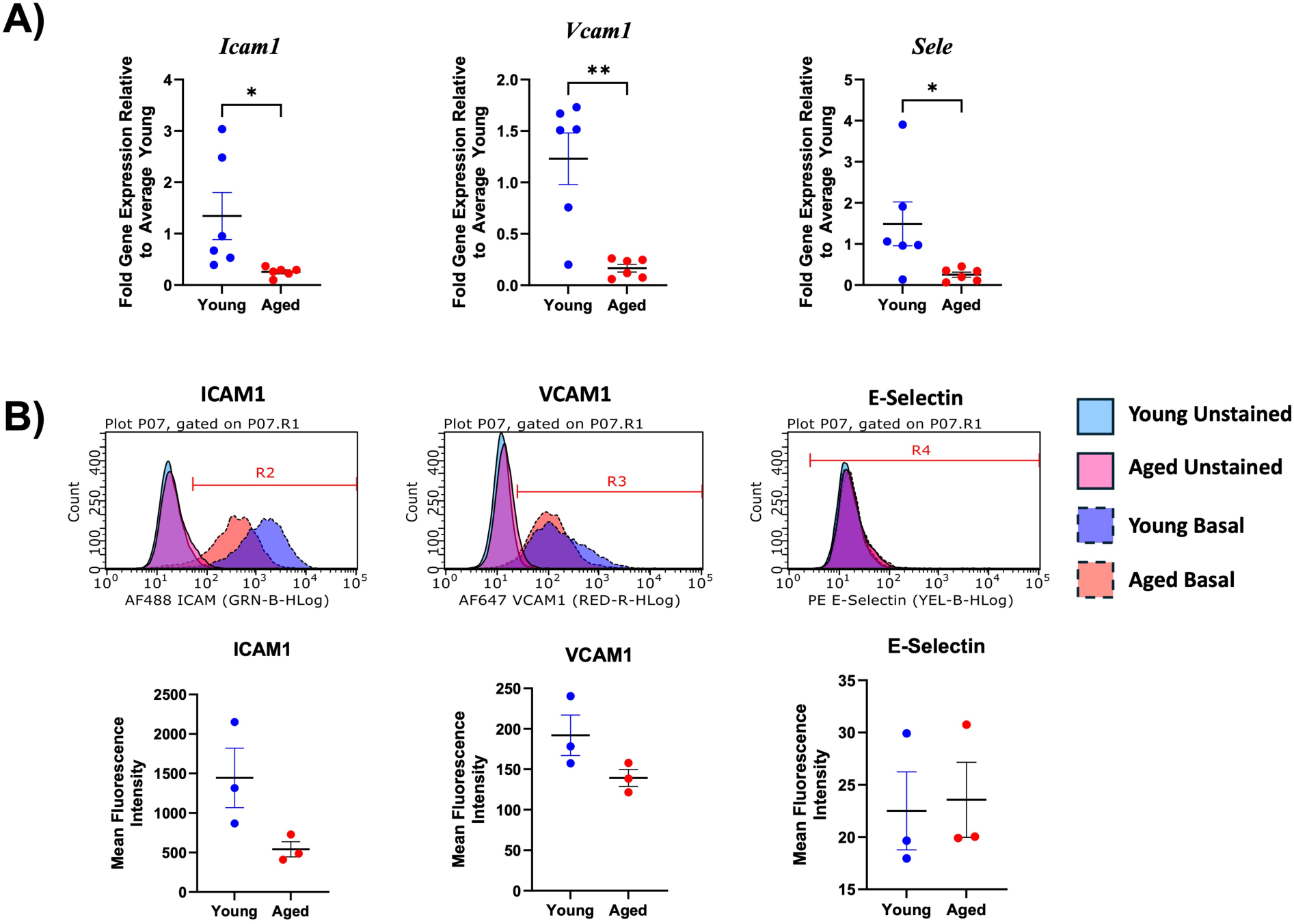
Assessment of cell adhesion molecule expression and cell surface abundance in pulmonary microvascular endothelial cells (PMVEC) from young and aged mice. Intercellular adhesion molecule-1 (*Icam1*/ICAM1), vascular cell adhesion molecule 1 (*Vcam1*/VCAM1), and E-selectin (*Sele*) expression and cell surface abundance were assessed by qPCR and flow cytometry, respectively. Relative to young, PMVEC from aged mice exhibited significantly decreased *Icam1*, *Vcam1*, and *Sele* expression (**A**). While there was a trend towards decreased ICAM1 and VCAM1, no statistically significant differences were observed in cell surface abundance mean fluorescence intensity for the three cell adhesion molecules (**B**). For qPCR experiments, n = 6; *p<0.05, **p<0.01; for flow cytometry experiments, n = 3; Unpaired t-test.

### Assessment of the role of aging on PMVEC barrier function under inflammatory conditions

To explore whether the greater basal leak in aged PMVEC led to augmented PMVEC barrier dysfunction under pathological conditions, PMVEC from young and aged mice were treated with PBS (control) or cytomix (pro-inflammatory), and VE-cadherin staining and local leak were assessed. While not significant, an increase in permeability in the PMVEC from young animals was observed at 4 hours following cytomix treatment, which persisted at 24 hours (Fig. 7A). Furthermore, the increase in leak was associated with disrupted and discontinuous VE-cadherin cell-surface localization (Fig. 7A). In aged PMVEC, treatment with cytomix led to a significant increase in avidin leak compared to baseline (Fig. 7A). Moreover, compared to young, the PMVEC dysfunction in response to cytomix, including VE-cadherin disruption and local leak, were significantly increased in the PMVEC isolated from the aged animals (Fig. 7A). Assessment of tight junctions was also performed by immunocytochemical analysis of claudin-5. Similar to VE-cadherin, PMVEC from young mice exhibited continuous claudin-5 around the periphery of the cells under basal conditions, with localization appearing to become disrupted as early as 4 hours post-cytomix stimulation (Fig. 7B). This disruption persisted and was associated with an apparent loss of claudin-5 localization in young PMVEC at 24 hours post-cytomix stimulation (Fig. 7B). As we have shown previously, PMVEC from the aged animals appeared to have sparse claudin-5 localization around the periphery of the cells under control conditions, which persisted following cytomix stimulation (Fig. 7B) (20).

**Figure 7:**
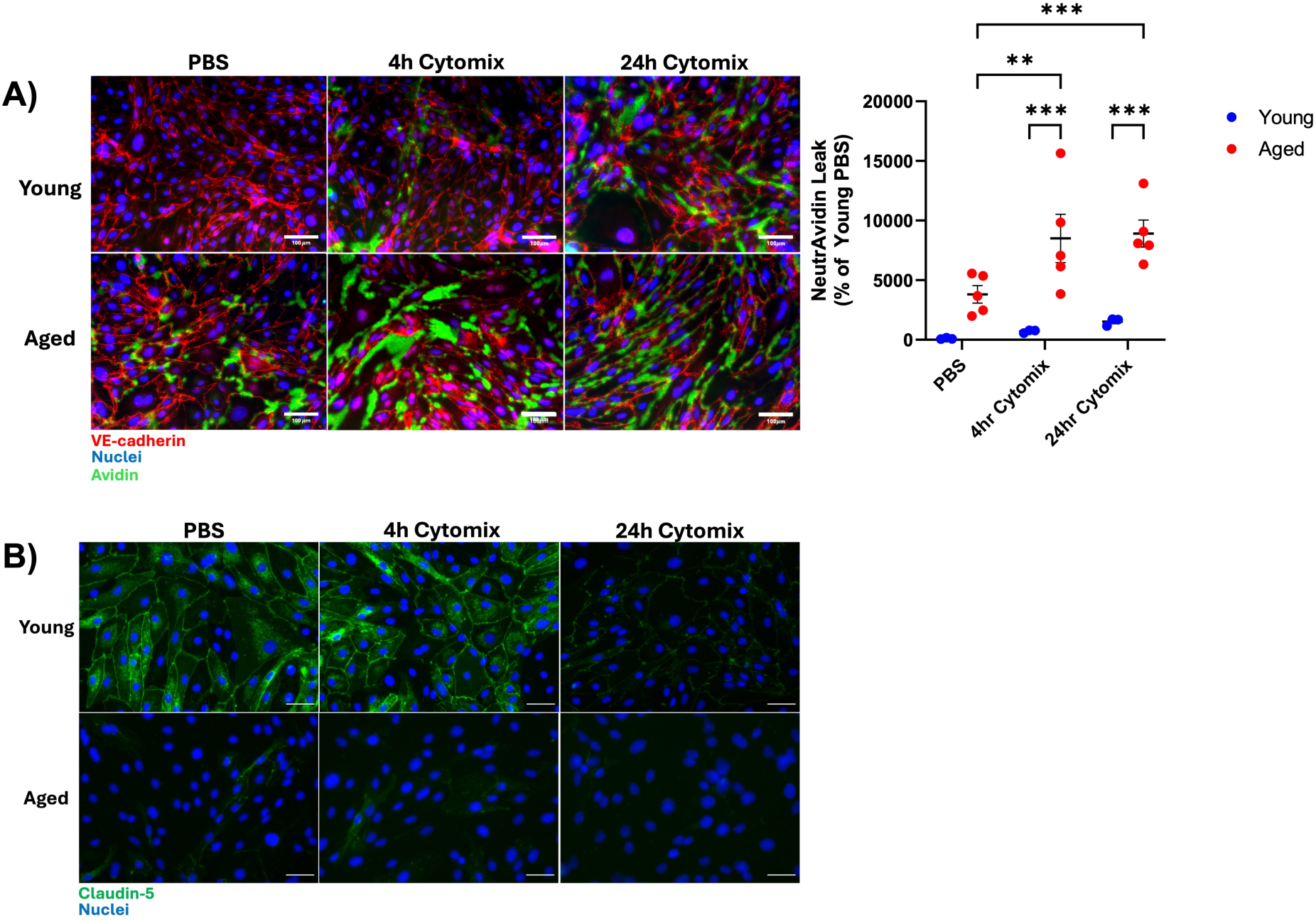
Effect of aging on localized leak and cell-cell adherens and tight junctions in pulmonary microvascular endothelial cells (PMVEC) under inflammatory conditions. **A)** Immunofluorescent (IF) staining of vascular endothelial (VE)-cadherin (RED) and local leak of NeutrAvidin (GREEN) was assessed in PMVEC from young and aged mice stimulated with PBS and cytomix. Compared to PBS, PMVEC from young animals treated with cytomix exhibited disruption of VE-cadherin and a trend toward increased avidin leak. PMVEC from aged animals had increased leak and VE-cadherin disruption under basal conditions, which was exacerbated, and significantly increased compared to young, following cytomix stimulation. **B)** IF staining of claudin-5 (GREEN) revealed strong and continuous cell-surface localization in PMVEC from young mice under basal conditions, with disrupted and decreased staining following cytomix stimulation, particularly at 24 hours. PMVEC from aged animals exhibited sparse claudin-5 cell-surface localization under control conditions, which persisted following cytomix stimulation. Above are representative images from n = 3-5 experiments. For NeutrAvidin Leak experiments, n = 3-5; **p<0.01, ***p<0.001; Two-way ANOVA. Scale bar = 100 μm.

### Effect of aging on actin cytoskeleton arrangement in PMVEC under inflammatory conditions

Given the differences observed in the actin cytoskeleton arrangement in young and aged PMVEC under basal conditions, subsequent analysis was conducted under inflammatory conditions. Similar to what we have shown previously, stress fiber formation began to occur 4 hours post-cytomix stimulation in the PMVEC from young animals, with a further increase being observed after 24 hours (Fig. 8) (55). PMVEC from the aged animals exhibited stress fiber formation under basal conditions, which was further augmented following stimulation with cytomix at both the 4- and 24-hour time points (Fig. 8).

**Figure 8:**
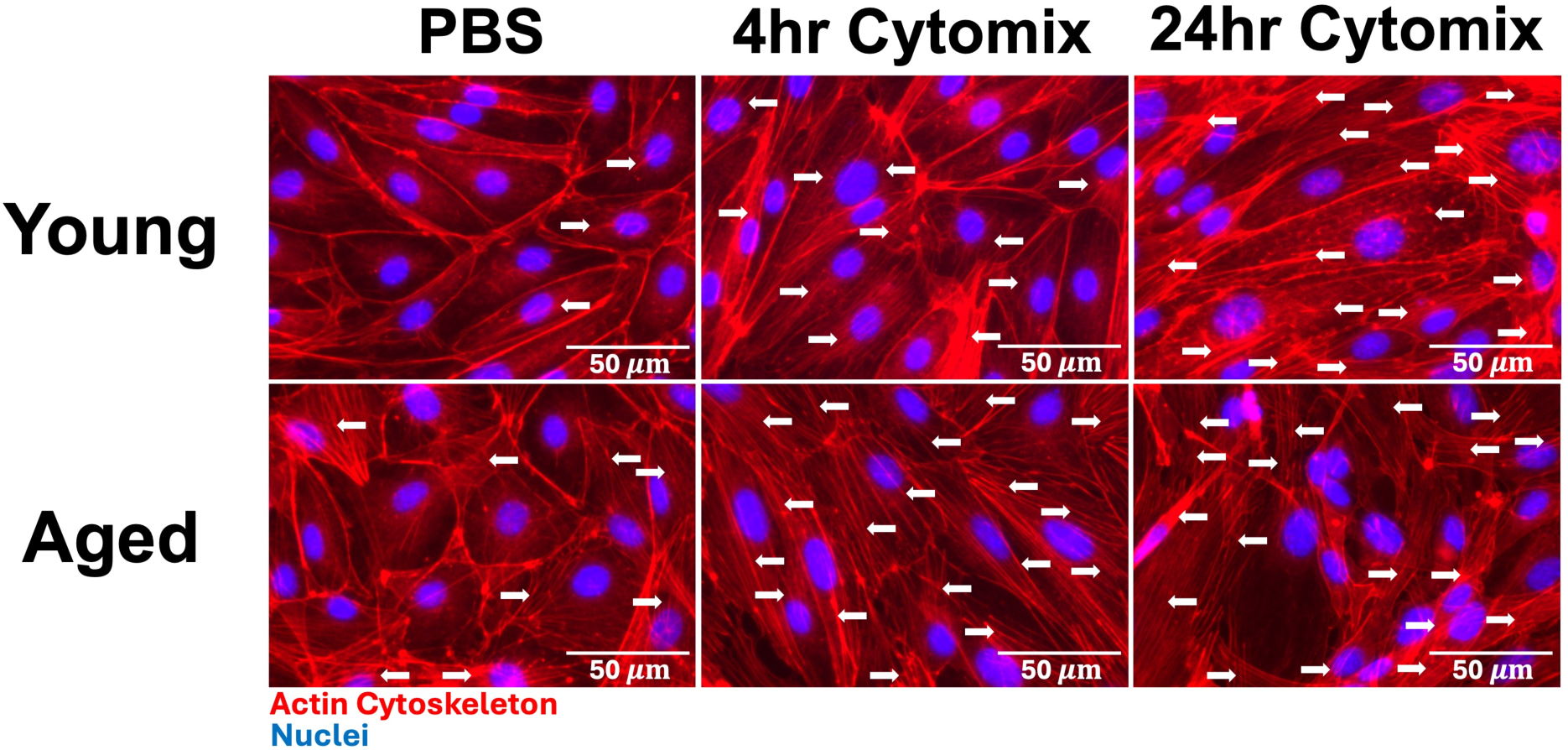
Actin cytoskeleton organization (phalloidin staining) in pulmonary microvascular endothelial cells (PMVEC) from young and aged mice under inflammatory conditions. PMVEC from young animals exhibit increased stress fiber formation beginning at 4 hours post-cytomix stimulation (RED), which persists at 24 hours. In comparison, PMVEC from aged animals exhibit stress fibers under basal conditions with augmented stress fiber formation at both 4 and 24 hours post-cytomix stimulation. Above are representative images from n = 3 experiments. Arrows indicate actin stress fibers. Scale bar = 50 μm.

## Discussion

The objective of this study was to investigate the impact of aging on PMVEC barrier function, both under basal and inflammatory conditions. To assess this, we conducted studies in isolated PMVEC from mice at 2-3 months of age and at 18-22 months of age, which correspond to approximately 20- and 65-years of age in humans, respectively (56). Overall, our findings supported our general hypothesis that aging was associated with PMVEC barrier dysfunction due to impairments in cell-cell junction integrity. Furthermore, we propose that the deficiencies in PMVEC barrier function with age may be mediated by actin cytoskeletal alterations, particularly the presence of stress fibers. To our knowledge, this is the first *in vitro* study to demonstrate impaired endothelial barrier function of the pulmonary microvasculature directly with age due to disrupted cell-cell junctions. Our findings may begin to elucidate the underlying mechanisms associated with pulmonary microvascular injury, including augmented permeability, which underpin the worse clinical outcomes with age during lung injury.

The primary outcome of this study was the observation of an increase in permeability in PMVEC monolayers isolated from the aged mice compared to young under basal and inflammatory conditions, and its direct association with cell-cell junctional disruption. The use of the XPerT macromolecule permeability assay in the current study, which enables the visualization of localized leak concomitantly with immunocytochemical analysis of junctional proteins, demonstrated that the increased leak with age was directly localized to paracellular regions of VE-cadherin discontinuity. Furthermore, following treatment with cytomix, the disrupted VE-cadherin and leak were further exacerbated. Additionally, immunocytochemical analysis of claudin-5 revealed minimal staining in the PMVEC from aged mice, which persisted following cytomix stimulation. Based on these findings, we conclude that the age-associated impairment in cell-cell junction integrity was directly contributing to the elevated microvascular permeability.

Other studies in the literature have observed similar findings with aging being associated with endothelial barrier dysfunction and impairments in cell junctions. A number of *in vivo* animal studies assessing blood-brain barrier permeability following injury demonstrated augmented microvascular permeability, along with disruptions in the cell-cell junctions with age (57–59). Furthermore, additional *in vivo* studies examining endothelial barrier function in aged rodents under healthy conditions have demonstrated increased vascular permeability within the blood-brain barrier, blood-retinal barrier, and thoracic aorta, concomitant with reduced junctional protein abundance or disrupted and reduced junctional protein localization (60–67). *In vitro* studies have also found similar outcomes in primary brain microvascular endothelial cells, human umbilical vein endothelial cells, human aortic endothelial cells, and human umbilical cord blood-derived endothelial cells (21, 22, 24–26, 28). Specific alterations were observed with VE-cadherin, claudin-5, occludin, zona occludens (ZO)-1, and ZO-2, including disrupted immunofluorescent staining or fragmentation of the proteins (21, 25, 26), decreased protein abundance (21, 22, 24), and decreased protein localization at the cell junctions (22, 25, 28). A more recent study using drug-induced senescence in primary human lung microvascular endothelial cells demonstrated reduced claudin-5, ZO-1, and VE-cadherin protein abundance and localization at the junctions (27). Notably, these previous *in vitro* studies that have examined endothelial barrier function with age have relied on the induction of senescence to model aging, which may not accurately recapitulate the microenvironment observed in elderly individuals (21, 22, 24–28). Our current work expands upon these previous studies through the use of primary endothelial cells from the microvasculature of the lungs, as well as the fact that these cells were isolated directly from young and aged mice. Furthermore, our *in vitro* data support the notion that the barrier dysfunction observed with age is intrinsic to the endothelial cells.

While the aforementioned discussion offers insights into the general behavior of aged PMVEC with respect to permeability, our subsequent analysis of inflammation, junctional protein abundance, and proteomics provides some molecular insight into the various proteins and pathways involved in this altered response. Inflammation is well-established as a contributor to endothelial cell-cell junction damage, thereby inducing endothelial barrier dysfunction (68–72). Furthermore, healthy aging is associated with elevated markers of inflammation, including increased cytokine expression and abundance, and increased activation of inflammatory cells; this increase in basal inflammatory signaling with age has been well characterized and is referred to as ‘inflammaging’ (73–77). However, examining inflammatory signaling in PMVEC, through quantification of inflammatory mediator expression and cell surface abundance of endothelial cell adhesion molecules, revealed no significant differences between young and aged PMVEC. In fact, there was a trend towards significantly lower cell-surface abundance of certain adhesion molecules, like ICAM1 and VCAM1. With regard to the junctional proteins, surprisingly, we observed a significant increase in VE-cadherin abundance with age, but a significant reduction in claudin-5 and *γ*-catenin. The inclusion of claudin-5 and *γ*-catenin within cell-cell junctions is thought to occur in mature endothelial junctions (78–81). Based on this, we speculate that the PMVEC from aged mice have deficiencies in forming mature cell-cell junctions, and to compensate for this deficit, there is an increase in VE-cadherin abundance. There is evidence to support this; for instance, VE-cadherin adhesion is known to be the primary adhesion event during vessel development, and VE-cadherin adhesion is known to promote activation of claudin-5 expression (78, 82). The disrupted VE-cadherin localization at the cell membrane observed in the PMVEC from aged mice may explain the subsequent reduction in proteins like claudin-5 and *γ*-catenin, as these latter proteins rely on the preliminary, early formation of the adherens junction. Collectively, we speculate that with age, PMVEC exhibit deficiencies in the formation of mature and stable junctions, and that elevated inflammatory signaling is not driving the barrier dysfunction observed.

Further interrogation of our proteomics analysis revealed several biological pathways that were altered in the PMVEC, including pathways associated with the actin cytoskeleton. Consistent with this observation, our recent publication, assessing the transcriptomic differences in pulmonary microvascular cells from young and aged mice using single-cell RNA sequencing (20), found this same pathway was also differentially enriched. The actin cytoskeleton is known to play a pivotal role in modulating endothelial barrier function (7, 9–11, 83). The presence of cortical actin is known to enhance barrier function through the generation of intracellular tension, which enhances clustering, assembly, development, and integrity of cell-cell junctions (9–11, 83, 84). Conversely, actin depolymerization and the formation of actin stress fibers lead to reduced junctional binding within the membrane, ultimately resulting in intercellular gap formation and endothelial barrier dysfunction; importantly, this has been suggested to occur during ARDS (3, 4, 7, 8, 85–87). In the present study, PMVEC from young animals exhibited cortical actin staining with minimal stress fiber formation under basal conditions. Cytomix stimulation induced stress fiber formation, which was consistent with previous work conducted by our group (55). Interestingly, PMVEC from the aged animals exhibited actin stress fiber formation under basal conditions, which was subsequently augmented following stimulation with cytomix. This aligns with the literature, which suggests that aging and senescence are associated with altered actin cytoskeleton protein expression, distribution, and protein-protein interactions, ultimately leading to changes in cytoskeletal dynamics and actin disorganization (27, 88–92). While further research is warranted, the presence of actin stress fibers may provide molecular insights into the loss of cell-cell junctional integrity and subsequent barrier dysfunction observed in the aged PMVEC. It also provides a putative mechanism for future studies to further pursue and explore as a potential therapeutic target to mitigate age-related vascular damage and elevated morbidity during lung injury.

Speculating on the clinical implications of our data in humans, we postulate that the endothelial barrier dysfunction observed with age may contribute to augmented morbidity during lung injury. We have previously demonstrated in a mouse model of mechanical ventilation that aging was associated with augmented vascular permeability (20). The work from the current study provides insight into differences in the ability of PMVEC from aged mice to reform a barrier once this barrier is disrupted, which is what is required during the formation of an intact monolayer under basal conditions in a cell culture model. These observed alterations may predispose the cells to greater injury following exposure to an additional insult. Specifically, we suggest that the impaired PMVEC cell-cell junctions and weakened barrier function with age lead to an inherent predisposition to pulmonary vascular damage, including following a secondary insult such as sepsis. Furthermore, we suggest that the deficiencies in vascular barrier function could underpin the elevated morbidity and mortality in older patients suffering from lung injury, such as during ARDS.

Our study has some limitations. First, the PMVEC isolated from young and aged mice were all male, which disregards the role of biological sex in mediating barrier function. Second, our *in vitro* model utilized PMVEC cultured alone, in the absence of other cell types. In particular, the presence of immune cells and/or immune cell factors has been shown to modulate PMVEC barrier function, particularly under inflammatory conditions (93). Lastly, our *in vitro* model relied on static conditions, which are not representative of *in vivo* vascular flow conditions. Previous studies have indicated that flow promotes endothelial barrier function (94). Despite these limitations, the findings from the current study support our previous work assessing the role of aging on PMVEC barrier function and cell-cell junctions in an animal model of mechanical ventilation. Specifically, aging was shown to promote PMVEC barrier dysfunction, which was associated with transcriptional changes to cell-cell junction proteins, as determined by single-cell RNA-sequencing (20).

Future studies will aim to address the above limitations. This will include isolating PMVEC from both male and female animals, employing co-culture systems with other cell types such as neutrophils, developing a system that facilitates the flow of growth media across the luminal side of the PMVEC, and using plasma from patients diagnosed with ARDS as a more pathologically relevant injurious/inflammatory stimulus. Additionally, future studies will delve deeper into the role of aging on the actin cytoskeleton and its modulation of PMVEC barrier function, including investigating the efficacy of known treatments or inhibitors of actin stress fiber formation to elucidate the effects on cell-cell junction disruption and barrier dysfunction.

In summary, this study aimed to assess the effect of aging on PMVEC barrier function, under both basal and inflammatory conditions. Our overall conclusion is that aging was associated with heightened PMVEC barrier dysfunction due to impaired cell-cell junctions associated with augmented actin cytoskeleton disorganization. These findings offer insights into microvascular-specific mechanisms underlying the increase in age-related susceptibility during lung injury and give possible avenues, including those associated with actin cytoskeleton disorganization, for future research to target and evaluate for therapeutic potential.

## Data Availability

The proteomics datasets are available in the PRIDE repository (project accession: PXD065409). All additional data can be made available from the corresponding author upon reasonable request.

## Supplemental Material

Supplemental Figs. S1 XXXXXXXX

Supplemental Tables S1-S3: XXXXXXXX

## Acknowledgments

The authors would like to thank Dr. Gediminas Cepinskas, Dr. Marat Slessarev, Dr. Thomas Drysdale, and both past and present members of the Lung Lab for their assistance and suggestions for study design and writing.

## Author Contributions

A.M., L.W., C.P., P.P., S.Y., E.P., and A.D., performed experiments; A.M., C.P., P.P., S.Y., A.D., D.Y., and S.E.G., analyzed data; A.M., P.P., S.Y., R.A.W.V., A.D., and S.E.G. interpreted results of experiments; A.M., and A.D. prepared figures; A.M., R.A.W.V., A.D., and S.E.G. drafted the manuscript; A.M., A.D., R.A.W.V., and S.E.G., edited and revised the manuscript; A.M., L.W., C.P., S.M., P.P., S.Y., E.P., R.A.W.V., A.D., and S.E.G., approved the final version of the manuscript; A.M., S.M., and S.E.G., conceived and designed the research.

