## Supplementary Figure 1 for "The role of aging on endothelial cell-cell junctions and pulmonary microvascular permeability"

**Supplemental Material**

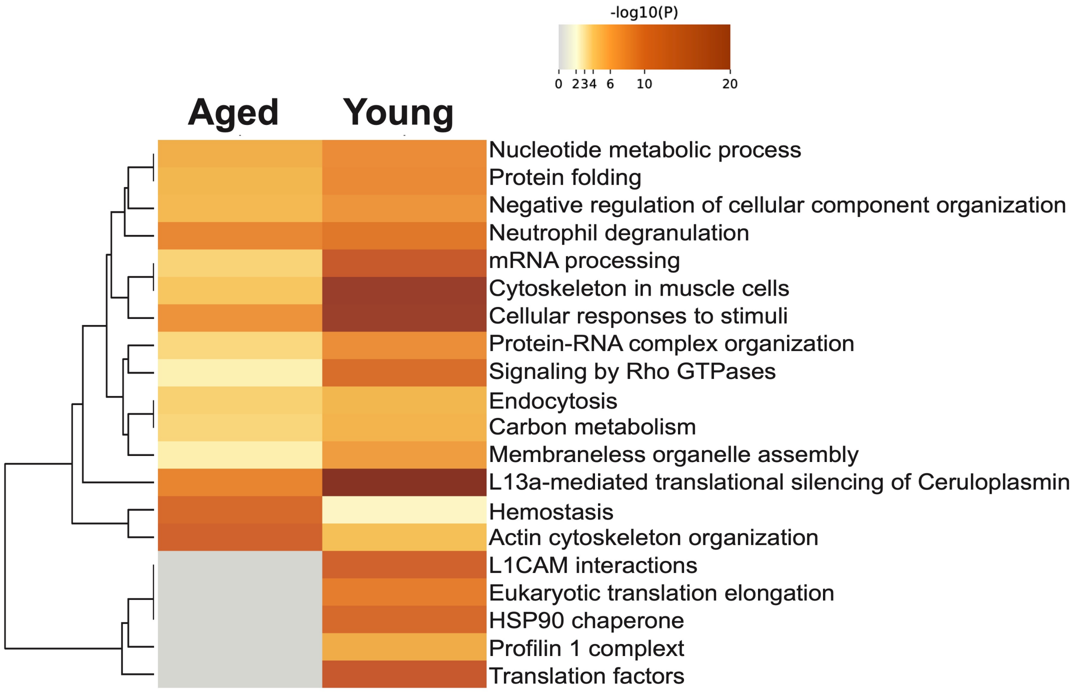


**Supplementary Figure 1:** Metascape analysis of the effect of age on the proteome of pulmonary microvascular endothelial cells (PMVEC). Accumulative hypergeometric p-values and enrichment factors were calculated and used for filtering as performed as a two-sided analysis. Remaining significant terms were then hierarchically clustered into a tree based on Kappa-statistical similarities among their gene's memberships. Then, 0.3 kappa score was applied as the threshold to cast the tree into term clusters. The analysis revealed a number of biological pathways that were altered between PMVEC from young and aged mice, including mRNA processing and translation, HSP90 chaperone signaling, neutrophil degranulation, and actin cytoskeleton organization.
