## Supplementary Table 1 for "The role of aging on endothelial cell-cell junctions and pulmonary microvascular permeability"

**Supplementary Table 1 |** Shotgun proteomics from aged 22 months versus young 3 months pulmonary microvascular endothelial cells (PMVECs).

Peptide sequences were identified from the mouse UniProt protein database with appended standard laboratory and common contamination protein entries and reverse decoy sequences using the Andromeda algorithm as implemented in the MaxQuant software package v1.6.0.1, using a peptide FDR of 0.01. Search parameters included a mass tolerance of 1 ppm for the parent ion and 0.5 Da for the fragment ions, carbamidomethylation of cysteine residues (+57.021464 Da), variable N-terminal modification by acetylation (+42.010565). Database searches were not constrained by enzyme specificity, but limited to a maximal length of 25 residues. The table shows unique identified peptides as listed in the "modificationSpecificPeptides" output file of MaxQuant. Peptide sequences matching reverse or contaminant entries were removed. Mass: peptide mass; charges: precursor charge states of associated MS/MS spectra; PEP: Andromeda Posterior Error Probability.

| Proteins | Gene Names | Protein Names | Potential Contaminant | Ratio H/L normalized | Intensity 1 (L) young | Intensity 1 (H) aged | Intensity 2 (L) young | Intensity 2 (H) aged | Intensity 3 (L) young | Intensity 3 (H) aged |
|---|---|---|---|---|---|---|---|---|---|---|
| Q792Z0;Q792Y8;Q9Z1R9;Q9 | rss3;Gm10334;Prss1;Try4;Try5;Try10;Prss | Anionic trypsin-2 | | 0.042526 | 23874000 | 490820 | 45539000 | 923290 | 18143000 | 784520 |
| A0A494BB51;A0A494BB76;C | Lzts2 | Leucine zipper putative tumor suppressor 2 | | 0.047953 | 33112000 | 2951800 | 55266000 | 15154000 | 18041000 | 577700 |
| O08807;B1AZS9 | Prdx4 | Peroxiredoxin-4 | | 0.0786 | 5191300 | 195100 | 3947800 | 272630 | 13729000 | 649060 |
| Q99P81;D3Z0Z4 | Abcg3 | ATP-binding cassette sub-family G member 3 | | 0.10113 | 0 | 0 | 14587000 | 154500 | 980100 | 549040 |
| Q3TC52;D3YXA6;Q7TNM2;E9 | Trim46 | Tripartite motif-containing protein 46 | | 0.11612 | 0 | 0 | 120310000 | 1780700 | 762700 | 556910 |
| CON__P00761;Q9CPN9 | | | + | 0.12621 | 79782000 | 3006100 | 185140000 | 8294100 | 221470000 | 12431000 |
| Q2QI47;Q2QI47-3 | Ush2A | Usherin | | 0.12912 | 4897400 | 218870 | 7487100 | 401800 | 0 | 0 |
| J3QQ47;Q3TTP0 | Shcbp1l | SHC SH2 domain-binding protein 1-like protein | | 0.15895 | 7300800 | 365420 | 13391000 | 1261700 | 0 | 0 |
| Q9WVE8;A0A2R8W6S4;A0A | Pacsin2 | rotein kinase C and casein kinase substrate in neurons protein 2 | | 0.1677 | 3970100 | 243950 | 7825200 | 807430 | 7117400 | 1007700 |
| A0A1Y7VN70;Q80Y14 | Glrx5 | Glutaredoxin-related protein 5, mitochondrial | | 0.18569 | 2072500 | 290690 | 6780900 | 721540 | 0 | 0 |
| P28667 | Marcksl1 | MARCKS-related protein | | 0.19681 | 0 | 0 | 0 | 0 | 3059800 | 1083500 |
| Q8VE11;Q8VE11-2 | Mtmr6 | Myotubularin-related protein 6 | | 0.21453 | 17044000 | 1486500 | 0 | 0 | 0 | 0 |
| D3YTR2;D3YTR1 | Reep6 | | | 0.26168 | 2036100 | 31748 | 0 | 0 | 23446000 | 23879000 |
| Q8BND5;Q8BND5-3;Q8BND5 | Qsox1 | Sulfhydryl oxidase 1 | | 0.26272 | 1379100 | 146780 | 2067000 | 304190 | 0 | 0 |
| Q9CQW9 | Ifitm3 | Interferon-induced transmembrane protein 3 | | 0.31897 | 0 | 0 | 1602900 | 3428100 | 11335000 | 2582500 |
| Q80UG5-3;A2A6U3;Q80UG5 | 09-Sep | Septin-9 | | 0.41688 | 2238400 | 25701 | 10273000 | 285220 | 922240 | 318800 |
| P97501 | Fmo3 | Dimethylaniline monooxygenase [N-oxide-forming] 3 | | 0.41946 | 1799300 | 345390 | 3944000 | 1118500 | 0 | 0 |
| Q8K310;A0A494BAZ2;A0A49 | Matr3 | Matrin-3 | | 0.42156 | 0 | 0 | 3502700 | 1395700 | 4369200 | 563270 |
| Q9CZD3 | Gars | Glycine--tRNA ligase | | 0.42385 | 428670 | 141470 | 1182500 | 519440 | 13415000 | 0 |
| A0A0N4SW89;P97429;S4R1F | Anxa4 | Annexin A4;Annexin | | 0.42487 | 2653900 | 994510 | 13890000 | 6332000 | 5214600 | 2853400 |
| P52480;P52480-2;A0A1L1SU | Pkm | Pyruvate kinase PKM | | 0.4255 | 3019000 | 228390 | 12158000 | 10075000 | 91883000 | 8918300 |
| P19157;A0A494B908;A0A494 | Gstp1;Gstp2 | Glutathione S-transferase P 1;Glutathione S-transferase P 2 | | 0.42842 | 36836000 | 8650500 | 79428000 | 29980000 | 24571000 | 18494000 |
| Q02053 | Uba1 | Ubiquitin-like modifier-activating enzyme 1 | | 0.4295 | 10384000 | 6154200 | 0 | 0 | 0 | 0 |
| Q7TQG5;E9QK04;P97798-5;F | Neo1 | Neogenin | | 0.4328 | 0 | 0 | 2766100 | 2069000 | 2504100 | 493940 |
| E9PZ16;B1B0C7;Q05793;A0A | Hspg2 | ane-specific heparan sulfate proteoglycan core protein;Endorepellin;LG3 peptide | | 0.43692 | 22501000 | 2452100 | 24550000 | 80473000 | 63186000 | 14574000 |
| P46664 | Adss | Adenylosuccinate synthetase isozyme 2 | | 0.46702 | 0 | 0 | 2688800 | 1577800 | 2898200 | 932500 |
| Q8BGQ7 | Aars | Alanine--tRNA ligase, cytoplasmic | | 0.47423 | 2332100 | 509110 | 3970200 | 1614200 | 0 | 0 |
| P21981;G3UXE8;REV__P024 | Tgm2 | Protein-glutamine gamma-glutamyltransferase 2 | | 0.48483 | 14032000 | 1667900 | 11016000 | 1930900 | 4844900 | 3137200 |
| A0A0R4J2B2;Q6WVG3 | Kctd12 | BTB/POZ domain-containing protein KCTD12 | | 0.49302 | 3774300 | 952520 | 8924600 | 2565500 | 5836500 | 1623100 |
| Q62419;A0A3B2W7K0;Q8BX | Sh3gl1;Sh3gl2 | Endophilin-A2;Endophilin-A1 | | 0.49739 | 2878900 | 712130 | 3509900 | 1414100 | 0 | 0 |
| P45376 | Akr1b1 | Aldose reductase | | 0.51941 | 2172400 | 568020 | 12190000 | 6087200 | 3255600 | 2206500 |
| F8WJK8;Q99L47;E9Q1X9;E9C | St13 | Hsc70-interacting protein | | 0.53073 | 3167100 | 718600 | 9050800 | 4976700 | 5672900 | 3562400 |
| G5E850;P56395;E0CY88 | Cyb5a | Cytochrome b5 | | 0.55808 | 5483600 | 1749900 | 14951000 | 5055000 | 7412700 | 4443900 |
| A0A494BAX5;A0A498WGK2; | Nars | Asparagine--tRNA ligase, cytoplasmic | | 0.57715 | 1382700 | 853020 | 3359600 | 2078400 | 0 | 0 |
| P26231;A0A494BAD0;E0CXB | Ctnna1;Ctnna2 | Catenin alpha-1;Catenin alpha-2 | | 0.5861 | 2086700 | 1995000 | 0 | 0 | 0 | 0 |
| E9PYH3;Q8BWU8 | Etnppl | Ethanolamine-phosphate phospho-lyase | | 0.59448 | 450300 | 422810 | 12211000 | 1472500 | 806590 | 651590 |

| | | | | | | | | | |
|---|---|---|---|---|---|---|---|---|---|
| B7ZNU9;Q61235 | Sntb2 | Beta-2-syntrophin | 0.59949 | 333310 | 131340 | 886700 | 478050 | 701950 | 688520 |
| Q9D1A2;A0A494B9S3;A0A49 | Cndp2 | Cytosolic non-specific dipeptidase | 0.602 | 5949800 | 2464200 | 7134900 | 3063900 | 12249000 | 7644000 |
| P51150;A0A0N4SVR6;A0A0N | Rab7a | Ras-related protein Rab-7a | 0.60278 | 2553300 | 895440 | 8671000 | 4813900 | 1959400 | 1181600 |
| A0A3B2WCD8;Q3U0V1;A0A3 | Khsrp | Far upstream element-binding protein 2 | 0.60827 | 0 | 0 | 1704200 | 1858300 | 2390100 | 960510 |
| Q9CQ75 | Ndufa2 | ADH dehydrogenase [ubiquinone] 1 alpha subcomplex subunit 2 | 0.6088 | 0 | 0 | 1272100 | 1145000 | 2056700 | 601700 |
| P26040 | Ezr | Ezrin | 0.60979 | 0 | 0 | 2022900 | 1057600 | 1654500 | 1103900 |
| Q9DCN2-2;Q9DCN2;F2Z456 | Cyb5r3 | 5 reductase 3 membrane-bound form;NADH-cytochrome b5 reductase 3 soluble form; | 0.61686 | 3558900 | 1717200 | 6148100 | 3125800 | 2125000 | 1538300 |
| A0A286YDT5;G5E8Q8 | Adgrf5 | | 0.61923 | 1514200 | 396720 | 3480300 | 2046300 | 0 | 0 |
| B1AWE0;Q6PFA2;B1AWE1;B | Clta | Clathrin light chain A | 0.62182 | 812080 | 396720 | 3549000 | 2082500 | 0 | 0 |
| H7BXC3;P17751;P17751-2 | Tpi1 | Triosephosphate isomerase | 0.62963 | 12011000 | 3960700 | 26878000 | 13025000 | 14867000 | 11111000 |
| P60335 | Pcbp1 | Poly(rC)-binding protein 1 | 0.63189 | 2304100 | 815640 | 5156100 | 2889400 | 0 | 0 |
| A0A494B9X6;A0A494BAB1;C | Gsto1 | Glutathione S-transferase omega-1 | 0.63693 | 2253300 | 573690 | 0 | 0 | 2820300 | 1140900 |
| O55125 | Nipsnap1 | Protein NipSnap homolog 1 | 0.64508 | 231370 | 157520 | 600570 | 301880 | 657560 | 433920 |
| P97807-2;P97807 | Fh | Fumarate hydratase, mitochondrial | 0.6455 | 345950 | 210520 | 1014200 | 630810 | 1257600 | 979190 |
| P14733 | Lmnb1 | Lamin-B1 | 0.65432 | 2020000 | 943890 | 5623800 | 5959200 | 961100 | 450830 |
| Q6X632 | Gpr75 | Probable G-protein coupled receptor 75 | 0.65604 | 1180000 | 698440 | 2152400 | 1921500 | 2476700 | 1126400 |
| Q9JKB3-2;Q9JKB3 | Ybx3 | Y-box-binding protein 3 | 0.65886 | 0 | 0 | 5254100 | 2461400 | 0 | 0 |
| H3BKM0;Q9DBG3;Q9DBG3-2 | Ap2b1 | AP-2 complex subunit beta | 0.66575 | 0 | 0 | 519300 | 310220 | 814280 | 259890 |
| A0A498WGD8;Q8CDN6;A0A4 | Txnl1 | Thioredoxin-like protein 1 | 0.66686 | 467920 | 188810 | 2769100 | 3046900 | 3669100 | 4930700 |
| Q9Z2X1;Q9Z2X1-2;J3QMT0;J | Hnrnpf | ibonucleoprotein F;Heterogeneous nuclear ribonucleoprotein F, N-terminally processe | 0.66919 | 2985700 | 2115700 | 27926000 | 18326000 | 17685000 | 11519000 |
| Q8BGD9;B2RWE8 | Eif4b | Eukaryotic translation initiation factor 4B | 0.67027 | 21485000 | 6100000 | 24741000 | 12162000 | 13539000 | 22444000 |
| P50516;P50516-2;D3YWH3;D | Atp6v1a | V-type proton ATPase catalytic subunit A | 0.67028 | 680190 | 349570 | 4458000 | 2161000 | 3188900 | 2645100 |
| F6WEU2;A0A338P6M1;A0A3 | Gm7873 | | 0.67324 | 3550600 | 1226400 | 8150700 | 4151000 | 5525200 | 3505600 |
| E9Q3X0;Q9EQK5;D3Z2N7 | Mvp | Major vault protein | 0.67559 | 1237300 | 551400 | 3431300 | 1882500 | 1211700 | 1000800 |
| P05202 | Got2 | Aspartate aminotransferase, mitochondrial | 0.67609 | 3324500 | 2071600 | 18352000 | 3214900 | 0 | 0 |
| A0A494B987;Q3U1J4 | Ddb1 | DNA damage-binding protein 1 | 0.68232 | 1112600 | 546920 | 817310 | 668690 | 1129200 | 788510 |
| Q7TMM9 | Tubb2a | Tubulin beta-2A chain | 0.68469 | 0 | 0 | 9589700 | 4373800 | 5197500 | 6410500 |
| P40142;A0A286YE28 | Tkt | Transketolase | 0.68559 | 5202300 | 2105300 | 20209000 | 11645000 | 14652000 | 8394400 |
| Q62167;P16381;Q3V086;Q61 | Ddx3x;D1Pas1 | endent RNA helicase DDX3X;Putative ATP-dependent RNA helicase Pl10 | 0.6857 | 2016900 | 642910 | 33432000 | 22327000 | 40036000 | 36964000 |
| Q6ZWX6 | Eif2s1 | Eukaryotic translation initiation factor 2 subunit 1 | 0.69041 | 0 | 0 | 1858000 | 1605000 | 3813000 | 2493900 |
| H7BWZ3;Q9JM76;D3Z2F7;D3 | Arpc3 | Actin-related protein 2/3 complex subunit 3 | 0.70231 | 973560 | 455260 | 0 | 0 | 1678700 | 916170 |
| P38647 | Hspa9 | Stress-70 protein, mitochondrial | 0.71157 | 29605000 | 11200000 | 43685000 | 29601000 | 43694000 | 26465000 |
| P14901 | Hmox1 | Heme oxygenase 1 | 0.71532 | 3968700 | 2140400 | 4687200 | 2721700 | 2520900 | 1413200 |
| V9GXV0;A0A668KL51;Q91VA | Idh3b | Isocitrate dehydrogenase [NAD] subunit, mitochondrial | 0.71723 | 390650 | 260770 | 2917700 | 1104900 | 0 | 0 |
| Q8R1Q8 | Dync1li1 | Cytoplasmic dynein 1 light intermediate chain 1 | 0.71759 | 1183200 | 626190 | 2375500 | 1874300 | 1793900 | 1404300 |
| Q9CWJ9 | Atic | protein PURH;Phosphoribosylaminoimidazolecarboxamide formyltransferase;IMP cyclo | 0.72115 | 5269200 | 1980000 | 7814100 | 3761900 | 1827400 | 1472500 |
| Q543K9;P23492;A0A2I3BQH2 | Pnp;Pnp2 | Purine nucleoside phosphorylase | 0.7251 | 0 | 0 | 1548900 | 1104300 | 2342400 | 863970 |
| P62305 | Snrpe | Small nuclear ribonucleoprotein E | 0.72749 | 412380 | 244670 | 1117900 | 1347300 | 1796200 | 1135600 |
| B1AXW5;B1AXW6;P35700;B | Prdx1 | Peroxiredoxin-1 | 0.72897 | 11724000 | 4474800 | 31003000 | 16389000 | 81336000 | 40899000 |
| O35350 | Capn1 | Calpain-1 catalytic subunit | 0.72959 | 763140 | 298990 | 1637800 | 898080 | 0 | 0 |
| H7BX95;Q6PDM2;Q6PDM2-3 | Srsf1 | Serine/arginine-rich splicing factor 1 | 0.72969 | 0 | 0 | 1989200 | 1019700 | 12402000 | 1618600 |
| A0A5F8MPB9;P26043;Q7TSC | Rdx | Radixin | 0.73156 | 402580 | 145500 | 15501000 | 7152300 | 8628300 | 7604500 |
| M0QWU8;P31786;Q4VWZ5 | Dbi | Acyl-CoA-binding protein | 0.73303 | 9307500 | 2993900 | 15398000 | 8591100 | 0 | 0 |
| A0A0R4J0Z1;P08003 | Pdia4 | Protein disulfide-isomerase A4 | 0.7439 | 0 | 0 | 9894500 | 5341500 | 2982800 | 2146900 |
| P10126;D3Z3I8;D3YZ68;P626 | Eef1a1 | Elongation factor 1-alpha 1 | 0.74439 | 45544000 | 18538000 | 100610000 | 64233000 | 10234000 | 7250100 |

| | | | | | | | | | |
|---|---|---|---|---|---|---|---|---|---|
| P62814;Q91YH6;A0A0U1RNU | Atp6v1b2 | V-type proton ATPase subunit B, brain isoform | 0.7453 | 5177800 | 2137400 | 4059500 | 1685400 | 0 | 0 |
| A0A0N4SV66;Q8CGP4;C0HKE | aa;Hist1h2ah;H2afj;Hist1h2ak;Hist1h2af;H | type 1-H;Histone H2A.J;Histone H2A type 1-K;Histone H2A type 1-F;Histone H2A type | 0.75093 | 45764000 | 16478000 | 117360000 | 72568000 | 144340000 | 89978000 |
| Q3TWW8 | Srsf6 | Serine/arginine-rich splicing factor 6 | 0.75189 | 0 | 0 | 0 | 0 | 6726200 | 5558900 |
| A2AW05;Q08943;Q08943-2 | Ssrp1 | FACT complex subunit SSRP1 | 0.75618 | 0 | 0 | 622880 | 501050 | 1218800 | 760560 |
| P20152;A0A0A6YWC8;A2AKJ | Vim | Vimentin | 0.75621 | 538490000 | 195470000 | 1035100000 | 553590000 | 1009100000 | 750250000 |
| F8WI35;A0A1W2P768;P8424 | H3f3a;Hist1h3b;Hist1h3a;H3f3c | istone H3;Histone H3.3;Histone H3.2;Histone H3.1;Histone H3.3C | 0.75629 | 232680000 | 79844000 | 533330000 | 334640000 | 278270000 | 133620000 |
| Q7TPV4 | Mybbp1a | Myb-binding protein 1A | 0.7573 | 0 | 0 | 425500 | 380120 | 1144200 | 458380 |
| P57784 | Snrpa1 | U2 small nuclear ribonucleoprotein A | 0.75822 | 759050 | 443720 | 1612300 | 1547800 | 1608900 | 775110 |
| P35550;A0A140LIR6;Q80WS | Fbl;Fbll1 | transferase fibrillarin;rRNA/tRNA 2-O-methyltransferase fibrillarin-like protein 1 | 0.75832 | 0 | 0 | 2616000 | 1852400 | 2607400 | 1904300 |
| Q5SXR6;Q68FD5;F6Z1R4 | Cltc | Clathrin heavy chain;Clathrin heavy chain 1 | 0.75928 | 11002000 | 4074000 | 59773000 | 22829000 | 10583000 | 6817400 |
| P42125 | Eci1 | Enoyl-CoA delta isomerase 1, mitochondrial | 0.75974 | 0 | 0 | 2986200 | 1444200 | 1486100 | 1341500 |
| P48962 | Slc25a4 | ADP/ATP translocase 1 | 0.76192 | 43900000 | 26560000 | 93559000 | 47921000 | 1464700 | 1665300 |
| A0A0A6YWP6;A0A0A6YX18; | Atp6v1h | V-type proton ATPase subunit H | 0.76357 | 1980100 | 751540 | 3489300 | 1812200 | 5223100 | 3691900 |
| A0A140LHC3;A0A140LJH1;A0 | Qars | | 0.76361 | 369810 | 202000 | 917140 | 614610 | 2532300 | 1564800 |
| A0A0N4SVM0;D6RCW7;P477 | Capza2 | F-actin-capping protein subunit alpha-2 | 0.76607 | 2565800 | 859040 | 9874000 | 11719000 | 6095000 | 3617500 |
| Q8R1B4;M0QWV3 | Eif3c | Eukaryotic translation initiation factor 3 subunit C | 0.7664 | 1200900 | 581780 | 2254100 | 1137400 | 3646900 | 2031200 |
| P20065-2;P20065 | Tmsb4x | Thymosin beta-4;Hematopoietic system regulatory peptide | 0.77177 | 16088000 | 6106300 | 20187000 | 15639000 | 1964700 | 1841600 |
| A0A087WP83;Q8VDJ3;A0A0 | Hdlbp | Vigilin | 0.77415 | 2799600 | 1253100 | 6773400 | 4969900 | 1312800 | 739310 |
| A0A1B0GSR9;A0A1B0GSX0; | Ldha | L-lactate dehydrogenase A chain | 0.77473 | 2094100 | 634050 | 4300500 | 3256500 | 5155500 | 3125800 |
| P62192 | Psmc1 | 26S protease regulatory subunit 4 | 0.77777 | 399900 | 384030 | 2671700 | 1643000 | 1507900 | 1574000 |
| Q3UMU9-4;Q3UMU9-2;Q3U | Hdgfrp2 | Hepatoma-derived growth factor-related protein 2 | 0.78199 | 0 | 0 | 1693300 | 1074700 | 1635600 | 716580 |
| P26350;A0A087WP98;A0A0 | Ptma | osin alpha;Prothymosin alpha, N-terminally processed;Thymosin alpha | 0.78572 | 4078300 | 1428600 | 14225000 | 8649100 | 9552000 | 6106400 |
| Q62465 | Vat1 | Synaptic vesicle membrane protein VAT-1 homolog | 0.7868 | 0 | 0 | 3435700 | 1454900 | 2086100 | 2139900 |
| E9Q0U7;Q61699-2;Q61699; | Hsph1 | Heat shock protein 105 kDa | 0.78761 | 1766300 | 794470 | 3673800 | 4101600 | 7927200 | 4259500 |
| Q8CB58;Q8BGJ5;Q922I7;P17 | Ptbp1 | Polypyrimidine tract-binding protein 1 | 0.78926 | 3819500 | 1870600 | 3140900 | 3155400 | 8865200 | 17868000 |
| A6ZI44;P05064;D3Z510;A0A | Aldoa;Aldoart2;Aldoart1 | uctose-bisphosphate aldolase;Fructose-bisphosphate aldolase A | 0.79094 | 4134100 | 1234800 | 12545000 | 8920600 | 15611000 | 12175000 |
| Q60737 | Csnk2a1 | Casein kinase II subunit alpha | 0.79402 | 0 | 0 | 1131300 | 1291100 | 948850 | 531950 |
| A2A817;A2A816;A2A815;A2 | Park7 | Protein deglycase DJ-1 | 0.79425 | 0 | 0 | 5802500 | 2240800 | 2166000 | 1954800 |
| A0A0U1RNQ6;Q8BFR5-2;Q8 | Tufm | Elongation factor Tu, mitochondrial | 0.79584 | 1265000 | 553410 | 2874900 | 2819700 | 2699800 | 1230900 |
| P29341;Q62029;Q9D4E6;A0 | Pabpc1 | Polyadenylate-binding protein 1 | 0.79749 | 1039500 | 561690 | 11185000 | 7988400 | 7201100 | 4438500 |
| E9Q9F5;E9Q1G8;O55131 | 07-Sep | Septin-7 | 0.79848 | 1459600 | 1428200 | 4267800 | 3979100 | 2949200 | 2404000 |
| Q9JIW9;P63321 | Ralb;Rala | Ras-related protein Ral-B;Ras-related protein Ral-A | 0.80408 | 0 | 0 | 850180 | 741560 | 1174500 | 519420 |
| P54227;D3Z5N2;D3Z1Z8 | Stmn1 | Stathmin | 0.80451 | 2365000 | 1298700 | 5240400 | 2905900 | 4244300 | 2416300 |
| P26041 | Msn | Moesin | 0.8061 | 21003000 | 7611200 | 60380000 | 101070000 | 26206000 | 9755600 |
| P26039;A2AIM2;E9PUM4;A0 | Tln1 | Talin-1 | 0.80866 | 0 | 0 | 927180 | 926560 | 10114000 | 3869100 |
| A0A140LIZ5;P54775 | Psmc4 | 26S protease regulatory subunit 6B | 0.80916 | 437500 | 231840 | 2431700 | 965420 | 1584000 | 890980 |
| Q05816 | Fabp5 | Fatty acid-binding protein, epidermal | 0.80987 | 3630300 | 1951700 | 20173000 | 16490000 | 21220000 | 13706000 |
| A0A1B0GSG5;Q91VI7;A0A1B | Rnh1 | Ribonuclease inhibitor | 0.81342 | 1946500 | 588170 | 1112200 | 1405600 | 1019400 | 1117000 |
| P62983;A0A0A6YW67;E9Q9J | Rps27a;Gm8797;Uba52;Kxd1;Ubc;Ubb | S ribosomal protein L40;Ubiquitin;60S ribosomal protein L40;Polyubiquitin-B;Ubiquitin | 0.81534 | 50257000 | 20577000 | 69851000 | 48125000 | 55049000 | 33806000 |
| Q9CR51 | Atp6v1g1 | V-type proton ATPase subunit G 1 | 0.81726 | 6432800 | 2681300 | 11997000 | 4588800 | 7831400 | 7539100 |
| A2AL12;Q8BG05-2;Q8BG05; | Hnrnpa3 | Heterogeneous nuclear ribonucleoprotein A3 | 0.81917 | 4744100 | 2157600 | 23532000 | 18092000 | 6070800 | 5709100 |
| Q62418-3;Q62418-2;Q62418 | Dbnl | Drebrin-like protein | 0.81943 | 639850 | 536980 | 1981800 | 1177900 | 2148700 | 1837300 |
| P62806 | Hist1h4a | Histone H4 | 0.82088 | 256300000 | 115230000 | 568650000 | 416500000 | 674350000 | 411570000 |
| P70372 | Elavl1 | ELAV-like protein 1 | 0.8212 | 625350 | 302430 | 1550900 | 1331100 | 2013600 | 1234600 |
| P61161 | Actr2 | Actin-related protein 2 | 0.82159 | 1638500 | 695260 | 2853100 | 2220000 | 1738300 | 964190 |

| | | | | | | | | | |
|---|---|---|---|---|---|---|---|---|---|
| P27612 | Plaa | Phospholipase A-2-activating protein | 0.82334 | 475120 | 261730 | 1251200 | 1323700 | 0 | 0 |
| P26516 | Psmd7 | 26S proteasome non-ATPase regulatory subunit 7 | 0.82356 | 1683000 | 874880 | 3885500 | 2422300 | 2764900 | 1686800 |
| B1AU25;Q9Z0X1;Q9Z0X1-2 | Aifm1 | Apoptosis-inducing factor 1, mitochondrial | 0.82396 | 2049200 | 870570 | 1816900 | 1477100 | 0 | 0 |
| Q8BFW7-4;Q8BFW7;Q8BFW | Lpp | Lipoma-preferred partner homolog | 0.83129 | 991930 | 708230 | 0 | 0 | 3285100 | 2082100 |
| Q8JZQ9 | Eif3b | Eukaryotic translation initiation factor 3 subunit B | 0.83283 | 1034500 | 927770 | 2876000 | 2222300 | 0 | 0 |
| Q60972 | Rbbp4 | Histone-binding protein RBBP4 | 0.8346 | 0 | 0 | 1602500 | 854420 | 1851900 | 1555200 |
| Q6ZWN5;F7CJS8;D3YWH9;Q | Rps9 | 40S ribosomal protein S9 | 0.83536 | 6094200 | 2338000 | 11414000 | 9562800 | 2035900 | 1339900 |
| Q9D8N0 | Eef1g | Elongation factor 1-gamma | 0.83563 | 2456800 | 115620 | 4405600 | 2889500 | 3739200 | 1143900 |
| D3YVM5;S4R1N1;P14869 | Rplp0 | 60S acidic ribosomal protein P0 | 0.83577 | 5203900 | 2602000 | 15957000 | 11777000 | 16565000 | 10205000 |
| Q78PY7;Q3TJ56;E9Q3E9 | Snd1 | Staphylococcal nuclease domain-containing protein 1 | 0.83584 | 2502000 | 1228300 | 4466600 | 3200900 | 3056600 | 2249500 |
| P97315 | Csrp1 | Cysteine and glycine-rich protein 1 | 0.83718 | 1816400 | 774890 | 4594300 | 2914300 | 3364900 | 1824200 |
| Q9QXS1-3;Q9QXS1-10;Q9QX | Plec | Plectin | 0.83776 | 20819000 | 7510300 | 55791000 | 20695000 | 55622000 | 26790000 |
| P11499;E9Q0C3;E9PX27;E9Q | Hsp90ab1 | Heat shock protein HSP 90-beta | 0.84005 | 43081000 | 17951000 | 76931000 | 49786000 | 28609000 | 25768000 |
| A0A0G2JGY8;A0A0G2JEY6;Q | Rpl34 | 60S ribosomal protein L34 | 0.84136 | 1975300 | 943830 | 10932000 | 8403200 | 11012000 | 7951000 |
| P63017;Q504P4;D3Z5E2 | Hspa8 | Heat shock cognate 71 kDa protein | 0.84249 | 43436000 | 21646000 | 127270000 | 103170000 | 81061000 | 52186000 |
| Z4YJV4;Q60597-2;Q60597;Q | Ogdh | 2-oxoglutarate dehydrogenase, mitochondrial | 0.84309 | 191600 | 167940 | 662920 | 363420 | 1701200 | 853760 |
| Q9WTM5;A0A1B0GSR4;A0A | Ruvbl2 | RuvB-like 2 | 0.84383 | 1371200 | 965860 | 961270 | 955920 | 1970900 | 1773500 |
| Q9Z204-4;Q9Z204-3;Q9Z204 | Hnrnpc | Heterogeneous nuclear ribonucleoproteins C1/C2 | 0.84458 | 7985500 | 2972600 | 4508400 | 3710200 | 4251100 | 3673300 |
| A0A3B2WDD2;A0A3B2WBL1 | Rpl10a | Ribosomal protein;60S ribosomal protein L10a | 0.84814 | 284200 | 225420 | 783880 | 552270 | 0 | 0 |
| Q8BTS0;Q61656;S4R1I6;B1A | Ddx5 | Probable ATP-dependent RNA helicase DDX5 | 0.85111 | 6483300 | 3998900 | 23975000 | 21029000 | 18050000 | 10333000 |
| P58252;G3UXK8;G3UZ34;A2 | Eef2 | Elongation factor 2 | 0.85217 | 24189000 | 11637000 | 36242000 | 27697000 | 30256000 | 22253000 |
| Q3U0I3;P80318;E9Q133;F6Q | Cct3 | T-complex protein 1 subunit gamma | 0.8523 | 338120 | 349990 | 1559700 | 1537200 | 2760300 | 1369500 |
| Q9D8S4;A0A1L1SS58;A0A1L | Rexo2 | Oligoribonuclease, mitochondrial | 0.8528 | 569940 | 298820 | 2332400 | 1362600 | 2105400 | 1771900 |
| P47911;A0A0J9YU32 | Rpl6 | 60S ribosomal protein L6 | 0.85304 | 2103900 | 1018200 | 6372800 | 4616500 | 8164900 | 4762400 |
| P62908;A0A140LI77;D3YV43 | Rps3 | 40S ribosomal protein S3 | 0.85463 | 1102400 | 502220 | 3074600 | 2023400 | 2525300 | 1173400 |
| Q9QUM9;E0CYT2;E0CXB1 | Psma6 | Proteasome subunit alpha type-6 | 0.85523 | 0 | 0 | 1262000 | 697800 | 6775900 | 4148500 |
| A0A338P6J9;Q921U7;Q8C28 | Cast | Calpastatin | 0.85925 | 664170 | 490240 | 3075300 | 2871700 | 0 | 0 |
| P99024;G3UZR1;CON__ENSI | Tubb5 | Tubulin beta-5 chain | 0.86165 | 18360000 | 7414100 | 95401000 | 66956000 | 63233000 | 63472000 |
| Q8BMS1 | Hadha | na, mitochondrial;Long-chain enoyl-CoA hydratase;Long chain 3-hydroxyacyl-CoA dehy | 0.8639 | 0 | 0 | 962950 | 827290 | 1321300 | 1012600 |
| P50518;A0A0N4SW34;A0A0I | Atp6v1e1 | V-type proton ATPase subunit E 1 | 0.86608 | 4675200 | 2121300 | 6330900 | 2366000 | 7357500 | 5344300 |
| Q8BP67 | Rpl24 | 60S ribosomal protein L24 | 0.86644 | 0 | 0 | 10452000 | 8092300 | 12077000 | 6493600 |
| Q6ZWV3;I7HLV2;P86048;A0 | Rpl10;Rpl10l | 60S ribosomal protein L10;60S ribosomal protein L10-like | 0.86734 | 5892000 | 2252100 | 10436000 | 14903000 | 4090700 | 2941900 |
| P60843;Q8BTU6;P10630;P10 | Eif4a1;Eif4a2 | A-I;Eukaryotic initiation factor 4A-II;Eukaryotic initiation factor 4A-II, N-terminally proc | 0.86954 | 13770000 | 7557200 | 14754000 | 11348000 | 29275000 | 15415000 |
| Q63844;A0A0U1RPX4;D3Z3G | Mapk3 | togen-activated protein kinase 3;Mitogen-activated protein kinase | 0.87114 | 0 | 0 | 1407200 | 1573800 | 2120400 | 1245500 |
| Q9JHJ0;A0A1L1SQ12;A0A1L1 | Tmod3 | Tropomodulin-3 | 0.87196 | 1547000 | 413880 | 0 | 0 | 1790100 | 1156300 |
| P62900;A0A0A6YXL3;A0A0A | Rpl31 | 60S ribosomal protein L31 | 0.87488 | 3651300 | 1972400 | 7058900 | 6506700 | 0 | 0 |
| P47963 | Rpl13 | 60S ribosomal protein L13 | 0.87559 | 6016100 | 3086600 | 10757000 | 9833100 | 9905800 | 6731200 |
| O08583-2;Q9JJW6-2;O08583 | Alyref;Alyref2 | THO complex subunit 4;Aly/REF export factor 2 | 0.87664 | 2073600 | 1144400 | 3264500 | 4108700 | 5058000 | 2631700 |
| P63028 | Tpt1 | Translationally-controlled tumor protein | 0.87698 | 4994300 | 2387200 | 10830000 | 6151600 | 35116000 | 31735000 |
| F6VQ81;Q3TUJ9;Q8BKP1;Q3 | Tpd52l2 | Tumor protein D54 | 0.88016 | 0 | 0 | 1337000 | 1134000 | 1428900 | 1190200 |
| Q9CQM8;O09167 | Rpl21 | 60S ribosomal protein L21 | 0.8814 | 3302300 | 1893200 | 15606000 | 14489000 | 26745000 | 17235000 |
| B2M1R6;P61979-3;P61979;P | Hnrnpk | Heterogeneous nuclear ribonucleoprotein K | 0.88229 | 2094300 | 688810 | 7569000 | 5961800 | 2481700 | 1659100 |
| E9QN08;Q80T06;A0A0R4J1E | Eef1d | Elongation factor 1-delta | 0.88404 | 1335800 | 7298600 | 17675000 | 13320000 | 19474000 | 15536000 |
| Q9CQF9;F7CIP8;D3Z275 | Pcyox1 | Prenylcysteine oxidase | 0.88777 | 0 | 0 | 1674000 | 1587900 | 1425600 | 1251900 |
| E9PZF0;Q01768;Q5NC80;P15 | Gm20390;Nme2;Nme1 | sphate kinase;Nucleoside diphosphate kinase B;Nucleoside diphosphate kinase A | 0.8882 | 24014000 | 10106000 | 61218000 | 39907000 | 26190000 | 29752000 |

| | | | | | | | | | |
|---|---|---|---|---|---|---|---|---|---|
| Q6ZWQ9;Q3THE2;D3YV37;Q | Myl12a;Myl12b | Myosin regulatory light chain 12B | 0.89045 | 874950 | 715290 | 5997200 | 6679500 | 5810900 | 3124600 |
| Q9D8E6 | Rpl4 | 60S ribosomal protein L4 | 0.89047 | 4542500 | 2041300 | 10347000 | 10813000 | 7094800 | 5003200 |
| A0A0N4SV32;Q3UMP4;Q9CY | Serbp1 | Plasminogen activator inhibitor 1 RNA-binding protein | 0.89181 | 12335000 | 6606800 | 10742000 | 7672200 | 10423000 | 6312600 |
| Q02819;A0A1B0GR41;A0A1 | Nucb1 | Nucleobindin-1 | 0.8932 | 0 | 0 | 918720 | 1091500 | 714470 | 590440 |
| A0A1B0GR11;Q93092 | Taldo1 | Transaldolase | 0.89344 | 649490 | 279870 | 1647900 | 2125100 | 1690700 | 891150 |
| A0A0A0MQC9;Q9D0R8 | Lsm12 | Protein LSM12 homolog | 0.89413 | 0 | 0 | 1616400 | 1933900 | 1861600 | 905880 |
| O88569-3;O88569-2;O88569 | Hnrnpa2b1 | Heterogeneous nuclear ribonucleoproteins A2/B1 | 0.89527 | 4115500 | 1439600 | 18512000 | 17128000 | 15414000 | 9657600 |
| Q61792;A2A6G6;A2A6G7;A2 | Lasp1 | LIM and SH3 domain protein 1 | 0.89596 | 1384400 | 680530 | 13322000 | 14196000 | 15359000 | 7681200 |
| Q9CXW4;A2BH06 | Rpl11 | 60S ribosomal protein L11 | 0.90095 | 6660200 | 3412700 | 21988000 | 14620000 | 18543000 | 8890000 |
| Q5EBP8;P49312;P49312-2 | Hnrnpa1 | onucleoprotein A1;Heterogeneous nuclear ribonucleoprotein A1, N-terminally process | 0.90158 | 17701000 | 11376000 | 37043000 | 40086000 | 48899000 | 25047000 |
| P60710;P63260;E9Q5F4;G3U | Actb;Actg1 | smic 1, N-terminally processed;Actin, cytoplasmic 2;Actin, cytoplasmic 2, N-terminally | 0.90226 | 401070000 | 202130000 | 822940000 | 870520000 | 1094100000 | 540690000 |
| Q64433;Q9JI95 | Hspe1;Cpn10-rs1 | 10 kDa heat shock protein, mitochondrial | 0.90281 | 0 | 0 | 2179500 | 1609200 | 16877000 | 11949000 |
| Q60864 | Stip1 | Stress-induced-phosphoprotein 1 | 0.90372 | 2044100 | 835550 | 3166400 | 2247000 | 948360 | 961910 |
| Q62318;Q62318-2 | Trim28 | Transcription intermediary factor 1-beta | 0.90513 | 1417100 | 934690 | 4826100 | 6697900 | 6085900 | 3488500 |
| P35979 | Rpl12 | 60S ribosomal protein L12 | 0.9059 | 2149800 | 1281400 | 3653200 | 2758900 | 30715000 | 23713000 |
| Q6P069-2;Q6P069 | Sri | Sorcin | 0.90592 | 1501900 | 550360 | 0 | 0 | 1921300 | 1443300 |
| P12970 | Rpl7a | 60S ribosomal protein L7a | 0.90777 | 2507400 | 1199700 | 11227000 | 8591400 | 5807500 | 3624000 |
| Q9CWK8 | Snx2 | Sorting nexin-2 | 0.90861 | 863230 | 481670 | 1819200 | 2244200 | 2893600 | 1594700 |
| P80316;E0CZA1 | Cct5 | T-complex protein 1 subunit epsilon | 0.90957 | 503910 | 323040 | 5449000 | 4530600 | 8304500 | 5875100 |
| Q01853 | Vcp | Transitional endoplasmic reticulum ATPase | 0.91254 | 20327000 | 8977600 | 37992000 | 30791000 | 55917000 | 45853000 |
| Q62261;A0A0A0MQG2;Q622 | Sptbn1 | Spectrin beta chain, non-erythrocytic 1 | 0.91495 | 1652400 | 1337800 | 5819300 | 6037100 | 17470000 | 9199800 |
| Q91VW3;I7HPY0 | Sh3bgrl3 | SH3 domain-binding glutamic acid-rich-like protein 3 | 0.91539 | 13405000 | 7588100 | 68336000 | 53178000 | 278710 | 1835400 |
| P51881 | Slc25a5 | /ATP translocase 2;ADP/ATP translocase 2, N-terminally processed | 0.91605 | 22307000 | 9875100 | 34823000 | 23743000 | 10958000 | 9318000 |
| P67984 | Rpl22 | 60S ribosomal protein L22 | 0.91811 | 1766600 | 858110 | 4431500 | 3011700 | 2264500 | 1867900 |
| Q06185;Q8BTB6 | Atp5i;Atp5k | ATP synthase subunit e, mitochondrial | 0.9198 | 3815400 | 2145900 | 8625400 | 7024300 | 7657800 | 5775900 |
| Q8BH80;Q9QY76 | Vapb | Vesicle-associated membrane protein-associated protein B | 0.92358 | 0 | 0 | 2384300 | 2290800 | 4831000 | 3511300 |
| Q3UA95;Q3THW5;P0C0S6;Q | H2afz;H2afv | Histone H2A;Histone H2A.V;Histone H2A.Z | 0.92369 | 307590 | 227360 | 1337800 | 1252300 | 0 | 0 |
| P14148;F6XI62 | Rpl7 | 60S ribosomal protein L7 | 0.92528 | 610490 | 371510 | 9702600 | 6651100 | 9197100 | 6135900 |
| Q61598-2;Q61598;A0A1Y7VI | Gdi2 | Rab GDP dissociation inhibitor beta | 0.92582 | 9785100 | 5048400 | 33093000 | 20271000 | 28706000 | 19705000 |
| P62751 | Rpl23a | 60S ribosomal protein L23a | 0.92842 | 11621000 | 6237100 | 15403000 | 14595000 | 84671000 | 31978000 |
| P05213;A0A2R8VHF3 | Tuba1b | Tubulin alpha-1B chain | 0.93121 | 47008000 | 21534000 | 65094000 | 41092000 | 14042000 | 13054000 |
| Q9JHU4;F6ZX84 | Dync1h1 | Cytoplasmic dynein 1 heavy chain 1 | 0.93183 | 4550600 | 3279300 | 9622300 | 4924000 | 5802200 | 4444600 |
| A2AFI4;A2AFI3;A0A2I3BRL8; | Rbmx;Rbmxl1 | NA-binding motif protein, X chromosome, N-terminally processed;RNA binding motif | 0.93206 | 1272800 | 938090 | 5007200 | 3435400 | 3476500 | 2970400 |
| P60867 | Rps20 | 40S ribosomal protein S20 | 0.93264 | 2481000 | 1634000 | 17760000 | 15948000 | 14096000 | 10085000 |
| P61982 | Ywhag | -3 protein gamma;14-3-3 protein gamma, N-terminally processed | 0.93499 | 1497200 | 896540 | 27470000 | 22209000 | 29934000 | 20730000 |
| P47757-2;P47757;P47757-4; | Capzb | F-actin-capping protein subunit beta | 0.93585 | 1504200 | 1005600 | 14755000 | 14724000 | 26052000 | 17909000 |
| P43274;I7HFT9;Q07133;P15 | Hist1h1e | Histone H1.4 | 0.93686 | 14867000 | 2023900 | 37974000 | 11030000 | 17115000 | 11128000 |
| P08228 | Sod1 | Superoxide dismutase [Cu-Zn] | 0.93711 | 9408600 | 5695300 | 29423000 | 28903000 | 29354000 | 14928000 |
| E9Q4Q2;D3YZC9;D3YZD0;Q6 | Sf1 | Splicing factor 1 | 0.93811 | 1637500 | 548390 | 2201700 | 2362900 | 5204500 | 2920900 |
| A0A1B0GSB2;P19253;A0A1B | Rpl13a | 60S ribosomal protein L13a | 0.93921 | 2698300 | 1394900 | 7404400 | 7976500 | 0 | 0 |
| P16460 | Ass1 | Argininosuccinate synthase | 0.94043 | 531910 | 294280 | 1658800 | 394960 | 1595900 | 2030800 |
| P13020-2;P13020;A0A0J9YU | Gsn | Gelsolin | 0.94184 | 0 | 0 | 2306800 | 2974300 | 0 | 0 |
| Q3UJB0;A0A494B9S9 | Sf3b2 | | 0.94509 | 0 | 0 | 2002600 | 2112000 | 16627000 | 481640 |
| P63038;D3Z7J9;D3Z2F2;P630 | Hspd1 | 60 kDa heat shock protein, mitochondrial | 0.94766 | 240780 | 108870 | 8109200 | 6606200 | 13175000 | 23608000 |
| Q6ZWZ7;Q9CPR4 | Rpl17 | 60S ribosomal protein L17 | 0.9482 | 2906100 | 1080100 | 3198500 | 3023100 | 2224600 | 820830 |

| | | | | | | | | | | |
|---|---|---|---|---|---|---|---|---|---|---|
| P62082 | Rps7 | 40S ribosomal protein S7 | | 0.9491 | 1772200 | 809680 | 12825000 | 8615300 | 10864000 | 7987200 |
| Q3UGB5;Q9JII5-2;Q9JII5;D3Z | Dazap1 | DAZ-associated protein 1 | | 0.94919 | 1217500 | 740700 | 2460400 | 2164100 | 1907300 | 2324800 |
| P20108 | Prdx3 | Thioredoxin-dependent peroxide reductase, mitochondrial | | 0.94952 | 0 | 0 | 0 | 0 | 10878000 | 7360900 |
| P14115 | Rpl27a | 60S ribosomal protein L27a | | 0.95012 | 2125700 | 1357500 | 9204600 | 8205700 | 17417000 | 11273000 |
| P68510 | Ywhah | 14-3-3 protein eta | | 0.95423 | 755840 | 470480 | 2804700 | 2005500 | 0 | 0 |
| P17182;Q6PHC1;B0QZL1;B1A | Eno1 | Alpha-enolase;Enolase | | 0.95505 | 30094000 | 14491000 | 62192000 | 59905000 | 70662000 | 37824000 |
| Q9CZY3;B7ZBY7;E9PY39;Q9C | Ube2v1;Gm20431 | Ubiquitin-conjugating enzyme E2 variant 1 | | 0.95749 | 2224400 | 1376800 | 5494400 | 4062700 | 6056000 | 3997600 |
| A2A547;P84099 | Rpl19 | Ribosomal protein L19;60S ribosomal protein L19 | | 0.95818 | 10832000 | 3222600 | 25419000 | 18032000 | 19675000 | 12763000 |
| A0A1L1SUX8;P01831 | Thy1 | Thy-1 membrane glycoprotein | | 0.95874 | 0 | 0 | 2237800 | 2151000 | 7092500 | 4861600 |
| P43276 | Hist1h1b | Histone H1.5 | | 0.96097 | 2526300 | 1384500 | 6696800 | 5564800 | 4913100 | 3497400 |
| P62264;D3YVF4;D3Z7I1 | Rps14 | 40S ribosomal protein S14 | | 0.96259 | 5420200 | 2700000 | 11246000 | 8337100 | 9619300 | 6584000 |
| D3YVX4;Q91VZ6 | Smap1 | Stromal membrane-associated protein 1 | | 0.96355 | 768730 | 780750 | 1911100 | 1353300 | 1932800 | 998820 |
| Q8VDN2;D3YYN7;A0A0G2JG | Atp1a1 | Sodium/potassium-transporting ATPase subunit alpha-1 | | 0.966 | 762160 | 459520 | 6224000 | 7185800 | 6042600 | 4282600 |
| Q9D3D9 | Atp5d | ATP synthase subunit delta, mitochondrial | | 0.96663 | 0 | 0 | 4243600 | 3599000 | 5574500 | 3966200 |
| Q9CZN7-2;Q9CZN7 | Shmt2 | Serine hydroxymethyltransferase | | 0.96667 | 1632700 | 515710 | 2780500 | 3342300 | 0 | 0 |
| CON__P04264;E9Q0F0 | | | + | 0.96947 | 7141700 | 2746200 | 44218000 | 19954000 | 13729000 | 22556000 |
| P20029 | Hspa5 | 78 kDa glucose-regulated protein | | 0.96954 | 25718000 | 12414000 | 56300000 | 50911000 | 54401000 | 41593000 |
| Q03265;D3Z6F5;D6RJ16 | Atp5a1 | synthase subunit alpha, mitochondrial;ATP synthase subunit alpha | | 0.97204 | 7705900 | 3677600 | 49448000 | 40533000 | 35041000 | 24494000 |
| F6YVP7;P62270;S4R1N6;A0A | Gm10260;Rps18 | 40S ribosomal protein S18 | | 0.97243 | 2445500 | 1020700 | 9071500 | 7696500 | 13400000 | 8070300 |
| P62960;A2BGG7;A0A0A0MC | Ybx1 | Nuclease-sensitive element-binding protein 1 | | 0.97257 | 7910400 | 4118400 | 10788000 | 10375000 | 1423600 | 1367600 |
| Q8VEK3-2;Q8VEK3 | Hnrnpu | Heterogeneous nuclear ribonucleoprotein U | | 0.97444 | 3315800 | 1411500 | 12959000 | 12977000 | 14362000 | 7554500 |
| Q6ZWV7 | Rpl35 | 60S ribosomal protein L35 | | 0.97616 | 6881200 | 3395100 | 14545000 | 11636000 | 845920 | 647540 |
| P62702 | Rps4x | 40S ribosomal protein S4, X isoform | | 0.97723 | 4748400 | 2592800 | 7729200 | 6691600 | 10636000 | 5966400 |
| Q9CQV8-2;Q9CQV8;A2A5N1 | Ywhab | protein beta/alpha;14-3-3 protein beta/alpha, N-terminally processed | | 0.97764 | 5253000 | 2985100 | 7569700 | 6857300 | 5082000 | 3631200 |
| Q60837 | Il12rb1 | Interleukin-12 receptor subunit beta-1 | | 0.9788 | 0 | 0 | 278920 | 288510 | 1560400 | 842920 |
| P80314;A0A1W2P7B7;A0A1\ | Cct2 | T-complex protein 1 subunit beta | | 0.98209 | 1547500 | 749050 | 4279500 | 4335700 | 5019900 | 3992800 |
| Q60692 | Psmb6 | Proteasome subunit beta type-6 | | 0.98524 | 587110 | 397050 | 2065100 | 2191300 | 3411500 | 2176100 |
| P14152;A0A5F8MPN8;B1ATC | Mdh1 | Malate dehydrogenase, cytoplasmic | | 0.9854 | 0 | 0 | 700850 | 517810 | 2067400 | 1976300 |
| P08113;F7C312;D3Z1R1 | Hsp90b1 | Endoplasmin | | 0.98684 | 18778000 | 10733000 | 27787000 | 34565000 | 20288000 | 23410000 |
| P09411;S4R2M7;P09041 | Pgk1;Pgk2 | glycerate kinase 1;Phosphoglycerate kinase;Phosphoglycerate kinase 2 | | 0.98826 | 20034000 | 9015500 | 65275000 | 52603000 | 67833000 | 59525000 |
| A0A1B0GQU8;P35980;A0A1I | Rpl18 | 60S ribosomal protein L18 | | 0.98896 | 8783800 | 4738000 | 13157000 | 11536000 | 5726600 | 3869700 |
| P09405 | Ncl | Nucleolin | | 0.98946 | 6409500 | 1839000 | 20697000 | 16678000 | 13333000 | 8075200 |
| Q99PT1 | Arhgdia | Rho GDP-dissociation inhibitor 1 | | 0.994 | 9258000 | 6016200 | 13881000 | 11319000 | 10731000 | 7822200 |
| Q9DB20;F7D3P8;A0A338P77 | Atp5o | ATP synthase subunit O, mitochondrial | | 0.99567 | 999740 | 509150 | 2777700 | 2575200 | 2744300 | 2356300 |
| P0DP28;P0DP27;P0DP26;A0A | Calml3 | Calmodulin-like protein 3 | | 0.99757 | 248570000 | 134090000 | 326460000 | 284500000 | 98629000 | 62279000 |
| P47962;D3YYV8 | Rpl5 | 60S ribosomal protein L5 | | 1.0025 | 11904000 | 5978300 | 21065000 | 16730000 | 36410000 | 24642000 |
| G3UVV4;P17710-3;P17710-4 | Hk1 | Hexokinase;Hexokinase-1 | | 1.003 | 2992900 | 1594500 | 5616700 | 3515900 | 2610100 | 2251300 |
| Q9WUK2-2;Q9WUK2 | Eif4h | Eukaryotic translation initiation factor 4H | | 1.004 | 3269100 | 1659800 | 7498400 | 6578800 | 0 | 0 |
| A0A0G2JES3;A0A140T8T4;P5 | Rpl9 | 60S ribosomal protein L9 | | 1.0077 | 0 | 0 | 1664500 | 2077700 | 1903100 | 2708600 |
| Q9CQR2 | Rps21 | 40S ribosomal protein S21 | | 1.0081 | 2979000 | 2108100 | 6618400 | 6400000 | 9684700 | 6637100 |
| P62267 | Rps23 | 40S ribosomal protein S23 | | 1.0131 | 3210500 | 2137700 | 6036700 | 5577700 | 348600 | 216940 |
| Q8VDD5;A0A2R8W6V7;A0A2 | Myh9 | Myosin-9 | | 1.0135 | 27208000 | 15226000 | 72087000 | 90002000 | 148050000 | 63468000 |
| A1BN54;Q7TPR4;O88990 | Actn1 | Alpha-actinin-1 | | 1.0145 | 2597700 | 950310 | 12254000 | 9808100 | 11188000 | 4528200 |
| Q8VIJ6 | Sfpq | Splicing factor, proline- and glutamine-rich | | 1.0176 | 389920 | 457270 | 5321700 | 4064000 | 6915400 | 5533000 |
| P50396;D6RI86;B7FAU8 | Gdi1 | Rab GDP dissociation inhibitor alpha | | 1.0189 | 4573300 | 1553000 | 1704000 | 1330900 | 0 | 0 |

| | | | | | | | | | | |
|---|---|---|---|---|---|---|---|---|---|---|
| A3KGU7;A3KGU9;P16546;A3 | Sptan1 | Spectrin alpha chain, non-erythrocytic 1 | | 1.0213 | 7160900 | 5089100 | 19796000 | 10136000 | 9393900 | 7321500 |
| O08749 | Dld | Dihydrolipoyl dehydrogenase, mitochondrial | | 1.0214 | 829340 | 610030 | 1436100 | 1729700 | 3567700 | 2876300 |
| Q80YQ1;P35441 | Thbs1 | Thrombospondin-1 | | 1.0273 | 0 | 0 | 358250 | 2375200 | 13425000 | 12894000 |
| S4R1W1;A0A0A0MQF6;P168 | Gm3839;Gapdh | Glyceraldehyde-3-phosphate dehydrogenase | | 1.0285 | 23145000 | 11946000 | 49313000 | 36896000 | 71338000 | 45695000 |
| B7FAU9;Q8BTM8;B7FAV1;F | Flna | Filamin-A | | 1.0309 | 5645800 | 1193000 | 20656000 | 7744100 | 13946000 | 10006000 |
| P63325;A0A338P731;Q3UW | Rps10 | 40S ribosomal protein S10 | | 1.0319 | 0 | 0 | 3445800 | 3356200 | 2062600 | 1766900 |
| P08249;A0A0G2JF23;A0A0G | Mdh2 | Malate dehydrogenase, mitochondrial | | 1.0378 | 1348200 | 751750 | 3834600 | 3904200 | 5725400 | 80384000 |
| P68372;Q9D6F9 | Tubb4b;Tubb4a | Tubulin beta-4B chain;Tubulin beta-4A chain | | 1.0384 | 2159600 | 1773300 | 24503000 | 28324000 | 30713000 | 31184000 |
| P62259;F6WA09;D6REF3 | Ywhae | 14-3-3 protein epsilon | | 1.0417 | 555720 | 406870 | 2679500 | 1983300 | 0 | 0 |
| A0A494BA97;Q62422 | Ostf1 | Osteoclast-stimulating factor 1 | | 1.0431 | 285080 | 96422 | 668310 | 363620 | 450850 | 687900 |
| Q91VD1-2 | | | | 1.0439 | 3559700 | 1742500 | 5329700 | 4821400 | 4328100 | 4066200 |
| P26443;F7CFA5 | Glud1 | Glutamate dehydrogenase 1, mitochondrial | | 1.0449 | 1245700 | 738180 | 4971000 | 4325400 | 2207500 | 1700600 |
| Q9DB05;A0A1B0GR35;P286 | Napa | Alpha-soluble NSF attachment protein | | 1.0476 | 0 | 0 | 2165900 | 4831000 | 0 | 0 |
| P62242 | Rps8 | 40S ribosomal protein S8 | | 1.0479 | 2281800 | 1575500 | 3647400 | 4113800 | 5926100 | 4852000 |
| P14131 | Rps16 | 40S ribosomal protein S16 | | 1.0498 | 3469400 | 2671200 | 8075800 | 6797400 | 6565200 | 5490200 |
| A0A1W2P777;A0A0G2JDL9;( | Rap1b;Rap1a | Ras-related protein Rap-1b;Ras-related protein Rap-1A | | 1.0512 | 665990 | 389950 | 996830 | 810780 | 15561000 | 13544000 |
| F7BHJ0;Q6IRU5-3;Q6IRU5-2 | Cltb | Clathrin light chain B | | 1.0529 | 1253000 | 595130 | 2323400 | 2246500 | 2363300 | 2885700 |
| A2BE92;A2BE93;Q9EQU5-2;( | Set | Protein SET | | 1.0573 | 2643400 | 1552800 | 6022400 | 7260300 | 5701700 | 3441000 |
| Q9CPQ1 | Cox6c | Cytochrome c oxidase subunit 6C | | 1.0575 | 0 | 0 | 1893300 | 1790600 | 1559100 | 1448900 |
| A0A1W2P6F6;A0A1W2P7Q9 | Myl6 | Myosin light polypeptide 6 | | 1.0582 | 2805800 | 2210000 | 10570000 | 13482000 | 11879000 | 14306000 |
| A0A494B969;A0A494BBK1;A | Add3 | Gamma-adducin | | 1.0604 | 798600 | 626550 | 1805300 | 1095500 | 1898800 | 1902400 |
| P34884 | Mif | Macrophage migration inhibitory factor | | 1.0613 | 3377200 | 699540 | 5458400 | 6560500 | 3071800 | 3502400 |
| O88531;B1B0P8;B1B0P9 | Ppt1 | Palmitoyl-protein thioesterase 1 | | 1.0626 | 1006700 | 735080 | 1718800 | 1692900 | 1663300 | 1095800 |
| A2AVJ7;Q99PL5 | Rrbp1 | Ribosome-binding protein 1 | | 1.0759 | 0 | 0 | 3525500 | 3780800 | 3984800 | 3364200 |
| P27773;F6Q404 | Pdia3 | Protein disulfide-isomerase A3 | | 1.0814 | 40369000 | 7115100 | 41025000 | 40199000 | 51473000 | 40840000 |
| P97351 | Rps3a | 40S ribosomal protein S3a | | 1.0817 | 0 | 0 | 673190 | 561400 | 870150 | 769440 |
| P57759 | Erp29 | Endoplasmic reticulum resident protein 29 | | 1.0889 | 0 | 0 | 2213800 | 2691600 | 2953200 | 2596300 |
| Q9DCX2;B1ASE2 | Atp5h | ATP synthase subunit d, mitochondrial | | 1.0895 | 0 | 0 | 4865400 | 4224800 | 8625800 | 5969400 |
| F8WJG3;P62996 | Tra2b | Transformer-2 protein homolog beta | | 1.0905 | 582330 | 306020 | 1212300 | 2201900 | 0 | 0 |
| P97379-2;P97379 | G3bp2 | Ras GTPase-activating protein-binding protein 2 | | 1.0909 | 1285100 | 989570 | 1949900 | 2218500 | 2464800 | 2044700 |
| P09103;E9Q8G8 | P4hb | Protein disulfide-isomerase | | 1.0944 | 12260000 | 7240400 | 54203000 | 19286000 | 33837000 | 21731000 |
| Q61599 | Arhgdib | Rho GDP-dissociation inhibitor 2 | | 1.0948 | 927190 | 638860 | 2226000 | 1298400 | 1181400 | 1988000 |
| P27546;P27546-3;P27546-2; | Map4 | Microtubule-associated protein 4 | | 1.0953 | 5415400 | 3601600 | 8349000 | 5697700 | 4615500 | 4201000 |
| A2ATP6;Q8C854-1;Q8C854-2 | Myef2 | Myelin expression factor 2 | | 1.0999 | 957240 | 479960 | 1582900 | 2152000 | 2899400 | 3104500 |
| Q9D0J8 | Ptms | Parathymosin | | 1.1001 | 7896900 | 2162400 | 135410000 | 110060000 | 100660000 | 72474000 |
| H7BX22;P34022 | Ranbp1 | Ran-specific GTPase-activating protein | | 1.101 | 1753600 | 1102000 | 3928500 | 4234000 | 5735100 | 3751700 |
| P63276 | Rps17 | 40S ribosomal protein S17 | | 1.1018 | 2723200 | 1368600 | 7482900 | 8708100 | 13138000 | 10556000 |
| P12787 | Cox5a | Cytochrome c oxidase subunit 5A, mitochondrial | | 1.1021 | 1472500 | 1001200 | 8396900 | 6684000 | 24815000 | 17185000 |
| A2AAN2;Q8BMA6 | Srp68 | Signal recognition particle subunit SRP68 | | 1.1082 | 703410 | 438880 | 1424500 | 1035800 | 931120 | 832950 |
| Q9CZU6;Q80X68 | Cs;Csl | Citrate synthase, mitochondrial;Citrate synthase | | 1.1138 | 1024800 | 568210 | 2803700 | 2701500 | 3400700 | 2103500 |
| CON__P34955 | | | + | 1.1157 | 2328700 | 1693600 | 5355000 | 3916900 | 5059500 | 4381900 |
| Q3U2G2;Q61316 | Hspa4 | Heat shock 70 kDa protein 4 | | 1.1218 | 4229900 | 1929200 | 3505500 | 5082200 | 2832600 | 2921500 |
| A0A1L1SQA8;P62852 | Rps25 | 40S ribosomal protein S25 | | 1.122 | 14075000 | 10005000 | 83426000 | 79628000 | 67495000 | 43999000 |
| A0A140LIU3;P56391 | Cox6b1 | Cytochrome c oxidase subunit 6B1 | | 1.1272 | 2380800 | 1651300 | 0 | 0 | 3291200 | 2160800 |
| Q9DBJ1 | Pgam1 | Phosphoglycerate mutase 1 | | 1.1316 | 9032500 | 5204800 | 17786000 | 21171000 | 106290000 | 57160000 |

| | | | | | | | | | | |
|---|---|---|---|---|---|---|---|---|---|---|
| G3UXT7;Q8CFQ9;P56959;G3 | Fus | RNA-binding protein FUS | | 1.1357 | 1695000 | 1422100 | 5331800 | 4822400 | 0 | 0 |
| P17742;A0A1L1SRX5;A0A1L1 | Ppia | -trans isomerase A;Peptidyl-prolyl cis-trans isomerase A, N-terminally processed | | 1.1358 | 13457000 | 6884700 | 37594000 | 27444000 | 127210000 | 83956000 |
| P24369 | Ppib | Peptidyl-prolyl cis-trans isomerase B | | 1.1397 | 1676000 | 1173000 | 14663000 | 16305000 | 3438200 | 2358500 |
| Q80X90 | Flnb | Filamin-B | | 1.1404 | 29907000 | 18967000 | 53417000 | 69045000 | 76516000 | 49491000 |
| Q5SX49;P62962;CON__P025 | Pfn1 | Profilin;Profilin-1 | + | 1.141 | 3443300 | 2347300 | 17720000 | 19594000 | 17799000 | 13933000 |
| P63101;D3YXN6;A0A2I3BQ0 | Ywhaz | 14-3-3 protein zeta/delta | | 1.1425 | 2747400 | 1691500 | 2913200 | 3812300 | 0 | 0 |
| A2AVR9;P62627 | Dynlrb1 | Dynein light chain roadblock-type 1 | | 1.1449 | 1558900 | 1105900 | 2445600 | 2711700 | 4620100 | 2726200 |
| P70349;B0R1E3 | Hint1 | Histidine triad nucleotide-binding protein 1 | | 1.1472 | 3981000 | 2339100 | 8575900 | 8296900 | 13944000 | 10754000 |
| Q9CQA3 | Sdhb | nate dehydrogenase [ubiquinone] iron-sulfur subunit, mitochondrial | | 1.1484 | 0 | 0 | 1795300 | 1482300 | 1443200 | 1560500 |
| P18760;F8WGL3;A0A494B9A | Cfl1 | Cofilin-1 | | 1.153 | 7890700 | 4419700 | 22477000 | 23672000 | 34493000 | 20842000 |
| D3YUT3;D3YUG3;D3Z5R8;D3 | Rps19 | 40S ribosomal protein S19 | | 1.1622 | 6213500 | 4809800 | 15614000 | 20412000 | 3250200 | 2177800 |
| Q8BH64 | Ehd2 | EH domain-containing protein 2 | | 1.1639 | 3449200 | 2663700 | 7114700 | 6090600 | 10515000 | 18088000 |
| Q80XR6;Q20BD0;Q99020 | Hnrnpab | Heterogeneous nuclear ribonucleoprotein A/B | | 1.164 | 1790700 | 1656200 | 5651300 | 9227400 | 7706600 | 9292200 |
| Q8BKE0;P49722 | Psma2 | Proteasome subunit alpha type-2 | | 1.165 | 1017800 | 701450 | 2840800 | 3865100 | 2058400 | 1629600 |
| K7N6B7;K9J7H2;K7N6J4;D3Y1201;Vmn1r121;Gm4214;Vmn1r126;Vmn1 | | Taste receptor type 2 | | 1.1685 | 0 | 0 | 1031100 | 1166000 | 2824100 | 1506900 |
| P62827 | Ran | GTP-binding nuclear protein Ran | | 1.1689 | 0 | 0 | 2323400 | 2403900 | 5729000 | 3477100 |
| Q6GT24;D3Z0Y2;O08709;Q8 | Prdx6 | Peroxiredoxin-6 | | 1.1775 | 0 | 0 | 3547700 | 5344000 | 0 | 0 |
| H3BKL5;Q921H8;H3BKA1;H3 | Acaa1a;Acaa1b | cyl-CoA thiolase A, peroxisomal;3-ketoacyl-CoA thiolase B, peroxisomal | | 1.1779 | 377810 | 345850 | 1281300 | 1894900 | 2772900 | 2599400 |
| Q6IRU2;A0A571BEU1 | Tpm4 | Tropomyosin alpha-4 chain | | 1.1803 | 30741000 | 24250000 | 41251000 | 50557000 | 52205000 | 31659000 |
| Q61171;D3Z4A4 | Prdx2 | Peroxiredoxin-2 | | 1.1827 | 1082600 | 456280 | 3809100 | 2634900 | 20838000 | 15004000 |
| H3BLF7;H3BKR2;P62874 | Gnb1 | uanine nucleotide-binding protein G(I)/G(S)/G(T) subunit beta-1 | | 1.1949 | 0 | 0 | 899980 | 1940900 | 1275000 | 744270 |
| D3Z6P1;D6RCU8;Q9DCL9 | Paics | horibosylaminoimidazole-succinocarboxamide synthase;Phosphoribosylaminoimidazol | | 1.1989 | 0 | 0 | 1491300 | 1040800 | 2001900 | 2994900 |
| P04117 | Fabp4 | Fatty acid-binding protein, adipocyte | | 1.1992 | 2680400 | 1399400 | 11925000 | 15105000 | 5160500 | 4392000 |
| Q8BVQ9;P46471 | Psmc2 | 26S protease regulatory subunit 7 | | 1.2009 | 760440 | 547610 | 0 | 0 | 4434100 | 1631100 |
| Q3TJD7;B8JJB3;Q8BVJ7;Q3T | Pdlim7 | PDZ and LIM domain protein 7 | | 1.2047 | 0 | 0 | 2957100 | 3824500 | 3363100 | 2474700 |
| A0A2I3BPS1;Q9CQI3;A0A2I3 | Gmfb | Glia maturation factor beta | | 1.208 | 0 | 0 | 456690 | 568580 | 635140 | 795780 |
| Q8BMK4 | Ckap4 | Cytoskeleton-associated protein 4 | | 1.2107 | 1400900 | 1175100 | 4370800 | 5238600 | 12999000 | 8198600 |
| P62245;F8WJ41;D3YVB4;D3 | Rps15a | 40S ribosomal protein S15a | | 1.2142 | 0 | 0 | 6759700 | 7511400 | 1373400 | 994100 |
| Q9CQX2 | Cyb5b | Cytochrome b5 type B | | 1.2213 | 9538700 | 5620300 | 4053300 | 2551500 | 13739000 | 14858000 |
| A2AE89;P10649;F6WHQ7;D3 | Gstm1 | Glutathione S-transferase Mu 1 | | 1.2245 | 0 | 0 | 936830 | 1885100 | 1319100 | 1039100 |
| P62754 | Rps6 | 40S ribosomal protein S6 | | 1.2256 | 0 | 0 | 5103400 | 4900300 | 8211000 | 7708000 |
| P68254-2;P68254 | Ywhaq | 14-3-3 protein theta | | 1.2314 | 1405400 | 630310 | 7211600 | 3655000 | 9473500 | 1421400 |
| Q921R2;P62301;A0A0U1RQ | Rps13 | 40S ribosomal protein S13 | | 1.2317 | 1129500 | 820440 | 1865900 | 3054900 | 0 | 0 |
| E9Q1K3;F8WHZ9;F8WGR0;C | Add1 | Alpha-adducin | | 1.2332 | 1252100 | 729020 | 1948800 | 4049400 | 4577600 | 4253400 |
| F8WIT2;P14824 | Anxa6 | Annexin;Annexin A6 | | 1.2358 | 1042000 | 455740 | 1382800 | 1987500 | 2411000 | 9448200 |
| O35639;Q3TET3;A0A0G2JGL | Anxa3 | Annexin A3;Annexin | | 1.2379 | 447770 | 1196700 | 0 | 0 | 8281300 | 4303700 |
| A0A5F8MPK9;Q9EP69 | Sacm1l | Phosphatidylinositide phosphatase SAC1 | | 1.2433 | 1933200 | 1339800 | 3544400 | 4251400 | 3455700 | 2911600 |
| Q8R411 | Myct1 | Myc target protein 1 | | 1.2437 | 0 | 0 | 0 | 0 | 2586900 | 2334800 |
| B0V2N7;P07356;B0V2N5;B0 | Anxa2 | Annexin;Annexin A2 | | 1.2591 | 2730700 | 2431400 | 14293000 | 12131000 | 12943000 | 9047100 |
| P48722-2;P48722;E0CY23 | Hspa4l | Heat shock 70 kDa protein 4L | | 1.2886 | 0 | 0 | 1721600 | 1758400 | 1141500 | 1296200 |
| CON__P02769 | | | + | 1.2928 | 296490 | 341940 | 873810 | 1089600 | 1235100 | 959490 |
| P48678;P48678-2;P48678-3; | Lmna | Prelamin-A/C;Lamin-A/C | | 1.2939 | 4438800 | 2247300 | 8106400 | 10831000 | 9351700 | 6046800 |
| Q9R0P5 | Dstn | Destrin | | 1.2944 | 0 | 0 | 10780000 | 14445000 | 14561000 | 12374000 |
| Q9WVA4;A0A0A6YXG6;Q9R | Tagln2 | Transgelin-2 | | 1.2964 | 4548300 | 2481900 | 25106000 | 40701000 | 68021000 | 32947000 |
| Q9CQ60;F6X8L5;Q8CBG6;D3 | Pgls | 6-phosphogluconolactonase | | 1.2979 | 1418600 | 1095400 | 2846500 | 2574700 | 2304700 | 2852300 |

| | | | | | | | | | | |
|---|---|---|---|---|---|---|---|---|---|---|
| P43346 | Dck | Deoxycytidine kinase | | 1.3098 | 19451000 | 12970000 | 0 | 0 | 48565000 | 43991000 |
| P14206;A0A1L1SRW0;A0A1L | Rpsa | 40S ribosomal protein SA | | 1.313 | 4759000 | 13199000 | 11892000 | 15103000 | 13937000 | 10877000 |
| O08795;O08795-2 | Prkcsh | Glucosidase 2 subunit beta | | 1.3228 | 1217400 | 641790 | 811710 | 1905600 | 0 | 0 |
| P56480 | Atp5b | ATP synthase subunit beta, mitochondrial | | 1.3444 | 14485000 | 6815600 | 24448000 | 18119000 | 19600000 | 13886000 |
| A0A1L1SV25;P57780;E9Q2W | Actn4 | Alpha-actinin-4 | | 1.3498 | 16581000 | 11537000 | 44455000 | 60350000 | 83516000 | 39949000 |
| P56382 | Atp5e | ATP synthase subunit epsilon, mitochondrial | | 1.3558 | 0 | 0 | 1634700 | 2470400 | 10135000 | 4287600 |
| O70400;S4R1V0 | Pdlim1 | PDZ and LIM domain protein 1 | | 1.3578 | 2619100 | 1255500 | 9872300 | 43027000 | 15066000 | 5089000 |
| D3Z7R6;Q08093 | Cnn2 | Calponin;Calponin-2 | | 1.3619 | 1057600 | 1165700 | 3529500 | 4071700 | 7364300 | 3072400 |
| Q8VHX6-2;Q8VHX6 | Flnc | Filamin-C | | 1.3635 | 2120000 | 1989600 | 4301600 | 5002200 | 6383700 | 6157500 |
| Q3UHL6;A0A087WR50;A0A0 | Fn1 | Fibronectin;Anastellin | | 1.365 | 22533000 | 17921000 | 65953000 | 180160000 | 238890000 | 62264000 |
| G3X973;Q8R4Y4-2;Q8R4Y4;F | Stab1 | Stabilin-1 | | 1.3782 | 689850 | 461640 | 3839500 | 4895400 | 14863000 | 5061500 |
| G3UWA1;Q5U465-2;Q5U465 | Ccdc125 | Coiled-coil domain-containing protein 125 | | 1.3787 | 0 | 0 | 48897000 | 66881000 | 65577000 | 62362000 |
| P07901;B7ZC50 | Hsp90aa1 | Heat shock protein HSP 90-alpha | | 1.381 | 1425000 | 739050 | 2310000 | 4714700 | 3737800 | 3035200 |
| Q8BP92 | Rcn2 | Reticulocalbin-2 | | 1.3873 | 1067100 | 1005500 | 1566100 | 1698200 | 2904400 | 3497000 |
| B1ARA3;B1ARA5;P61255 | Rpl26 | 60S ribosomal protein L26 | | 1.3912 | 1681900 | 1302700 | 6385100 | 4953600 | 5908200 | 7469300 |
| Q8K297 | Colgalt1 | Procollagen galactosyltransferase 1 | | 1.406 | 1130800 | 2083200 | 1145300 | 908430 | 1089700 | 4388100 |
| A2APB8 | Tpx2 | Targeting protein for Xklp2 | | 1.4336 | 1466200 | 1186700 | 2670800 | 11756000 | 2635900 | 6666300 |
| E9QP09;S4R1Y1;Q80TT8-4;Q | Cul9 | Cullin-9 | | 1.4348 | 0 | 0 | 3673700 | 5029200 | 4645200 | 5077900 |
| P35564 | Canx | Calnexin | | 1.435 | 0 | 0 | 2595700 | 4176200 | 2264900 | 1707000 |
| Q8VCQ8;E9QA16;F6RGN9;F6 | Cald1 | | | 1.4491 | 3696600 | 1988300 | 7727900 | 9721700 | 7326800 | 4531900 |
| O88325 | Naglu | | | 1.4545 | 432640 | 551870 | 2002300 | 1685700 | 0 | 0 |
| Q9CWF2 | Tubb2b | Tubulin beta-2B chain | | 1.4596 | 0 | 0 | 1025000 | 1762300 | 3175400 | 3125800 |
| P48036;A0A0G2JGQ0 | Anxa5 | Annexin A5 | | 1.4727 | 32937000 | 4567200 | 93840000 | 27262000 | 81089000 | 35323000 |
| F6USD5;F6T4L3;Q80ZP8;Q3T | Manf | Mesencephalic astrocyte-derived neurotrophic factor | | 1.4758 | 440390 | 224840 | 1147700 | 1861400 | 870040 | 1079400 |
| Q9DCT8;A0A0G2JF37 | Crip2 | Cysteine-rich protein 2 | | 1.4828 | 1025300 | 778860 | 1727600 | 3062000 | 2216900 | 1828800 |
| P70698 | Ctps1 | CTP synthase 1 | | 1.4939 | 335980 | 343550 | 447370 | 1427900 | 1836700 | 1216100 |
| P26883;F6X9I3 | Fkbp1a | -prolyl cis-trans isomerase FKBP1A;Peptidyl-prolyl cis-trans isomerase | | 1.5196 | 5294700 | 10230000 | 5223500 | 7235300 | 0 | 0 |
| P21107-2;D3Z2H9;E9Q5J9 | Tpm3;Tpm3-rs7 | Tropomyosin alpha-3 chain | | 1.561 | 37153000 | 31194000 | 65480000 | 96097000 | 138150000 | 74394000 |
| Q8K2B3;A0A1Y7VJ55 | Sdha | ate dehydrogenase [ubiquinone] flavoprotein subunit, mitochondrial | | 1.5943 | 560950 | 312230 | 688500 | 1187500 | 1380000 | 1831800 |
| CON__P12763 | | | + | 1.61 | 2765800 | 2487800 | 245280 | 447180 | 4445100 | 5555000 |
| F7DBQ0;Q3TML0;Q922R8 | Pdia6 | Protein disulfide-isomerase A6 | | 1.6198 | 872300 | 694470 | 2329300 | 4497200 | 2856600 | 2638800 |
| G3X924;Q91VC4 | Plvap | Plasmalemma vesicle-associated protein | | 1.6381 | 977440 | 1844300 | 0 | 0 | 4171000 | 2871400 |
| A0A338P6J5;A0A338P6S7;Q9 | Ppil2 | Peptidyl-prolyl cis-trans isomerase-like 2 | | 1.6391 | 0 | 0 | 1649700 | 2403200 | 1192900 | 889620 |
| P16045;A0A2R8VHJ0 | Lgals1 | Galectin-1 | | 1.6602 | 0 | 0 | 0 | 0 | 5262700 | 4607700 |
| E9PWE8;Q3TT92;Q62188 | Dpysl3 | Dihydropyrimidinase-related protein 3 | | 1.6925 | 523400 | 973960 | 3158700 | 3323200 | 3474100 | 5402100 |
| Q60668-3;Q60668;E9Q5B6;F | Hnrnpd | Heterogeneous nuclear ribonucleoprotein D0 | | 1.7138 | 1183800 | 525030 | 2180600 | 3479800 | 1548800 | 1464600 |
| Q9R0P9 | Uchl1 | Ubiquitin carboxyl-terminal hydrolase isozyme L1 | | 1.7193 | 4190500 | 2258600 | 1404800 | 2509100 | 9903800 | 5802600 |
| Q9WVK4 | Ehd1 | EH domain-containing protein 1 | | 1.7544 | 0 | 0 | 1489200 | 2380700 | 0 | 0 |
| E9Q616;A0A494BBD5;G5E8K | Ahnak | | | 1.7751 | 1754500 | 1640800 | 2684300 | 4327800 | 0 | 0 |
| B8JK33;B8JK32;Q9D0E1-2;Q9 | Hnrnpm | Heterogeneous nuclear ribonucleoprotein M | | 1.8003 | 5771800 | 3510800 | 10538000 | 16099000 | 5491700 | 10276000 |
| Q9WTI7;Q9WTI7-4;Q9WTI7- | Myo1c | Unconventional myosin-Ic | | 1.8579 | 1668600 | 1507100 | 3834100 | 11601000 | 6773500 | 5302700 |
| CON__P13645;A2A513;CON_ | Krt10 | Keratin, type I cytoskeletal 10 | + | 1.8772 | 0 | 0 | 1337200 | 2331700 | 610330 | 1389700 |
| P24549;O35945;A0A286YCZ( | Aldh1a1;Aldh1a7 | Retinal dehydrogenase 1;Aldehyde dehydrogenase, cytosolic 1 | | 1.8986 | 96459 | 1189100 | 0 | 0 | 4374600 | 5508700 |
| A0A1L1STC6;Q6ZWR6-4;Q6Z | Syne1 | Nesprin-1 | | 1.9996 | 0 | 0 | 3622900 | 6540500 | 10605000 | 17163000 |
| CON__P02768-1 | | | + | 2.205 | 0 | 0 | 466280 | 1997700 | 13179000 | 10434000 |

| | | | | | | | | |
|---|---|---|---|---|---|---|---|---|
| F7A1B4;Q3UAM9;Q63961;F7 | Eng | Endoglin | 2.2681 | 0 | 0 | 2175500 | 4865800 | 0 | 0 |
| Q8R3C7;Q8BZY3;Q61655 | Ddx19b;Ddx19a | ATP-dependent RNA helicase DDX19A | 2.2734 | 1562400 | 3784000 | 2892400 | 6804700 | 0 | 0 |
| Q6NSP9;P52927 | Hmga2 | High mobility group protein HMGI-C | 2.3683 | 0 | 0 | 196460 | 8751100 | 4878100 | 1130600 |
| Q9Z0M3 | Cdh20 | Cadherin-20 | 2.57 | 1614600 | 1682100 | 3850100 | 8288000 | 0 | 0 |
| H3BKG0;P49817 | Cav1 | Caveolin-1 | 2.6385 | 1580600 | 3059200 | 4354700 | 9647900 | 0 | 0 |
| B1AYB7;B1AYB5;B1AYB6 | Mbd5 | Methyl-CpG-binding domain protein 5 | 2.6677 | 5357400 | 3827500 | 14351000 | 29553000 | 6408600 | 9687100 |
| Q99N15;A2AFQ2;O08756 | Hsd17b10 | 3-hydroxyacyl-CoA dehydrogenase type-2 | 2.8122 | 460530 | 1487100 | 1540100 | 3447100 | 0 | 0 |
| P67778;Q5SQG5 | Phb | Prohibitin | 3.9236 | 3110100 | 19673000 | 9143900 | 31749000 | 6123500 | 23395000 |
| P14069 | S100a6 | Protein S100-A6 | 3.9588 | 2822400 | 8325600 | 3620300 | 15408000 | 5711000 | 6247800 |
| P62309 | Snrpg | Small nuclear ribonucleoprotein G | 4.0688 | 0 | 0 | 675050 | 1347000 | 399450 | 1514900 |
| B1ARB3;Q08481-4;Q08481-3 | Pecam1 | Platelet endothelial cell adhesion molecule | 6.5046 | 472080 | 1639400 | 1028700 | 4844200 | 0 | 0 |
| A0A0A6YWA9;P08752 | Gnai2 | Guanine nucleotide-binding protein G(i) subunit alpha-2 | 11.818 | 0 | 0 | 1627700 | 17310000 | 891660 | 11549000 |
| E9Q390;A0A286YDF5;Q69ZN | Myof | Myoferlin | 16.125 | 0 | 0 | 2302600 | 15581000 | 1886700 | 1104500 |
| O70435;E0CZ34;F8WH02;E0( | Psma3 | Proteasome subunit alpha type-3 | 20.819 | 531440 | 6810500 | 1725100 | 21355000 | 0 | 0 |
| Q8BH97;A0A1B0GS22;A0A1I | Rcn3 | Reticulocalbin-3 | 21.702 | 0 | 1782100 | 144790 | 7085400 | 884890 | 13919000 |
| Q792F9;Q00651 | Itga4 | Integrin alpha-4 | 30.998 | 3109000 | 25302000 | 2033100 | 46864000 | 2320500 | 26802000 |
| Q3V116;E0CXV8;Q7TT35;Q64 | Phc1 | Polyhomeotic-like protein 1 | 49.2 | 188190 | 5966600 | 947170 | 60844000 | 567780 | 14699000 |
| A0A087WNK8;P61375;P6300 | Lhx1;Lhx5 | LIM/homeobox protein Lhx5;LIM/homeobox protein Lhx1 | NaN | 0 | 0 | 0 | 0 | 1224900 | 784860 |
| A0A087WNP6;Q4VAA2-2;Q4 | Cdv3 | Protein CDV3 | NaN | 0 | 0 | 0 | 0 | 0 | 0 |
| A0A087WNS0;A0A087WQK0 | Rpl3 | 60S ribosomal protein L3 | NaN | 0 | 0 | 0 | 0 | 0 | 1332700 |
| A0A087WQE6;A0A087WNT1 | Tceb1 | Transcription elongation factor B polypeptide 1 | NaN | 0 | 0 | 0 | 0 | 0 | 0 |
| A0A087WQF8;A0A087WP48 | Ktn1 | Kinectin | NaN | 0 | 0 | 0 | 0 | 0 | 0 |
| A0A087WNY2;A0A087WRT0 | Atg4b | Cysteine protease ATG4B | NaN | 0 | 0 | 0 | 0 | 0 | 0 |
| A0A087WNZ7;G5E870;A0A0 | Trip12 | E3 ubiquitin-protein ligase TRIP12 | NaN | 0 | 0 | 0 | 0 | 0 | 0 |
| A0A087WP00;A0A0A0MQ76 | Nop58 | Nucleolar protein 58 | NaN | 0 | 0 | 0 | 0 | 0 | 0 |
| E0CY18;A0A087WPF5;Q91X5 | Zfand2b | AN1-type zinc finger protein 2B | NaN | 0 | 0 | 0 | 0 | 0 | 0 |
| A0A087WPL5;E9QNN1;O701 | Dhx9 | ATP-dependent RNA helicase A | NaN | 0 | 0 | 0 | 0 | 0 | 0 |
| A0A087WRU0;A0A087WQ94 | Tns1 | | NaN | 0 | 0 | 0 | 0 | 0 | 0 |
| E9QAN4;G3UW47;Q6TA13;E | Kif1a;Kif1b | sin-like protein;Kinesin-like protein KIF1A;Kinesin-like protein KIF1B | NaN | 0 | 0 | 0 | 0 | 0 | 0 |
| A0A087WQM1;Q07235;A0A( | Serpine2 | Glia-derived nexin | NaN | 0 | 0 | 722960 | 457220 | 0 | 0 |
| A0A087WQQ3;A0A087WRE9 | Rnf2 | E3 ubiquitin-protein ligase RING2 | NaN | 0 | 0 | 0 | 0 | 0 | 0 |
| A0A087WR70;A0A087WQS1 | Sgk3 | Serine/threonine-protein kinase Sgk3 | NaN | 0 | 0 | 0 | 0 | 0 | 0 |
| A0A087WR97;Q921F2 | Tardbp | TAR DNA-binding protein 43 | NaN | 0 | 0 | 0 | 0 | 0 | 0 |
| A0A087WRM2;Q91VK4 | Itm2c | Integral membrane protein 2C;CT-BRI3 | NaN | 0 | 0 | 0 | 0 | 0 | 0 |
| A0A087WS46;O70251 | Eef1b2;Eef1b | Elongation factor 1-beta | NaN | 0 | 0 | 1049700 | 826430 | 0 | 0 |
| A0A0A0MQA5;P68368;Q9JJZ | Tuba4a;Tuba8 | Tubulin alpha-4A chain;Tubulin alpha-8 chain | NaN | 0 | 0 | 0 | 0 | 0 | 0 |
| A0A0A0MQF5;Q8K2H2 | Otud6b | OTU domain-containing protein 6B | NaN | 0 | 0 | 0 | 0 | 0 | 0 |
| A0A0A0MQM0;P63242;J3QP | Eif5a | translation initiation factor 5A;Eukaryotic translation initiation factor 5A-1 | NaN | 0 | 0 | 967990 | 603800 | 0 | 0 |
| A0A0A6YVU8;Q9JKV1 | Gm9774;Adrm1 | Proteasomal ubiquitin receptor ADRM1 | NaN | 0 | 0 | 0 | 0 | 0 | 0 |
| B7ZNR9;B2RUE8;A0A0A6YW | Map4k4 | Mitogen-activated protein kinase kinase kinase kinase 4 | NaN | 0 | 0 | 0 | 0 | 0 | 0 |
| A0A0A6YX73;Q8K1M3 | Prkar2a | | NaN | 0 | 0 | 0 | 0 | 0 | 0 |
| A0A0A6YXG0;Q9JKK1-2;Q9JK | Stx6 | Syntaxin-6 | NaN | 0 | 0 | 0 | 0 | 0 | 0 |
| A0A0A6YXS4;A0A0A6YY72;P | Impdh2 | Inosine-5-monophosphate dehydrogenase 2 | NaN | 0 | 0 | 0 | 0 | 0 | 0 |
| A0A0A6YXV0;A0A0R4J107;Q | Apeh | Acylamino-acid-releasing enzyme | NaN | 0 | 0 | 0 | 0 | 0 | 0 |
| A0A0A6YY39;A2AJ72;Q3TIX6 | Fubp3 | | NaN | 0 | 0 | 73890 | 4800400 | 0 | 0 |

| | | | | | | | | | |
|---|---|---|---|---|---|---|---|---|---|
| A0A0B4J1E2;Q9CSN1 | Snw1 | SNW domain-containing protein 1 | NaN | 0 | 0 | 0 | 0 | 0 | 0 |
| A0A0B4J1J2 | Igkv5-43 | | NaN | 0 | 0 | 0 | 0 | 0 | 0 |
| A0A0G2JEF6;A0A0G2JGZ1;A | Lrrfip2 | | NaN | 0 | 0 | 0 | 0 | 0 | 0 |
| A0A0H2UH17;A0A0G2JDV6;( | Ubap2l | Ubiquitin-associated protein 2-like | NaN | 0 | 0 | 0 | 0 | 0 | 0 |
| A0A0G2JE27;Q5RKN9;P4775 | Capza1 | F-actin-capping protein subunit alpha-1 | NaN | 0 | 0 | 0 | 0 | 3596500 | 2323100 |
| A0A0G2JEC4;Q9JK48-3;Q9JK | Sh3glb1 | Endophilin-B1 | NaN | 0 | 0 | 0 | 0 | 0 | 0 |
| A0A0G2JEF4;A0A0G2JFG5;Q | Camkk2 | Calcium/calmodulin-dependent protein kinase kinase 2 | NaN | 0 | 0 | 0 | 0 | 0 | 0 |
| A0A0H2UH27;A0A0G2JEP0;C | Fxr1 | Fragile X mental retardation syndrome-related protein 1 | NaN | 0 | 0 | 0 | 0 | 0 | 0 |
| A0A0G2JEU1;P47738;D3YYF3 | Aldh2 | Aldehyde dehydrogenase, mitochondrial | NaN | 0 | 0 | 0 | 0 | 0 | 0 |
| A0A0G2JFX7;Q9CWZ3;A0A0( | Rbm8a | RNA-binding protein 8A | NaN | 0 | 0 | 0 | 0 | 0 | 0 |
| A0A0G2JG00;Q3TUE1;A0A0( | Fubp1 | Far upstream element-binding protein 1 | NaN | 0 | 0 | 684940 | 4264500 | 0 | 0 |
| A0A2R8VHB7;A0A2R8VI28;A | Smarcd3;Smarcd2;Smarcd1 | natrix-associated actin-dependent regulator of chromatin subfamily D member 2;SWI | NaN | 0 | 0 | 0 | 0 | 0 | 0 |
| A0A0J9YUG2;P62141 | Ppp1cb | erine/threonine-protein phosphatase PP1-beta catalytic subunit | NaN | 0 | 0 | 0 | 0 | 0 | 0 |
| A0A0J9YUS5;E9PVC6;E9Q9E1 | Eif4g1 | Eukaryotic translation initiation factor 4 gamma 1 | NaN | 0 | 0 | 0 | 0 | 11981000 | 1270000 |
| A0A0J9YUT8;Q8R3C6 | Rbm19 | Probable RNA-binding protein 19 | NaN | 0 | 0 | 18949000 | 1657200 | 0 | 0 |
| A0A0J9YVG0;Q61074 | Ppm1g | Protein phosphatase 1G | NaN | 0 | 0 | 0 | 0 | 2609000 | 912800 |
| E9QL12;E9PXU9;A0A0N4SUJ | Dysf | Dysferlin | NaN | 0 | 0 | 0 | 0 | 0 | 0 |
| A0A0N4SV00;P80313;A0A0N | Cct7 | T-complex protein 1 subunit eta | NaN | 0 | 0 | 0 | 0 | 0 | 0 |
| A0A0N4SV15;Q80UY2-2;Q80 | Kcmf1 | E3 ubiquitin-protein ligase KCMF1 | NaN | 0 | 0 | 0 | 0 | 0 | 0 |
| A0A0N4SVC2;E9QP00;Q6PFF | Tra2a | Transformer-2 protein homolog alpha | NaN | 0 | 0 | 0 | 0 | 0 | 0 |
| A2AMI7;Z4YKC4;A0A0R4J112 | Eif4g3 | Eukaryotic translation initiation factor 4 gamma 3 | NaN | 0 | 0 | 0 | 0 | 0 | 0 |
| A0A0N4SVS6;P53996-2;P539 | Cnbp | Cellular nucleic acid-binding protein | NaN | 0 | 0 | 0 | 0 | 0 | 0 |
| A0A0N4SW65;A0A0N4SWC3 | Cmas | N-acylneuraminate cytidylyltransferase | NaN | 0 | 0 | 0 | 0 | 0 | 0 |
| A0A0N4SW94;O35682 | Myadm | Myeloid-associated differentiation marker | NaN | 0 | 0 | 0 | 0 | 0 | 0 |
| A0A0R3P9C8;Q9DC69 | Ndufa9 | hydrogenase [ubiquinone] 1 alpha subcomplex subunit 9, mitochondrial | NaN | 0 | 0 | 0 | 0 | 0 | 0 |
| A0A0R4IZW8;O88456;A0A0F | Capns1 | Calpain small subunit 1 | NaN | 0 | 0 | 0 | 0 | 0 | 0 |
| A0A0R4IZW9;Q9DCT5 | Sdf2 | Stromal cell-derived factor 2 | NaN | 0 | 0 | 0 | 0 | 0 | 0 |
| Q6NS54;A0A0R4IZX2;Q9JHL1 | Slc9a3r2 | ge regulatory cofactor NHE-RF;Na(+)/H(+) exchange regulatory cofactor NHE-RF2 | NaN | 0 | 0 | 0 | 0 | 0 | 0 |
| A0A0R4IZY9;Q91XI1-2;Q91XI | Dus3l | tRNA-dihydrouridine(47) synthase [NAD(P)(+)]-like | NaN | 0 | 0 | 0 | 0 | 0 | 0 |
| A0A0R4J047;Q9CYI4;D6RGP6 | Luc7l | Putative RNA-binding protein Luc7-like 1 | NaN | 0 | 0 | 1947400 | 1002300 | 0 | 0 |
| A0A0R4J078;Q8VCH8 | Ubxn4 | UBX domain-containing protein 4 | NaN | 0 | 0 | 0 | 0 | 0 | 0 |
| A0A0R4J079;Q8BMP6 | Acbd3 | Golgi resident protein GCP60 | NaN | 0 | 0 | 0 | 0 | 0 | 0 |
| A0A0R4J080;Q62130 | Ptpn14 | Tyrosine-protein phosphatase non-receptor type 14 | NaN | 0 | 0 | 0 | 0 | 3254000 | 0 |
| A0A0R4J083;P51174 | Acadl | Long-chain specific acyl-CoA dehydrogenase, mitochondrial | NaN | 0 | 0 | 956100 | 269870 | 0 | 0 |
| A0A0R4J0A0;F6W2Q5;Q5JC2 | Eps15 | Epidermal growth factor receptor substrate 15 | NaN | 0 | 0 | 0 | 0 | 0 | 0 |
| A0A0R4J0E4 | | | NaN | 0 | 0 | 0 | 0 | 0 | 0 |
| A0A0R4J0I4;Q8BIW9 | Chtf18 | Chromosome transmission fidelity protein 18 homolog | NaN | 0 | 0 | 11860000 | 7363600 | 0 | 0 |
| A0A0R4J0M9;Q8CDG3 | Vcpip1 | Deubiquitinating protein VCIP135 | NaN | 0 | 0 | 0 | 0 | 0 | 0 |
| A0A0R4J0Q5;P21619-2;P216 | Lmnb2 | Lamin-B2 | NaN | 0 | 0 | 0 | 0 | 0 | 0 |
| A0A0R4J0R1;O70404 | Vamp8 | Vesicle-associated membrane protein 8 | NaN | 0 | 0 | 0 | 0 | 0 | 0 |
| A0A0R4J0S1;Q91W92 | Cdc42ep1 | Cdc42 effector protein 1 | NaN | 0 | 0 | 0 | 0 | 0 | 0 |
| A0A2I3BQY0;A0A0R4J0T8;Q( | Arfgap3 | ADP-ribosylation factor GTPase-activating protein 3 | NaN | 0 | 0 | 0 | 0 | 0 | 0 |
| F6TQN9;E9QL31;A0A0R4J10 | Dab2 | Disabled homolog 2 | NaN | 0 | 0 | 3391100 | 2568500 | 0 | 0 |
| A0A0R4J138;P50429 | Arsb | Arylsulfatase B | NaN | 0 | 0 | 0 | 0 | 0 | 0 |
| A0A0R4J1E3;Q9QXS6-3;Q9Q | Dbn1 | Drebrin | NaN | 0 | 0 | 0 | 0 | 0 | 0 |

| | | | | | | | | | |
|---|---|---|---|---|---|---|---|---|---|
| D3Z0Q8;A0A0R4J1H6;E9QP9 | Golga3 | Golgin subfamily A member 3 | NaN | 0 | 0 | 0 | 0 | 0 | 0 |
| A0A0R4J1L2;F6ZFU0;P57776 | Eef1d | Elongation factor 1-delta | NaN | 0 | 0 | 0 | 0 | 0 | 0 |
| D6RE33;A0A0R4J1Q0;G5E89 | Edc4 | Enhancer of mRNA-decapping protein 4 | NaN | 0 | 0 | 0 | 0 | 0 | 0 |
| A0A0R4J1W0;B2RXQ9;D3Z0 | Sorbs2 | Sorbin and SH3 domain-containing protein 2 | NaN | 0 | 0 | 0 | 0 | 0 | 0 |
| A0A0R4J1Y7;E9PXX7;Q91W9 | Txndc5 | Thioredoxin domain-containing protein 5 | NaN | 0 | 0 | 0 | 0 | 0 | 0 |
| A0A0R4J275;Q7TMF3 | Ndufa12 | ADH dehydrogenase [ubiquinone] 1 alpha subcomplex subunit 12 | NaN | 0 | 0 | 0 | 0 | 0 | 0 |
| A0A0U1RNJ1;P19096 | Fasn | er-protein] synthase;3-oxoacyl-[acyl-carrier-protein] reductase;3-hydroxyacyl-[acyl-car | NaN | 0 | 0 | 2529900 | 4710200 | 0 | 0 |
| A0A0U1RNT6;A0A0U1RNK6; | Mat2a | S-adenosylmethionine synthase isoform type-2 | NaN | 4403500 | 846910 | 0 | 0 | 0 | 0 |
| A0A0U1RNM2;A1L3S7;Q8VH | Gatad2b | Transcriptional repressor p66-beta | NaN | 0 | 0 | 0 | 0 | 0 | 0 |
| A0A0U1RP97;P06745;A0A0U | Gpi | Glucose-6-phosphate isomerase | NaN | 0 | 0 | 0 | 0 | 0 | 0 |
| Q9R166;A0A0U1RPC5 | Zfp109 | | NaN | 0 | 0 | 0 | 0 | 13674000 | 1024000 |
| D3YZ84;D3YY06;D3YWG5;D6 | Tsen34 | tRNA-splicing endonuclease subunit Sen34 | NaN | 0 | 0 | 0 | 0 | 0 | 0 |
| A0A140LHW5;A0A140LIK0;A | Spcs2 | Signal peptidase complex subunit 2 | NaN | 0 | 0 | 0 | 0 | 0 | 0 |
| A0A140LHU9;A0A140LI54;Q | Pnpla6 | Neuropathy target esterase | NaN | 0 | 0 | 0 | 0 | 0 | 0 |
| D3Z6W1;A0A140LIT9;Q7TSJ | Map6 | Microtubule-associated protein 6 | NaN | 0 | 0 | 0 | 0 | 0 | 0 |
| A0A140LJ36;Q9JIG8 | Praf2 | PRA1 family protein 2 | NaN | 0 | 0 | 0 | 0 | 0 | 0 |
| A0A171EBL2;E9Q555 | Rnf213 | E3 ubiquitin-protein ligase RNF213 | NaN | 0 | 0 | 0 | 0 | 161930000 | 0 |
| A0A1B0GQZ1;A0A1B0GSZ9; | Mrpl23 | 39S ribosomal protein L23, mitochondrial | NaN | 0 | 0 | 0 | 0 | 0 | 0 |
| A0A1B0GR08;Q921G8 | Tubgcp2 | Gamma-tubulin complex component 2 | NaN | 0 | 0 | 0 | 0 | 0 | 0 |
| A0A1B0GS13;A0A1B0GT81; | Bax | Apoptosis regulator BAX | NaN | 0 | 0 | 0 | 0 | 0 | 0 |
| Q8K4L2;E9Q3Z5;A0A1B0GS9 | Svil | Supervillin | NaN | 0 | 0 | 0 | 0 | 0 | 0 |
| D3Z0B9;A0A1B0GSU0;Q571 | Aldh16a1 | Aldehyde dehydrogenase family 16 member A1 | NaN | 0 | 0 | 0 | 0 | 0 | 0 |
| A0A1B0GSX6;O70209 | Pdlim3 | PDZ and LIM domain protein 3 | NaN | 0 | 0 | 0 | 0 | 0 | 0 |
| A0A1B0GSY1;Q9ESW8 | Pgpep1 | Pyroglutamyl-peptidase 1 | NaN | 0 | 0 | 0 | 0 | 0 | 0 |
| A0A1B0GX15;G3UWN5;P082 | Apoe | Apolipoprotein E | NaN | 0 | 0 | 0 | 0 | 0 | 0 |
| D6RCH8;F8WHJ1;A0A1C7CYU | Fam160a2 | FTS and Hook-interacting protein | NaN | 0 | 0 | 0 | 0 | 0 | 0 |
| E9QA74;E9Q405;A0A1C7ZN1 | Myo18a | Unconventional myosin-XVIIIa | NaN | 0 | 0 | 0 | 0 | 0 | 0 |
| A0A1D5RM74;A0A1D5RLE4; | Csnk2a2 | Casein kinase II subunit alpha | NaN | 0 | 0 | 0 | 0 | 0 | 0 |
| A0A1D5RLU9;A0A1D5RLE8;A | Cmtm3 | CKLF-like MARVEL transmembrane domain-containing protein 3 | NaN | 0 | 0 | 0 | 0 | 0 | 0 |
| A0A1D5RLF2;A0A1D5RLI3;A | Fam192a | Protein FAM192A | NaN | 0 | 0 | 0 | 0 | 0 | 0 |
| A0A1D5RLS1;Q60902-3;Q60 | Eps15l1 | Epidermal growth factor receptor substrate 15-like 1 | NaN | 0 | 0 | 0 | 0 | 0 | 0 |
| A0A1D5RLW5;A0A1D5RM85 | Rpl18a | 60S ribosomal protein L18a | NaN | 0 | 0 | 2578700 | 1907900 | 0 | 0 |
| A0A1D5RLY6;Q8C052 | Map1s | otubule-associated protein 1S;MAP1S heavy chain;MAP1S light chain | NaN | 0 | 0 | 0 | 0 | 0 | 0 |
| A0A1D5RLZ6;Q9DCC8 | Tomm20 | Mitochondrial import receptor subunit TOM20 homolog | NaN | 0 | 0 | 0 | 0 | 0 | 0 |
| A0A1D5RMI0;F8WHV1;Q9D1 | Fam96b | Mitotic spindle-associated MMXD complex subunit MIP18 | NaN | 0 | 0 | 0 | 0 | 0 | 0 |
| A0A1L1SR69;A0A1L1ST61;A | Higd1a | HIG1 domain family member 1A, mitochondrial | NaN | 0 | 0 | 0 | 0 | 0 | 0 |
| A0A1L1SRS6;D3YV69;F7BE3 | 4;Rab6b;Rab39a;Rab27b;Rab27a;Rasef;Cr | ted protein Rab-27A;Ras-related protein Rab-34;Ras and EF-hand domain-containing | NaN | 0 | 0 | 0 | 0 | 0 | 0 |
| A0A1L1SS87;A0A1L1SU40 | | | NaN | 0 | 0 | 17137000 | 13594000 | 0 | 0 |
| J3QN87;A0A1L1SSA3;Q9CXU | Eif1;Eif1b | c translation initiation factor 1b;Eukaryotic translation initiation factor 1 | NaN | 0 | 0 | 0 | 0 | 0 | 0 |
| A0A1L1SSA8;Q91XE8 | Tmem205 | Transmembrane protein 205 | NaN | 0 | 0 | 0 | 0 | 0 | 0 |
| Q3UDS7;A0A1L1SSF2;Q8VDL | Adpgk | ADP-dependent glucokinase | NaN | 0 | 0 | 416520 | 506060 | 0 | 0 |
| A0A1L1STC0;Q61081 | Cdc37 | o-chaperone Cdc37;Hsp90 co-chaperone Cdc37, N-terminally processed | NaN | 0 | 0 | 0 | 0 | 0 | 0 |
| Q45VK5;A0A1L1STE4;Q9Z1X | Ilf3 | Interleukin enhancer-binding factor 3 | NaN | 0 | 0 | 0 | 0 | 0 | 0 |
| A0A1L1STF0;Q6PDI6-3;Q6PD | Fam63b | Protein FAM63B | NaN | 0 | 0 | 0 | 0 | 0 | 0 |
| A0A1L1STY4;A0A1L1SVG0;Q | Uaca | Uveal autoantigen with coiled-coil domains and ankyrin repeats | NaN | 4246200 | 0 | 0 | 0 | 0 | 0 |

| | | | | | | | | | |
|---|---|---|---|---|---|---|---|---|---|
| A0A1L1SVK0;Q61206;A0A1L1 | Pafah1b2 | Platelet-activating factor acetylhydrolase IB subunit beta | NaN | 0 | 0 | 0 | 0 | 0 | 0 |
| A0A1W2P6H2;A0A1W2P7I2;( | Epb4.1l2;Epb41l2 | Band 4.1-like protein 2 | NaN | 0 | 0 | 0 | 0 | 0 | 0 |
| A0A1W2P7I5;A0A1W2P6P1;( | Sgta | ll glutamine-rich tetratricopeptide repeat-containing protein alpha | NaN | 0 | 0 | 0 | 0 | 0 | 0 |
| A0A1W2P6Y0;Q6P8X1 | Snx6 | Sorting nexin-6;Sorting nexin-6, N-terminally processed | NaN | 0 | 0 | 0 | 0 | 0 | 0 |
| A0A1W2P8D6;A0A1W2P727; | Ilvbl | Acetolactate synthase-like protein | NaN | 0 | 0 | 0 | 0 | 0 | 0 |
| A0A1W2P729;Q9QXT0 | Cnpy2 | Protein canopy homolog 2 | NaN | 0 | 0 | 0 | 0 | 0 | 0 |
| A0A1W2P733;Q922D8 | Mthfd1 | se;Methenyltetrahydrofolate cyclohydrolase;Formyltetrahydrofolate synthetase;C-1-te | NaN | 0 | 0 | 0 | 0 | 0 | 0 |
| A0A1W2P7H2;P62075 | Timm13 | litochondrial import inner membrane translocase subunit Tim13 | NaN | 0 | 0 | 0 | 0 | 0 | 0 |
| A0A1W2P7Z8;A0A1W2P7H4; | Cdc5l | Cell division cycle 5-like protein | NaN | 0 | 0 | 0 | 0 | 0 | 0 |
| A0A1W2P7Q6;Q9CRT8 | Xpot | Exportin-T | NaN | 0 | 0 | 0 | 0 | 0 | 0 |
| A0A1W2P7X0;E9QMV2;Q4KI | Abracl | Costars family protein ABRACL | NaN | 0 | 0 | 0 | 0 | 0 | 0 |
| A0A1W2P7X3;Q9D358;Q561 | Acp1 | Low molecular weight phosphotyrosine protein phosphatase | NaN | 0 | 0 | 0 | 0 | 0 | 0 |
| B2RRF0;B1AQN2;A0A1W2P7 | Ptprk;Ptprt | eceptor-type tyrosine-protein phosphatase T;Receptor-type tyrosine-protein phosphata | NaN | 0 | 0 | 2278100 | 1222400 | 0 | 0 |
| A0A1Y7VJ71;P45591 | Cfl2 | Cofilin-2 | NaN | 0 | 0 | 0 | 0 | 2304600 | 1701900 |
| A0A1Y7VM39;Q3TZZ7 | Esyt2 | Extended synaptotagmin-2 | NaN | 0 | 0 | 0 | 0 | 0 | 0 |
| A0A1Y7VM80;E9QLA5;Q0GN | Inf2 | Inverted formin-2 | NaN | 0 | 0 | 0 | 0 | 0 | 0 |
| F8VPK5;A0A1Y7VMN0;P7033 | Rock2 | Rho-associated protein kinase;Rho-associated protein kinase 2 | NaN | 4287600 | 0 | 0 | 0 | 1450600 | 1599100 |
| A0A286YDB7;A0A286YCT4;A | Ssr1 | Translocon-associated protein subunit alpha | NaN | 0 | 0 | 0 | 0 | 0 | 0 |
| A0A286YD22;A0A286YDT9;A | Gkap1 | G kinase-anchoring protein 1 | NaN | 0 | 0 | 0 | 0 | 0 | 0 |
| A0A286YD68;Q8JZR0 | Acsl5 | Long-chain-fatty-acid--CoA ligase 5 | NaN | 0 | 0 | 0 | 0 | 0 | 0 |
| A0A286YE75;Q59J78 | Ndufaf2 | Mimitin, mitochondrial | NaN | 0 | 0 | 0 | 0 | 0 | 0 |
| A0A2C9F2D2;Q07076;A0A28 | Anxa7 | Annexin A7 | NaN | 0 | 0 | 0 | 0 | 0 | 0 |
| A0A2I3BR03;A0A2I3BPT1;A0 | App | oid protein 40;C83;P3(42);P3(40);C80;Gamma-secretase C-terminal fragment 59;Gam | NaN | 0 | 0 | 0 | 0 | 0 | 0 |
| A0A2I3BQ39;Q9JKB1;P58321 | Uchl3;Uchl4 | -terminal hydrolase isozyme L3;Ubiquitin carboxyl-terminal hydrolase isozyme L4 | NaN | 0 | 0 | 1695900 | 1773900 | 0 | 0 |
| A0A2I3BQF4;P62889 | Rpl30 | 60S ribosomal protein L30 | NaN | 0 | 0 | 0 | 0 | 0 | 0 |
| A0A2I3BQJ3;Q8K072 | Reep4 | Receptor expression-enhancing protein 4 | NaN | 0 | 0 | 0 | 0 | 0 | 0 |
| A0A2I3BQZ0;Q7TN22-2;Q7TI | Txndc16 | Thioredoxin domain-containing protein 16 | NaN | 0 | 0 | 0 | 0 | 0 | 0 |
| A0A2R8VI25;Q3TT81;A0A2R8 | Pcbp2;Pcbp3 | Poly(rC)-binding protein 2;Poly(rC)-binding protein 3 | NaN | 0 | 0 | 2441700 | 2187700 | 0 | 0 |
| A0A2R8VK76;Q9EPU4 | Cpsf1 | Cleavage and polyadenylation specificity factor subunit 1 | NaN | 0 | 0 | 0 | 0 | 0 | 0 |
| A0A2R8W6Y5;A0A2R8VKL5;I | Larp4 | La-related protein 4 | NaN | 0 | 0 | 0 | 0 | 0 | 0 |
| A0A2U3TZ82;Q99KW3-1;Q9 | Triobp | TRIO and F-actin-binding protein | NaN | 0 | 0 | 973380 | 1996000 | 0 | 0 |
| A0A338P675;P47941 | Crkl | Crk-like protein | NaN | 0 | 0 | 0 | 0 | 0 | 0 |
| A0A338P7E5;A0A338P786;P6 | Ube2l3 | Ubiquitin-conjugating enzyme E2 L3 | NaN | 0 | 0 | 0 | 0 | 0 | 0 |
| A0A338P7D7;Q9Z2U0;B7ZM | Psma7;Psma8 | alpha type-7;Proteasome subunit alpha type;Proteasome subunit alpha type-7-like | NaN | 0 | 0 | 0 | 0 | 0 | 0 |
| A0A338P7L3;H3BLR8;I1E4X7; | Nudt3 | Diphosphoinositol polyphosphate phosphohydrolase 1 | NaN | 0 | 0 | 0 | 0 | 0 | 0 |
| A0A338P7L9;Q9CQ49 | Ncbp2 | Nuclear cap-binding protein subunit 2 | NaN | 0 | 0 | 0 | 0 | 0 | 0 |
| A0A3B2W7I6;A0A3B2W883; | Srpk1 | SRSF protein kinase 1 | NaN | 0 | 0 | 0 | 0 | 0 | 0 |
| A0A3B2WBC6 | | | NaN | 0 | 0 | 0 | 0 | 0 | 0 |
| A0A3B2WBH9;A0A3B2WCN | Tjp2 | Tight junction protein ZO-2 | NaN | 0 | 0 | 0 | 0 | 0 | 0 |
| A0A3B2WCL5;Q9CT10 | Ranbp3 | Ran-binding protein 3 | NaN | 0 | 0 | 0 | 0 | 0 | 0 |
| A0A452J8C7;Q8K4Q8 | Colec12 | Collectin-12 | NaN | 0 | 0 | 0 | 0 | 0 | 0 |
| A0A494B8X7;A0A494B9A2;B | Asah2 | Neutral ceramidase;Neutral ceramidase soluble form | NaN | 0 | 0 | 0 | 0 | 0 | 0 |
| A0A494BA39;A0A494B9F0;A | Nedd4l | E3 ubiquitin-protein ligase NEDD4-like | NaN | 0 | 0 | 0 | 0 | 0 | 0 |
| G3UXW2;A0A494BA33;A0A4 | H2-K1;H2-D1 | hain;H-2 class I histocompatibility antigen, K-K alpha chain;H-2 class I histocompatibili | NaN | 0 | 0 | 0 | 0 | 0 | 0 |
| A0A494BA44;Q04207-2;Q04 | Rela | Transcription factor p65 | NaN | 0 | 0 | 0 | 0 | 0 | 0 |

| | | | | | | | | | |
|---|---|---|---|---|---|---|---|---|---|
| A0A494BA52 | | | NaN | 0 | 0 | 0 | 0 | 0 | 0 |
| A0A494BAY0;A0A494BAR3;( | Vps37c | Vacuolar protein sorting-associated protein 37C | NaN | 0 | 0 | 0 | 0 | 0 | 0 |
| A0A494BB04;Q5FWI3 | Tmem2 | Transmembrane protein 2 | NaN | 0 | 0 | 0 | 0 | 0 | 0 |
| A0A494BB86;P61164 | Actr1a | Alpha-centractin | NaN | 0 | 0 | 0 | 0 | 0 | 0 |
| A0A494BBM6;R4H4V1;A0A4 | Scyl1 | N-terminal kinase-like protein | NaN | 0 | 0 | 0 | 0 | 0 | 0 |
| A0A494BBD8;P10107 | Anxa1 | Annexin A1 | NaN | 0 | 0 | 0 | 0 | 494670 | 1029700 |
| A0A498WFS2;Q922Y1 | Ubxn1 | UBX domain-containing protein 1 | NaN | 0 | 0 | 1071800 | 975180 | 0 | 0 |
| G3UZM6;G3UZD6;A0A571BI | Ube4b | Ubiquitin conjugation factor E4 B | NaN | 0 | 0 | 0 | 0 | 0 | 0 |
| A0A571BGH5;A2A610;A0A57 | Gnas | s) subunit alpha isoforms short;Guanine nucleotide-binding protein G(s) subunit alpha | NaN | 0 | 0 | 0 | 0 | 0 | 0 |
| A0A571BEI2;Q6PIU9 | | Uncharacterized protein FLJ45252 homolog | NaN | 0 | 0 | 0 | 0 | 0 | 0 |
| D3YU22;A0A571BG24;Q3UH | Limch1 | LIM and calponin homology domains-containing protein 1 | NaN | 0 | 0 | 0 | 0 | 0 | 0 |
| A0A589M675;Q4ACU6-10;Q4 | Shank3 | SH3 and multiple ankyrin repeat domains protein 3 | NaN | 0 | 0 | 0 | 0 | 0 | 0 |
| A0A5F8MPF6;A0A5F8MPH6; | Arhgef18 | Rho guanine nucleotide exchange factor 18 | NaN | 0 | 0 | 0 | 0 | 0 | 0 |
| A0A5F8MPW8;Q3U9D6;Q8R | Exoc6 | Exocyst complex component 6 | NaN | 0 | 0 | 0 | 0 | 0 | 0 |
| A0A5K1VVQ1;E9Q8N1;E9Q8 | Ttn | Titin | NaN | 3759900 | 652160 | 927390 | 0 | 0 | 0 |
| A0A668KLV9;A0A668KLD3;Q | Akap12 | A-kinase anchor protein 12 | NaN | 0 | 0 | 0 | 0 | 0 | 0 |
| A0A6I8MWW6;Q5SS00 | Zdbf2 | DBF4-type zinc finger-containing protein 2 homolog | NaN | 0 | 0 | 0 | 0 | 0 | 0 |
| Q80YW6;Q80YW9;A0A6I8M\ | Fkbp15 | Peptidyl-prolyl cis-trans isomerase;FK506-binding protein 15 | NaN | 0 | 0 | 0 | 0 | 0 | 0 |
| A0A7N4FLU7;P28028;P2802 | Braf | Serine/threonine-protein kinase B-raf | NaN | 0 | 0 | 0 | 0 | 0 | 0 |
| A0A7N9VR94 | | | NaN | 0 | 0 | 0 | 0 | 0 | 0 |
| A0A7N9VRC4;Q8R0W0 | Eppk1 | Epiplakin | NaN | 0 | 0 | 0 | 0 | 0 | 0 |
| A2A4A6;Q62077 | Plcg1 | ospholipase C;1-phosphatidylinositol 4,5-bisphosphate phosphodiesterase gamma-1 | NaN | 0 | 0 | 0 | 0 | 0 | 0 |
| A2A4H9;Q61576;F6W360 | Fkbp10 | -prolyl cis-trans isomerase FKBP10;Peptidyl-prolyl cis-trans isomerase | NaN | 0 | 0 | 0 | 0 | 0 | 0 |
| A6PW84;A2A7Q5;Q3V1T4-2; | P3h1;Lepre1 | Prolyl 3-hydroxylase 1 | NaN | 0 | 0 | 0 | 0 | 0 | 0 |
| A2A7S7;Q91WQ3 | Yars | ine--tRNA ligase, cytoplasmic;Tyrosine--tRNA ligase, cytoplasmic, N-terminally proce | NaN | 0 | 0 | 0 | 0 | 0 | 0 |
| A2A863-3;A2A864;A2A863-2 | Itgb4 | Integrin beta-4;Integrin beta | NaN | 0 | 0 | 0 | 0 | 0 | 0 |
| A2A9K7 | Cnksr1 | | NaN | 0 | 0 | 0 | 0 | 0 | 0 |
| A6PWC3;A2A9Q2;Q8BHG1;C | Nrd1 | Nardilysin | NaN | 0 | 0 | 0 | 0 | 0 | 0 |
| A2A9X5;Q9JM14 | Nt5c | 5(3)-deoxyribonucleotidase, cytosolic type | NaN | 0 | 0 | 0 | 0 | 0 | 0 |
| A2AAW9;Q9Z0N1 | Eif2s3x | Eukaryotic translation initiation factor 2 subunit 3, X-linked | NaN | 630070 | 3825800 | 0 | 0 | 0 | 0 |
| A2AC13;Q3TLP8;Q05144;P63 | Rac3;Rac1;Rac2 | bstrate 2;Ras-related C3 botulinum toxin substrate 1;Ras-related C3 botulinum toxin s | NaN | 0 | 0 | 0 | 0 | 0 | 0 |
| A2AC29;P70245 | Ebp | 3-beta-hydroxysteroid-Delta(8),Delta(7)-isomerase | NaN | 0 | 0 | 0 | 0 | 3959300 | 2115000 |
| A2ACG7;Q9DBG6 | Rpn2 | nyl-diphosphooligosaccharide--protein glycosyltransferase subunit 2 | NaN | 0 | 0 | 0 | 0 | 0 | 0 |
| A2ADF3;A2ADF2;A2ADF1;A2 | Fblim1 | Filamin-binding LIM protein 1 | NaN | 0 | 0 | 0 | 0 | 0 | 0 |
| A2ADR8;Q8R3G1 | Ppp1r8 | Nuclear inhibitor of protein phosphatase 1 | NaN | 0 | 0 | 0 | 0 | 0 | 0 |
| A2ADY9 | Ddi2 | Protein DDI1 homolog 2 | NaN | 0 | 0 | 0 | 0 | 0 | 0 |
| A2AFQ0;Q7TMY8-4;Q7TMY8 | Huwe1 | E3 ubiquitin-protein ligase HUWE1 | NaN | 0 | 0 | 0 | 0 | 0 | 0 |
| A2AGN7;B7ZCF1;O88685;F6 | Psmc3 | 26S protease regulatory subunit 6A | NaN | 0 | 0 | 2427500 | 1306800 | 0 | 0 |
| A2AHZ5;Q9ESG4 | Tmem27 | Collectrin | NaN | 0 | 0 | 0 | 0 | 0 | 0 |
| A2AJ26;P41234 | Abca2 | ATP-binding cassette sub-family A member 2 | NaN | 0 | 0 | 0 | 0 | 405110 | 2811500 |
| A2BIN0;B8JI96;L7MUC7;Q58 | Mup15;Mup13;Mup10;Mup4;Mup8;Mup9 | nary protein 1;Major urinary protein 6;Major urinary protein 17;Major urinary proteins | NaN | 0 | 0 | 0 | 702530 | 0 | 0 |
| A7TU71;A2ALU4;A2ALU4-2 | Shroom2 | Protein Shroom2 | NaN | 0 | 0 | 0 | 0 | 0 | 0 |
| A2AM80 | Fam43b | | NaN | 0 | 0 | 874320 | 29571000 | 0 | 0 |
| Q6PBC0;A2AMQ5;Q99L43 | Cds2 | phatidate cytidylyltransferase;Phosphatidate cytidylyltransferase 2 | NaN | 0 | 0 | 0 | 0 | 0 | 0 |
| A2AMY5;Q91VX2;A2AMY7 | Ubap2 | Ubiquitin-associated protein 2 | NaN | 0 | 0 | 0 | 0 | 0 | 0 |

| | | | | | | | | | | |
|---|---|---|---|---|---|---|---|---|---|---|
| Z4YMA7;A2AN08-2;A2AN08- | Ubr4 | E3 ubiquitin-protein ligase UBR4 | | NaN | 0 | 0 | 0 | 0 | 0 | 0 |
| A2API8;F7AA26;O54931-5;O! | Akap2;Pakap | A-kinase anchor protein 2 | | NaN | 0 | 0 | 0 | 0 | 0 | 0 |
| A2AQR0;Q64521 | Gpd2 | osphate dehydrogenase;Glycerol-3-phosphate dehydrogenase, mitochondrial | | NaN | 0 | 0 | 2709700 | 2172900 | 0 | 0 |
| A2ATI8;A2ATI6;A2ATI9;Q99J. | Gorasp2 | Golgi reassembly-stacking protein 2 | | NaN | 0 | 0 | 0 | 0 | 0 | 0 |
| Q6PG65;A2ATQ5;P53995 | Anapc1 | Anaphase-promoting complex subunit 1 | | NaN | 0 | 0 | 0 | 0 | 0 | 0 |
| A2AU62;Q64012-2;Q64012 | Raly | RNA-binding protein Raly | | NaN | 0 | 0 | 0 | 0 | 0 | 0 |
| A2AWA9 | Rabgap1 | Rab GTPase-activating protein 1 | | NaN | 0 | 0 | 0 | 0 | 0 | 0 |
| A2AWI9;A2AWI7;Q8R3V5-3; | Sh3glb2 | Endophilin-B2 | | NaN | 0 | 0 | 0 | 0 | 0 | 0 |
| Q3TPJ8;A2BFF9;O88487;A2B | Dync1i2 | Cytoplasmic dynein 1 intermediate chain 2 | | NaN | 0 | 0 | 0 | 0 | 0 | 0 |
| A2CG44;E9Q3M0;Q8BUN5;C | Smad3;Smad2;Smad9 | st decapentaplegic homolog 3;Mothers against decapentaplegic homolog 9;Mothers a | | NaN | 0 | 0 | 0 | 0 | 0 | 0 |
| A3KG36;Q00612 | G6pdx | 5-phosphate 1-dehydrogenase;Glucose-6-phosphate 1-dehydrogenase X | | NaN | 0 | 0 | 0 | 0 | 0 | 0 |
| E9Q0P6;A3KGA8;P46737-2;P | Brcc3 | Lys-63-specific deubiquitinase BRCC36 | | NaN | 0 | 0 | 0 | 0 | 0 | 0 |
| A6BLY7 | Krt28 | Keratin, type I cytoskeletal 28 | | NaN | 0 | 0 | 0 | 0 | 9968600 | 2389400 |
| A6H634 | Gm266 | | | NaN | 0 | 0 | 0 | 0 | 1144600 | 0 |
| A6H6E2 | Mmrn2 | Multimerin-2 | | NaN | 0 | 0 | 0 | 0 | 0 | 0 |
| A6X8Z5 | Arhgap31 | Rho GTPase-activating protein 31 | | NaN | 0 | 0 | 0 | 0 | 0 | 0 |
| B0QZL3;Q8BIV7 | Slc45a1 | Proton-associated sugar transporter A | | NaN | 0 | 0 | 11569000 | 419610 | 0 | 0 |
| B0QZN5;P63044 | Vamp2 | Vesicle-associated membrane protein 2 | | NaN | 0 | 0 | 0 | 0 | 0 | 0 |
| Q3V2Y9;H3BKD1;B1AQF4;Q9 | Dusp3 | Dual specificity protein phosphatase 3 | | NaN | 0 | 0 | 0 | 0 | 0 | 0 |
| B1AQR8;G3X9T7;O08573-3;( | Lgals9 | Galectin;Galectin-9 | | NaN | 0 | 0 | 0 | 0 | 0 | 0 |
| B7ZC46;B1AQY9;B1AQZ0;Q8 | 08-Sep | Septin-8 | | NaN | 0 | 0 | 0 | 0 | 0 | 0 |
| E9QA63;B1ARU4;E9PVY8;Q9 | Macf1 | Microtubule-actin cross-linking factor 1 | | NaN | 0 | 0 | 0 | 0 | 0 | 0 |
| Q3TRH2;B1AT36;Q9D8W5 | Psmd12 | 26S proteasome non-ATPase regulatory subunit 12 | | NaN | 0 | 0 | 0 | 0 | 0 | 0 |
| B1ATL6;P47809 | Map2k4 | Dual specificity mitogen-activated protein kinase kinase 4 | | NaN | 0 | 0 | 0 | 0 | 0 | 0 |
| B1AUD9;Q9DBG9 | Tax1bp3 | Tax1-binding protein 3 | | NaN | 0 | 0 | 0 | 0 | 0 | 0 |
| B1AUX2;Q61191 | Hcfc1 | -terminal chain 5;HCF N-terminal chain 6;HCF C-terminal chain 1;HCF C-terminal chair | | NaN | 0 | 0 | 0 | 0 | 0 | 0 |
| B1AZ42;P0C0A3 | Chmp6 | Charged multivesicular body protein 6 | | NaN | 0 | 0 | 0 | 0 | 0 | 0 |
| Q8BLJ6;B1AZQ9;B1AZR0 | Klhl4 | | | NaN | 0 | 0 | 0 | 0 | 0 | 0 |
| D3Z5B1;B2RPU8;Q9D1L0 | Zbed5;Chchd2 | Coiled-coil-helix-coiled-coil-helix domain-containing protein 2 | | NaN | 0 | 0 | 0 | 0 | 0 | 0 |
| E9QAI5;G3UWN2;B2RQC6-2 | Cad | pendent carbamoyl-phosphate synthase;Aspartate carbamoyltransferase;Dihydroorota | | NaN | 0 | 0 | 0 | 0 | 0 | 0 |
| B2RQS1;Q9ERG2 | Strn3 | Striatin-3 | | NaN | 0 | 0 | 0 | 0 | 0 | 0 |
| S4R2B0;B2RY56 | Rbm25 | RNA-binding protein 25 | | NaN | 0 | 0 | 0 | 0 | 0 | 0 |
| Q9DBQ6;B5B2N5;B5B2N4;Q | Nfatc1 | Nuclear factor of activated T-cells, cytoplasmic 1 | | NaN | 0 | 0 | 0 | 0 | 8774700 | 995480 |
| D6RFS0;Q3THM8;B7FAU5;I7 | Emd | Emerin | | NaN | 0 | 0 | 0 | 0 | 0 | 0 |
| B7ZC18;P42227-2;P42227-3; | Stat3 | r and activator of transcription;Signal transducer and activator of transcription 3 | | NaN | 0 | 0 | 0 | 0 | 0 | 0 |
| B7ZC21;B7ZC22;O70139 | Pkig | cAMP-dependent protein kinase inhibitor gamma | | NaN | 0 | 0 | 0 | 0 | 0 | 0 |
| B7ZCM8;B7ZCN0;B7ZCM9;P( | Pla2g4b;Gm28042 | Cytosolic phospholipase A2 beta | | NaN | 0 | 0 | 0 | 0 | 0 | 0 |
| V9GXM6;F6R587;B7ZCP4;Q8 | Cpne1 | Copine-1 | | NaN | 0 | 0 | 0 | 0 | 0 | 0 |
| B7ZNL2;Q78ZA7;A0A140LJ37 | Nap1l4 | Nucleosome assembly protein 1-like 4 | | NaN | 0 | 0 | 0 | 0 | 0 | 0 |
| B8JJ90;Q52KR6;B8JJ92;B8JJ9 | Acin1 | Apoptotic chromatin condensation inducer in the nucleus | | NaN | 0 | 0 | 0 | 0 | 0 | 0 |
| B9EHJ3;P39447;A0A0U1RPW | Tjp1 | Tight junction protein ZO-1 | | NaN | 0 | 0 | 1690400 | 2310400 | 0 | 0 |
| B9EJR8 | Dnaaf5 | Dynein assembly factor 5, axonemal | | NaN | 0 | 0 | 0 | 0 | 0 | 0 |
| Q6NZJ5;E9Q3I8;E9Q3I9;E9Q3 | Itsn1 | Intersectin-1 | + | NaN | 0 | 0 | 19267000 | 0 | 0 | 0 |
| CON__ENSEMBL:ENSBTAP00000007350 | | | + | NaN | 0 | 0 | 0 | 0 | 0 | 0 |
| CON__ENSEMBL:ENSBTAP00000016046;Q08879;Q08879-2 | | | + | NaN | 0 | 0 | 0 | 0 | 0 | 0 |

| | | | | | | | | | | |
|---|---|---|---|---|---|---|---|---|---|---|
| CON__Q3MHN5;CON__ENSE | Gc | Vitamin D-binding protein | + | NaN | 0 | 0 | 0 | 0 | 0 | 0 |
| CON__ENSEMBL:ENSBTAP00000024146 | | | + | NaN | 0 | 0 | 0 | 0 | 0 | 0 |
| CON__P00978 | | | + | NaN | 0 | 0 | 0 | 0 | 0 | 0 |
| CON__P01966 | | | + | NaN | 0 | 0 | 0 | 0 | 0 | 0 |
| CON__P02070;CON__Q3SX09 | | | + | NaN | 0 | 0 | 0 | 0 | 0 | 0 |
| CON__P02533;CON__Q61782 | | | + | NaN | 0 | 0 | 0 | 0 | 0 | 0 |
| CON__P02666 | | | + | NaN | 0 | 0 | 0 | 0 | 0 | 0 |
| CON__P35527 | | | + | NaN | 0 | 0 | 0 | 0 | 0 | 0 |
| CON__P35908;CON__P48668;CON__P04259;CON__P02538;CON__P19013;CON__Q3TTY5;Q3TTY5 | | | + | NaN | 0 | 0 | 0 | 0 | 0 | 0 |
| CON__Q03247 | | | + | NaN | 0 | 0 | 0 | 0 | 0 | 0 |
| CON__Q2KJF1 | | | + | NaN | 0 | 0 | 0 | 0 | 0 | 0 |
| CON__Q2UVX4 | | | + | NaN | 0 | 0 | 0 | 0 | 0 | 0 |
| CON__Q3T052 | | | + | NaN | 0 | 0 | 0 | 0 | 0 | 0 |
| CON__Q3ZBS7;P29788 | | | + | NaN | 0 | 0 | 0 | 0 | 0 | 0 |
| CON__Q6IFZ6;Q6IFZ6 | Krt77 | Keratin, type II cytoskeletal 1b | + | NaN | 0 | 0 | 7559700 | 6885600 | 0 | 0 |
| CON__Q7Z794 | | | + | NaN | 0 | 0 | 0 | 0 | 0 | 0 |
| D3YU17;Q8VCM8 | Ncln | Nicalin | | NaN | 0 | 0 | 0 | 0 | 0 | 0 |
| D6RET7;D3YUE7;Q8CB44 | Gramd4 | GRAM domain-containing protein 4 | | NaN | 0 | 0 | 0 | 0 | 0 | 0 |
| D6RCZ7;D6RG51;D3YUH9;Q6 | Map4k2 | Mitogen-activated protein kinase kinase kinase kinase 2 | | NaN | 0 | 0 | 0 | 0 | 0 | 0 |
| D3YV10 | Ccdc13 | Coiled-coil domain-containing protein 13 | | NaN | 0 | 0 | 72261 | 9620400 | 0 | 0 |
| D6RH38;D3Z412;D3YVK9;F7E | Syne2 | Nesprin-2 | | NaN | 0 | 0 | 815340 | 997480 | 0 | 0 |
| F6RUD1;D3YVW2;Q8BXA1 | Golim4 | Golgi integral membrane protein 4 | | NaN | 0 | 0 | 0 | 0 | 0 | 0 |
| D3YW40;O08915 | Aip | AH receptor-interacting protein | | NaN | 0 | 0 | 0 | 0 | 0 | 0 |
| D3Z7K0;D3YWF6;Q7TQI3 | Otub1 | Ubiquitin thioesterase OTUB1 | | NaN | 0 | 0 | 0 | 0 | 0 | 0 |
| F7BX63;F7CV24;D6RHL8;D3Y | Donson | Protein downstream neighbor of Son | | NaN | 0 | 0 | 0 | 0 | 0 | 0 |
| D3YWT1;D3Z3N4 | Hnrnph3 | | | NaN | 0 | 0 | 0 | 0 | 0 | 0 |
| F6YTL8;D3YWX2 | Ylpm1 | | | NaN | 0 | 0 | 0 | 0 | 0 | 0 |
| D3Z4R0;D3YWZ1;F6R8S6;E9 | Akt1s1 | Proline-rich AKT1 substrate 1 | | NaN | 0 | 0 | 0 | 0 | 0 | 0 |
| D3YX34;E9Q586;E9Q3M3;O0 | Dctn1 | Dynactin subunit 1 | | NaN | 0 | 0 | 0 | 0 | 0 | 0 |
| D3YX62;O70252 | Hmox2 | Heme oxygenase 2 | | NaN | 0 | 0 | 292380 | 6036300 | 0 | 0 |
| D3YXF8;J3QQ40;Q3UE61;Q9 | Tor1aip1 | Torsin-1A-interacting protein 1 | | NaN | 0 | 0 | 0 | 0 | 0 | 0 |
| D3YXG2;Q9D997;Q9QZ08 | Nagk | N-acetyl-D-glucosamine kinase | | NaN | 0 | 0 | 0 | 0 | 0 | 0 |
| D3YXG6;Q9CVB6 | Arpc2 | Actin-related protein 2/3 complex subunit 2 | | NaN | 0 | 0 | 0 | 0 | 0 | 0 |
| D3YXU1;Q99JY0 | Hadhb | ctional enzyme subunit beta, mitochondrial;3-ketoacyl-CoA thiolase | | NaN | 0 | 0 | 0 | 0 | 0 | 0 |
| D3YY48;Q99KJ6;P12265 | Gusb | Beta-glucuronidase | | NaN | 0 | 0 | 0 | 0 | 0 | 0 |
| F6UFG6;D3YYE1;D3Z7M9;O3 | Anp32a | Acidic leucine-rich nuclear phosphoprotein 32 family member A | | NaN | 0 | 0 | 0 | 0 | 0 | 0 |
| D3YYK8;E9Q6X0;Q8R001-2;C | Mapre2 | Microtubule-associated protein RP/EB family member 2 | | NaN | 0 | 0 | 0 | 0 | 0 | 0 |
| D3YYM6;D3Z1S8;Q91V55;P9 | Rps5 | bosomal protein S5;40S ribosomal protein S5, N-terminally processed | | NaN | 0 | 0 | 0 | 0 | 5015800 | 5107200 |
| D3YZ06;P14602-2;P14602 | Hspb1 | Heat shock protein beta-1 | | NaN | 0 | 0 | 0 | 0 | 0 | 0 |
| D3YZA1;D3Z7T7;E9PW20;Q9 | Chtop | Chromatin target of PRMT1 protein | | NaN | 0 | 0 | 0 | 0 | 0 | 0 |
| D3Z0M2;D3Z191;Q80X71 | Tmem106b | Transmembrane protein 106B | | NaN | 0 | 0 | 0 | 0 | 0 | 0 |
| D6RHS6;D3Z1V4;P70296 | Pebp1 | anolamine-binding protein 1;Hippocampal cholinergic neurostimulating peptide | | NaN | 0 | 0 | 0 | 0 | 0 | 0 |
| E9Q986;D3Z7H6;E9Q907;E9C | Ctnnd1 | Catenin delta-1 | | NaN | 0 | 0 | 422020 | 302420 | 0 | 0 |
| D3Z5X8;D3Z7N3;D3Z345;Q0 | Lyar | Cell growth-regulating nucleolar protein | | NaN | 0 | 0 | 0 | 0 | 0 | 0 |
| D3Z3F1;Q9D824-4;Q9D824-3 | Fip1l1 | Pre-mRNA 3-end-processing factor FIP1 | | NaN | 0 | 0 | 0 | 0 | 0 | 0 |

| | | | | | | | | | |
|---|---|---|---|---|---|---|---|---|---|
| D3Z3Q3;Q921U8-2;Q921U8; | Smtn | Smoothelin | NaN | 0 | 0 | 0 | 0 | 0 | 0 |
| D3Z5B2;Q3UFK8 | Frmd8 | FERM domain-containing protein 8 | NaN | 0 | 0 | 0 | 0 | 0 | 0 |
| D3Z5G4;O88532 | Zfr | Zinc finger RNA-binding protein | NaN | 0 | 0 | 0 | 0 | 0 | 0 |
| D3Z6C9;Q8BJ48 | Nagpa | etylglucosamine-1-phosphodiester alpha-N-acetylglucosaminidase | NaN | 0 | 0 | 0 | 0 | 0 | 0 |
| D3Z6G3 | Mapre3 | | NaN | 0 | 0 | 0 | 0 | 0 | 0 |
| D3Z6K5 | Mrps10 | | NaN | 0 | 0 | 0 | 0 | 12015000 | 0 |
| E9QAT0;D3Z6U8;E9QNF5;Q6 | Fmr1 | Fragile X mental retardation protein 1 homolog | NaN | 0 | 0 | 0 | 0 | 0 | 0 |
| D3Z6Z0;Q78ZM0;D3Z789;O7 | Snx3 | Sorting nexin-3 | NaN | 0 | 0 | 0 | 0 | 0 | 0 |
| D3Z7C0;Q9Z2Q5 | Mrpl40 | 39S ribosomal protein L40, mitochondrial | NaN | 0 | 0 | 0 | 0 | 0 | 0 |
| G3X922;D4AFX7;A0A1L1STR | Dnajc13 | | NaN | 0 | 0 | 0 | 0 | 0 | 0 |
| D6REI7;D6RD00;Q9D7H3 | Rtca;RtcA | RNA 3-terminal phosphate cyclase | NaN | 0 | 0 | 0 | 0 | 0 | 0 |
| D6RFB1;Q9D8C4 | Ifi35 | Interferon-induced 35 kDa protein homolog | NaN | 0 | 0 | 0 | 0 | 0 | 0 |
| F7BHM8;D6RFU2;Q99LR1-2; | Abhd12 | Monoacylglycerol lipase ABHD12 | NaN | 0 | 0 | 0 | 0 | 0 | 0 |
| D6RH37;P70268;P70268-2 | Pkn1 | Serine/threonine-protein kinase N1 | NaN | 0 | 0 | 0 | 0 | 0 | 0 |
| D6RI64;Q9R0P4 | 1110004F10Rik;Smap | Small acidic protein | NaN | 0 | 0 | 0 | 0 | 0 | 0 |
| D9J2Z9;D9J300;D9J301;D9J3 | Pdlim5 | PDZ and LIM domain protein 5 | NaN | 527080 | 1098400 | 0 | 0 | 0 | 0 |
| E0CYQ2;Q9CQ48 | Nudcd2 | NudC domain-containing protein 2 | NaN | 0 | 0 | 0 | 0 | 0 | 0 |
| F6TXE3;F7D432;E9PWE0;E0 | Pcmt1 | ate O-methyltransferase;Protein-L-isoaspartate(D-aspartate) O-methyltransferase | NaN | 0 | 0 | 0 | 0 | 0 | 0 |
| E0CZ22 | Mroh1 | | NaN | 0 | 0 | 0 | 0 | 0 | 0 |
| E9PU87 | Sik3 | | NaN | 0 | 0 | 0 | 0 | 0 | 0 |
| E9PUB0;Q4LDD4-3;Q4LDD4- | Arap1 | with Rho-GAP domain, ANK repeat and PH domain-containing protein 1 | NaN | 0 | 0 | 0 | 0 | 0 | 0 |
| E9PUD2;Q8K1M6-4;Q8K1M6 | Dnm1l | Dynamin-1-like protein | NaN | 0 | 0 | 0 | 0 | 0 | 0 |
| E9PUF7;Q61210-2;Q61210;C | Arhgef1 | Rho guanine nucleotide exchange factor 1 | NaN | 0 | 0 | 0 | 0 | 0 | 0 |
| E9PUX0;Q80WJ7 | Mtdh | Protein LYRIC | NaN | 0 | 0 | 0 | 0 | 0 | 0 |
| E9PV22;Q505F5 | Lrrc47 | Leucine-rich repeat-containing protein 47 | NaN | 0 | 0 | 0 | 0 | 0 | 0 |
| E9PV48;Q64345 | I830012O16Rik;Ifit3 | Interferon-induced protein with tetratricopeptide repeats 3 | NaN | 0 | 0 | 0 | 0 | 0 | 0 |
| E9PV80;Q61026 | Ncoa2 | Nuclear receptor coactivator 2 | NaN | 0 | 0 | 0 | 0 | 0 | 0 |
| E9PVA6;Q9JLQ2;F6SLJ2;F6U | Git2 | ARF GTPase-activating protein GIT2 | NaN | 0 | 0 | 0 | 0 | 0 | 0 |
| E9PVA8 | Gcn1l1 | | NaN | 0 | 0 | 0 | 0 | 0 | 0 |
| E9PVM9;E9Q0T0;G5E8W7;F | Ppt2 | Lysosomal thioesterase PPT2 | NaN | 0 | 0 | 0 | 0 | 0 | 0 |
| E9PVQ9;E9PVN6;Q9D6K5-2; | Gm20498;Synj2bp | Synaptojanin-2-binding protein | NaN | 0 | 0 | 0 | 0 | 0 | 0 |
| E9Q175;E9PVU0;E9Q3L1;E9 | Myo6 | Unconventional myosin-VI | NaN | 0 | 0 | 0 | 0 | 0 | 0 |
| E9QAH1;E9PVZ8 | Golgb1 | | NaN | 0 | 0 | 0 | 0 | 0 | 0 |
| E9PW43;Q9CQS8 | Gm10320;Sec61b | Protein transport protein Sec61 subunit beta | NaN | 0 | 0 | 0 | 0 | 0 | 0 |
| E9PWC5;Q9JLB0-2;Q9JLB0 | Mpp6 | MAGUK p55 subfamily member 6 | NaN | 0 | 0 | 0 | 0 | 0 | 0 |
| F6YTS6;E9PWK1;Q9D379 | Ephx1 | Epoxide hydrolase 1 | NaN | 0 | 0 | 0 | 0 | 0 | 0 |
| E9PWN3;E9PWN2;Q8BIJ6 | Iars2 | Isoleucine--tRNA ligase, mitochondrial | NaN | 0 | 0 | 0 | 0 | 0 | 0 |
| E9PWY9;Q8C0C7;D6RIJ2 | Farsa | Phenylalanine--tRNA ligase alpha subunit | NaN | 0 | 0 | 0 | 0 | 0 | 0 |
| F6V294;E9PX53;E9QPR5;Q8 | Ppp4r1 | Serine/threonine-protein phosphatase 4 regulatory subunit 1 | NaN | 0 | 0 | 0 | 0 | 0 | 0 |
| E9PXW9;Q08369 | Gata4 | Transcription factor GATA-4 | NaN | 0 | 0 | 0 | 0 | 0 | 0 |
| E9PYI8;Q9JMA1 | Usp14 | n carboxyl-terminal hydrolase;Ubiquitin carboxyl-terminal hydrolase 14 | NaN | 0 | 0 | 0 | 0 | 0 | 0 |
| E9PYX7;E9Q852;E9Q9C3;Q9 | Mllt4 | Afadin | NaN | 0 | 0 | 0 | 0 | 0 | 0 |
| E9PZ00;Q8BFQ1;K3W4L3;J3 | Psap | Prosaposin | NaN | 0 | 0 | 0 | 0 | 0 | 0 |
| E9PZ69;P58021 | Tm9sf2 | Transmembrane 9 superfamily member 2 | NaN | 0 | 0 | 0 | 0 | 0 | 0 |
| E9PZ92;Q3TPX4 | Exoc5 | Exocyst complex component 5 | NaN | 0 | 0 | 0 | 0 | 0 | 0 |

| | | | | | | | | | | |
|---|---|---|---|---|---|---|---|---|---|---|
| E9Q9E4;E9PZX7;P30285;D6R | Cdk4 | Cyclin-dependent kinase 4 | | NaN | 0 | 0 | 0 | 0 | 0 | 0 |
| E9Q6F4;E9Q039;Q11011;F6V | Npepps | Puromycin-sensitive aminopeptidase | | NaN | 0 | 0 | 0 | 0 | 0 | 0 |
| E9Q3Y1;E9Q6X2;E9Q108;E9C | Serpinb6a;Serpinb6 | Serpin B6 | | NaN | 0 | 0 | 0 | 0 | 0 | 0 |
| E9Q475;E9Q646;E9Q0Y6;P70 | Ufd1l | Ubiquitin fusion degradation protein 1 homolog | | NaN | 0 | 0 | 0 | 0 | 0 | 0 |
| E9Q1G1;E9Q1H3;G3UY72;G3 | Aldh7a1 | Alpha-aminoadipic semialdehyde dehydrogenase | | NaN | 0 | 0 | 0 | 0 | 0 | 0 |
| E9Q1J7;Q99MN9;A0A087WC | Pccb | Propionyl-CoA carboxylase beta chain, mitochondrial | | NaN | 0 | 0 | 0 | 0 | 0 | 0 |
| J3QMH1;J3QQ55;E9Q1R7;G3 | Tiam1 | T-lymphoma invasion and metastasis-inducing protein 1 | | NaN | 0 | 0 | 0 | 0 | 10233000 | 570410 |
| E9Q1S3;Q01405 | Sec23a | Protein transport protein Sec23A | | NaN | 0 | 0 | 2535400 | 331760 | 0 | 0 |
| F6ZEW4;E9Q1T9;Q9ERK4;E9 | Cse1l | Exportin-2 | | NaN | 0 | 0 | 0 | 0 | 0 | 0 |
| E9Q242;P54822;E9Q3T7;A0A | Adsl | Adenylosuccinate lyase | | NaN | 0 | 0 | 0 | 0 | 0 | 0 |
| G3UY29;E9Q3P9;F8WGS1;G3 | Rab11b;Rab11a | Ras-related protein Rab-11A;Ras-related protein Rab-11B | | NaN | 0 | 0 | 0 | 0 | 0 | 0 |
| E9Q3V6;P42208;F6UKN5;D3Y | 02-Sep | Septin-2 | | NaN | 0 | 0 | 0 | 0 | 0 | 0 |
| Q8BP43;E9Q450;E9Q452;Q8 | Tpm1;Tpm2 | Tropomyosin alpha-1 chain;Tropomyosin beta chain | + | NaN | 0 | 0 | 0 | 0 | 5033700 | 739380 |
| E9Q453;G5E8R2;E9Q456;G5 | Tpm1 | | | NaN | 0 | 0 | 0 | 0 | 6632900 | 2740300 |
| Q6W4W7;E9Q4U7;O70566 | Diap2;Diaph2 | Protein diaphanous homolog 2 | | NaN | 0 | 0 | 0 | 0 | 0 | 0 |
| H3BJG4;E9Q512 | Trip11 | | | NaN | 0 | 0 | 0 | 0 | 0 | 0 |
| E9Q565;Q3UIJ9 | Myzap | Myocardial zonula adherens protein | | NaN | 0 | 0 | 0 | 0 | 0 | 0 |
| E9QNH6;E9Q580;Q7TQD7;P4 | Myo1b | Unconventional myosin-Ib | | NaN | 0 | 0 | 0 | 0 | 0 | 0 |
| E9Q5B2;G5E8X1;Q9DCS2 | 0610011F06Rik | UPF0585 protein C16orf13 homolog | | NaN | 0 | 0 | 0 | 0 | 0 | 0 |
| E9Q5G3 | Kif23 | Kinesin-like protein KIF23 | | NaN | 7182200 | 0 | 0 | 0 | 0 | 0 |
| E9Q5L3;Q9DBL1 | Acadsb | rt/branched chain specific acyl-CoA dehydrogenase, mitochondrial | | NaN | 0 | 0 | 0 | 0 | 0 | 0 |
| E9Q6J8;Q08274 | Dmwd | Dystrophia myotonica WD repeat-containing protein | | NaN | 0 | 0 | 0 | 0 | 0 | 0 |
| E9Q6R3;O08547;A0A0G2JF0 | Sec22b | Vesicle-trafficking protein SEC22b | | NaN | 0 | 0 | 0 | 0 | 0 | 0 |
| E9Q6R7;A0A1W2P7C0;Q616 | Utrn | | | NaN | 0 | 0 | 0 | 0 | 0 | 0 |
| F6T5L3;E9Q717;Q5M8S1;Q9 | Atxn3 | Ataxin-3 | | NaN | 0 | 0 | 0 | 0 | 0 | 0 |
| Q3U3A7;E9Q794;Q3UII8;Q9 | Mta3;Mta1 | tasis-associated protein MTA3;Metastasis-associated protein MTA1 | | NaN | 0 | 0 | 0 | 0 | 168970 | 15344000 |
| E9Q7B0;Q60715-2;Q60715 | P4ha1 | Prolyl 4-hydroxylase subunit alpha-1 | | NaN | 0 | 0 | 0 | 0 | 0 | 0 |
| E9Q7G0;F6ZQA3;A0A1B0GS | Numa1 | | | NaN | 0 | 0 | 0 | 0 | 0 | 0 |
| E9Q800;Q8CAQ8-3;Q8CAQ8- | Immt | MICOS complex subunit Mic60 | | NaN | 0 | 0 | 0 | 0 | 0 | 0 |
| E9Q855;Q3UXS0;O35609 | Scamp3 | Secretory carrier-associated membrane protein 3 | | NaN | 0 | 0 | 0 | 0 | 0 | 0 |
| F7AA45;E9Q8F0;Q8VH51-3;C | Rbm39 | RNA-binding protein 39 | | NaN | 0 | 0 | 0 | 0 | 0 | 0 |
| E9Q9Q7;E9Q9D1;E9Q9D2;E9 | Ablim1 | Actin-binding LIM protein 1 | | NaN | 0 | 0 | 0 | 0 | 0 | 0 |
| H7BWY4;E9Q9H0;Q811D0-2; | Dlg1 | Disks large homolog 1 | | NaN | 0 | 0 | 0 | 0 | 0 | 0 |
| E9Q9X4;P70313 | Nos3 | Nitric oxide synthase;Nitric oxide synthase, endothelial | | NaN | 0 | 0 | 0 | 0 | 0 | 0 |
| E9QAD6;P97450 | Atp5j | ATP synthase-coupling factor 6, mitochondrial | | NaN | 0 | 0 | 0 | 0 | 0 | 0 |
| E9QAF9;Q0VGY8 | Tanc1 | Protein TANC1 | | NaN | 0 | 0 | 0 | 0 | 0 | 0 |
| E9QAS4;E9QAS5;Q6PDQ2;F6 | Chd4;Chd5 | helicase-DNA-binding protein 4;Chromodomain-helicase-DNA-binding protein 5 | | NaN | 0 | 0 | 0 | 0 | 0 | 0 |
| E9QAT4;E9QAT4-2;F7BPW6 | Sec16a | | | NaN | 0 | 0 | 0 | 0 | 0 | 0 |
| E9QMK9;Q8BH86-2;Q8BH86 | 9030617O03Rik | UPF0317 protein C14orf159 homolog, mitochondrial | | NaN | 0 | 0 | 0 | 0 | 0 | 0 |
| E9QN70;P02469 | Lamb1 | Laminin subunit beta-1 | | NaN | 0 | 0 | 0 | 0 | 0 | 0 |
| G5E8R3;E9QPD7;Q05920 | Pcx;Pc | Pyruvate carboxylase;Pyruvate carboxylase, mitochondrial | | NaN | 0 | 0 | 0 | 0 | 0 | 0 |
| E9QPI5;Q6A026;A0A0J9YV33 | Pds5a | Sister chromatid cohesion protein PDS5 homolog A | | NaN | 0 | 0 | 0 | 0 | 0 | 0 |
| E9QPX1;P39061-2;P39061-1; | Col18a1 | Collagen alpha-1(XVIII) chain;Endostatin | | NaN | 0 | 0 | 0 | 0 | 0 | 0 |
| F6Q8V7;O35129;F6QPR1 | Phb2 | Prohibitin-2 | | NaN | 0 | 0 | 0 | 0 | 0 | 0 |
| F6QA74;P28352 | Apex1 | apyrimidinic site) lyase;DNA-(apurinic or apyrimidinic site) lyase, mitochondrial | | NaN | 0 | 0 | 0 | 0 | 0 | 0 |

| | | | | | | | | | |
|---|---|---|---|---|---|---|---|---|---|
| F6ZML1;F6RXI4 | BC067074 | | NaN | 0 | 0 | 0 | 0 | 0 | 0 |
| F6SFF5;Q8K1Z0 | Coq9 | Ubiquinone biosynthesis protein COQ9, mitochondrial | NaN | 0 | 0 | 0 | 0 | 0 | 0 |
| F6SXM5;P32067 | Ssb | Lupus La protein homolog | NaN | 0 | 0 | 0 | 0 | 0 | 0 |
| F7CUP3;F6UP77;Q8JZV7 | Amdhd2 | Putative N-acetylglucosamine-6-phosphate deacetylase | NaN | 0 | 0 | 3081800 | 455320 | 0 | 0 |
| F6XQZ4;Q8BWZ3-2;Q8BWZ3 | Naa25 | N-alpha-acetyltransferase 25, NatB auxiliary subunit | NaN | 0 | 0 | 1848900 | 300060 | 0 | 0 |
| F6XVP7;Q924T7-2;Q924T7 | Rnf31 | E3 ubiquitin-protein ligase RNF31 | NaN | 0 | 0 | 3196100 | 0 | 0 | 0 |
| F6YLI0;Q60865 | Caprin1 | Caprin-1 | NaN | 0 | 0 | 0 | 0 | 0 | 0 |
| Q7M739;F6ZDS4;F6RX08 | Tpr | Nucleoprotein TPR | NaN | 504080 | 813220 | 0 | 0 | 0 | 0 |
| F7ALS6;P05201 | Got1 | Aspartate aminotransferase, cytoplasmic | NaN | 0 | 0 | 0 | 0 | 0 | 0 |
| F7B5B5;Q8VHM5;G3UXU5;S | Hnrnpr;Syncrip | Heterogeneous nuclear ribonucleoprotein Q | NaN | 0 | 0 | 0 | 0 | 0 | 0 |
| F7CBP1;G3XA17;Q62448-2;C | Eif4g2 | Eukaryotic translation initiation factor 4 gamma 2 | NaN | 0 | 0 | 0 | 0 | 0 | 0 |
| F7CUQ1;Q91WX5;O08734 | Bak1 | Bcl-2 homologous antagonist/killer | NaN | 0 | 0 | 0 | 0 | 0 | 0 |
| V9GXH3;V9GXF0;V9GXP8;F8 | Erc1 | ELKS/Rab6-interacting/CAST family member 1 | NaN | 0 | 0 | 0 | 0 | 0 | 0 |
| F8VQJ3;P02468;F6TLW1 | Lamc1 | Laminin subunit gamma-1 | NaN | 0 | 0 | 0 | 0 | 0 | 0 |
| F8VQN6;Q8R4H2 | Arhgef12 | Rho guanine nucleotide exchange factor 12 | NaN | 0 | 0 | 0 | 0 | 0 | 0 |
| F8WGE3;Q8K442 | Abca8a | ATP-binding cassette sub-family A member 8-A | NaN | 0 | 0 | 945270 | 513720 | 0 | 0 |
| F8WGG3;Q9JK81;F7A3N3 | Myg1 | UPF0160 protein MYG1, mitochondrial | NaN | 0 | 0 | 0 | 0 | 0 | 0 |
| F8WHL2;Q8CIE6;F6XJN3 | Copa | Coatomer subunit alpha;Coatomer subunit alpha;Xenin;Proxenin | NaN | 0 | 0 | 0 | 0 | 0 | 0 |
| F8WHM5;Q61543;F6RSH1 | Glg1 | Golgi apparatus protein 1 | NaN | 0 | 0 | 0 | 0 | 0 | 0 |
| F8WHR6;Q569Z5-2;Q569Z5; | Ddx46 | Probable ATP-dependent RNA helicase DDX46 | NaN | 0 | 0 | 0 | 0 | 0 | 0 |
| F8WI30 | Snx7 | | NaN | 0 | 0 | 0 | 0 | 0 | 0 |
| F8WJ30;O70591 | Pfdn2 | Prefoldin subunit 2 | NaN | 0 | 0 | 0 | 0 | 0 | 0 |
| G3UW70;Q9D1L9 | Lamtor5 | Ragulator complex protein LAMTOR5 | NaN | 0 | 0 | 0 | 0 | 0 | 0 |
| G3UXA3;G3UWV3;G3V004;C | Calu | Calumenin | NaN | 0 | 0 | 1874700 | 1975600 | 0 | 0 |
| G3UX48;Q7TSC1 | Prrc2a | Protein PRRC2A | NaN | 0 | 0 | 0 | 0 | 0 | 0 |
| G3UXK7;P53702 | Hccs | Cytochrome c-type heme lyase | NaN | 0 | 0 | 0 | 0 | 0 | 0 |
| G3UXX3;Q91XH5;Q64105;G3 | Spr | Sepiapterin reductase | NaN | 0 | 0 | 1519300 | 709300 | 0 | 0 |
| G3UY42;Q8CCS6-2;Q8CCS6;C | Pabpn1;Gm20521 | Polyadenylate-binding protein 2 | NaN | 0 | 0 | 0 | 0 | 0 | 0 |
| G3UY93;Q9Z1Q9;G3UZ22;G3 | Vars | Valine--tRNA ligase | NaN | 300140 | 288280 | 0 | 0 | 0 | 0 |
| G3UYF9;Q03958 | Pfdn6 | Prefoldin subunit 6 | NaN | 0 | 0 | 4652500 | 1948800 | 0 | 0 |
| G3UYG6;Q6Y7W8;E9Q2M5;C | Gigyf2 | PERQ amino acid-rich with GYF domain-containing protein 2 | NaN | 0 | 0 | 0 | 0 | 0 | 0 |
| G3UYG7;Q6P5E4 | Uggt1 | UDP-glucose:glycoprotein glucosyltransferase 1 | NaN | 0 | 0 | 0 | 0 | 0 | 0 |
| G3UYQ2;G3UZT6;Q3UF95;Q | Bag6 | Large proline-rich protein BAG6 | NaN | 0 | 0 | 0 | 0 | 0 | 0 |
| G3UYU4;O08917 | Flot1 | Flotillin-1 | NaN | 356460 | 228680 | 0 | 0 | 0 | 0 |
| G3UYV7;P62858 | Rps28 | 40S ribosomal protein S28 | NaN | 0 | 0 | 0 | 0 | 0 | 0 |
| G3UYW7;P46938-2;P46938;C | Yap1 | Transcriptional coactivator YAP1 | NaN | 0 | 0 | 0 | 0 | 0 | 0 |
| G3UZ21;Q61193 | Rgl2 | Ral guanine nucleotide dissociation stimulator-like 2 | NaN | 0 | 0 | 0 | 0 | 0 | 0 |
| G3UZJ4;H3BJQ7;P99029-2;P9 | Prdx5 | Peroxiredoxin-5, mitochondrial | NaN | 0 | 0 | 0 | 0 | 0 | 0 |
| G3UZP7;P01899;P01897;V9G | 2-L;H2-Bl;H2-Q7;H2-Q6;H2-Q4;H2-Q2;H2-C | hain;H-2 class I histocompatibility antigen, Q9 alpha chain;H-2 class I histocompatibil | NaN | 0 | 0 | 1094900 | 3777100 | 0 | 0 |
| G3X928;Q6NZC7 | Sec23ip | SEC23-interacting protein | NaN | 0 | 0 | 0 | 0 | 0 | 0 |
| G5DDB7;Q8R3B1;Q05DG3 | Plcd1 | hospholipase C;1-phosphatidylinositol 4,5-bisphosphate phosphodiesterase delta-1 | NaN | 0 | 0 | 0 | 0 | 0 | 0 |
| G5E839;P80315 | Cct4 | T-complex protein 1 subunit delta | NaN | 0 | 0 | 0 | 0 | 0 | 0 |
| G5E884;Q8CIN4;Q61036-2;O | Pak1;Pak2;Pak3 | ne-protein kinase PAK 2;PAK-2p27;PAK-2p34;Serine/threonine-protein kinase PAK 3;Se | NaN | 0 | 0 | 0 | 0 | 0 | 0 |
| G5E897;A0A1L1SRB2 | Kdelc2 | | NaN | 0 | 0 | 436650 | 1175200 | 0 | 0 |
| G5E8E1;A0A087WSF5;A0A0 | Lrrfip1 | Leucine-rich repeat flightless-interacting protein 1 | NaN | 0 | 0 | 0 | 0 | 0 | 0 |

| | | | | | | | | | |
|---|---|---|---|---|---|---|---|---|---|
| G5E8V9;E9QAY5;G5E8V9-2;( | Arfip1 | | NaN | 0 | 0 | 0 | 0 | 0 | 0 |
| G5E902;Q8VEM8 | Slc25a3 | Phosphate carrier protein, mitochondrial | NaN | 0 | 0 | 0 | 0 | 0 | 0 |
| H3BJ02;I1E4X1;Q8K1E0-2;Q8 | Stx5a;Stx5 | Syntaxin-5 | NaN | 0 | 0 | 0 | 0 | 0 | 0 |
| H3BKW0;H3BJ30;H3BJW3;Q( | Cpsf6 | Cleavage and polyadenylation specificity factor subunit 6 | NaN | 0 | 0 | 0 | 0 | 0 | 0 |
| H3BJU7;H3BJ45;H3BJ40;H3B | Arhgef2 | Rho guanine nucleotide exchange factor 2 | NaN | 0 | 0 | 0 | 0 | 0 | 2965000 |
| H3BJ97;Q99JR5 | Tinagl1 | Tubulointerstitial nephritis antigen-like | NaN | 0 | 0 | 243600 | 202360 | 0 | 0 |
| H3BJB6;H3BL49;P42932 | Cct8 | T-complex protein 1 subunit theta | NaN | 0 | 0 | 0 | 0 | 0 | 0 |
| H3BJZ7;Q4KUS2 | Unc13a | Protein unc-13 homolog A | NaN | 97460 | 12678000 | 0 | 0 | 0 | 0 |
| H3BK43;H3BKH6;Q9R0P3;H3I | Esd | S-formylglutathione hydrolase | NaN | 0 | 0 | 1650600 | 1925300 | 0 | 0 |
| H3BLH2;Q9R112 | Sqrdl | Sulfide:quinone oxidoreductase, mitochondrial | NaN | 0 | 0 | 0 | 0 | 0 | 0 |
| H7BWY7;Q9Z1K5 | Arih1 | E3 ubiquitin-protein ligase ARIH1 | NaN | 0 | 0 | 0 | 0 | 0 | 0 |
| H9H9R4;Q9JHJ3;H3BK59;H3B | Glmp | Glycosylated lysosomal membrane protein | NaN | 0 | 0 | 0 | 0 | 0 | 0 |
| H9KV02;H9KV01;H9KV15;H9I | Son | Protein SON | NaN | 0 | 0 | 0 | 0 | 0 | 0 |
| J3QMM7;K3W4M4;Q9CZ42-: | Carkd | ATP-dependent (S)-NAD(P)H-hydrate dehydratase | NaN | 0 | 0 | 0 | 0 | 0 | 0 |
| J3QP56;P97823-2;P97823;J3( | Lypla1;Gm37988 | Acyl-protein thioesterase 1 | NaN | 0 | 0 | 0 | 0 | 0 | 0 |
| J3QP71;K3W4Q8;P18572-2;P | Bsg | Basigin | NaN | 0 | 0 | 0 | 0 | 0 | 0 |
| K3W4T3;Q9Z1G4-3;Q9Z1G4- | Atp6v0a1 | oton ATPase subunit a;V-type proton ATPase 116 kDa subunit a isoform 1 | NaN | 0 | 0 | 0 | 0 | 0 | 0 |
| L7N451;Q80SU7;H3BL64 | Gvin1 | Interferon-induced very large GTPase 1 | NaN | 0 | 0 | 0 | 0 | 0 | 0 |
| M0QWY0;M0QWP2;O70378 | Emc8 | ER membrane protein complex subunit 8 | NaN | 0 | 0 | 0 | 0 | 0 | 0 |
| M0QWP9;Q9D6J6-2;Q9D6J6 | Ndufv2 | IADH dehydrogenase [ubiquinone] flavoprotein 2, mitochondrial | NaN | 0 | 0 | 0 | 0 | 0 | 0 |
| O08529 | Capn2 | Calpain-2 catalytic subunit | NaN | 0 | 0 | 0 | 0 | 0 | 0 |
| O08553 | Dpysl2 | Dihydropyrimidinase-related protein 2 | NaN | 0 | 0 | 0 | 0 | 1456700 | 1685700 |
| O08912 | Galnt1 | ctosaminyltransferase 1;Polypeptide N-acetylgalactosaminyltransferase 1 soluble fori | NaN | 0 | 0 | 0 | 0 | 0 | 0 |
| O08972;Q99LD8;G3UZR0 | Ddah2 | N(G),N(G)-dimethylarginine dimethylaminohydrolase 2 | NaN | 0 | 0 | 0 | 0 | 1122700 | 563370 |
| Q3TMX0;O08992;A2AKJ9;A2 | Sdcbp | Syntenin-1 | NaN | 0 | 0 | 1566400 | 1297500 | 0 | 0 |
| O09005 | Degs1 | Sphingolipid delta(4)-desaturase DES1 | NaN | 0 | 0 | 0 | 0 | 4636800 | 1520700 |
| O09053 | Wrn | Werner syndrome ATP-dependent helicase homolog | NaN | 0 | 0 | 0 | 0 | 0 | 7481100 |
| O09159 | Man2b1 | Lysosomal alpha-mannosidase | NaN | 0 | 0 | 1460900 | 1118000 | 0 | 0 |
| O35127 | Grcc10 | Protein C10 | NaN | 0 | 0 | 0 | 0 | 0 | 0 |
| O35295 | Purb | Transcriptional activator protein Pur-beta | NaN | 1431100 | 3449900 | 0 | 0 | 0 | 0 |
| Q8C391;Q9CXE1;O35382 | Exoc4 | Exocyst complex component 4 | NaN | 0 | 0 | 0 | 0 | 0 | 0 |
| O35465;O35465-2;F6WP10 | Fkbp8 | Peptidyl-prolyl cis-trans isomerase FKBP8 | NaN | 0 | 0 | 0 | 0 | 0 | 0 |
| O35598 | Adam10 | Disintegrin and metalloproteinase domain-containing protein 10 | NaN | 0 | 0 | 0 | 0 | 0 | 0 |
| O35608 | Angpt2 | Angiopoietin-2 | NaN | 0 | 0 | 0 | 0 | 0 | 0 |
| O35704 | Sptlc1 | Serine palmitoyltransferase 1 | NaN | 0 | 0 | 0 | 0 | 0 | 0 |
| Q8C2Q7;O35737 | Hnrnph1 | ibonucleoprotein H;Heterogeneous nuclear ribonucleoprotein H, N-terminally processe | NaN | 0 | 0 | 0 | 0 | 0 | 0 |
| O35857 | Timm44 | Mitochondrial import inner membrane translocase subunit TIM44 | NaN | 0 | 0 | 0 | 0 | 0 | 0 |
| O35901;O35900 | Lsm2 | U6 snRNA-associated Sm-like protein LSm2 | NaN | 0 | 0 | 0 | 0 | 0 | 0 |
| O54724 | Ptrf | Polymerase I and transcript release factor | NaN | 0 | 0 | 0 | 0 | 0 | 0 |
| O54734 | Ddost | -diphosphooligosaccharide--protein glycosyltransferase 48 kDa subunit | NaN | 0 | 0 | 0 | 0 | 0 | 0 |
| O54782;F6TMZ3 | Man2b2 | Epididymis-specific alpha-mannosidase | NaN | 0 | 0 | 1439200 | 697550 | 0 | 0 |
| O54941 | Smarce1 | natrix-associated actin-dependent regulator of chromatin subfamily E member 1 | NaN | 0 | 0 | 0 | 0 | 0 | 0 |
| O54988-2;O54988 | Slk | STE20-like serine/threonine-protein kinase | NaN | 0 | 0 | 0 | 0 | 0 | 0 |
| O55135;A6PWZ2 | Eif6 | Eukaryotic translation initiation factor 6 | NaN | 0 | 0 | 0 | 0 | 0 | 0 |
| O55201-2;O55201 | Supt5h | Transcription elongation factor SPT5 | NaN | 0 | 0 | 0 | 0 | 0 | 0 |

| | | | | | | | | | |
|---|---|---|---|---|---|---|---|---|---|
| Q8BTY5;O55234 | Psmb5 | Proteasome subunit beta type-5 | NaN | 0 | 0 | 0 | 0 | 0 | 0 |
| O70194 | Eif3d | Eukaryotic translation initiation factor 3 subunit D | NaN | 0 | 0 | 675440 | 275220 | 0 | 0 |
| Q8BH40;O70439 | Stx7 | Syntaxin-7 | NaN | 0 | 0 | 0 | 0 | 0 | 0 |
| O70475;D3YXP9 | Ugdh | UDP-glucose 6-dehydrogenase | NaN | 0 | 0 | 0 | 0 | 0 | 0 |
| O88342;A0A0J9YU05 | Wdr1 | WD repeat-containing protein 1 | NaN | 0 | 0 | 0 | 0 | 0 | 0 |
| Q5D0E0;O88351 | Ikbkb | Inhibitor of nuclear factor kappa-B kinase subunit beta | NaN | 0 | 0 | 0 | 0 | 0 | 0 |
| Q91XH6;O88384;F6UHS3;E0( | Vti1b | 'esicle transport through interaction with t-SNAREs homolog 1B | NaN | 0 | 0 | 0 | 0 | 0 | 0 |
| O88544;F6QTS1;D3Z1R9;D3Y | Cops4 | COP9 signalosome complex subunit 4 | NaN | 286850 | 346310 | 0 | 0 | 0 | 0 |
| O88587-2;O88587 | Comt | Catechol O-methyltransferase | NaN | 0 | 0 | 940330 | 512620 | 0 | 0 |
| O88630 | Gosr1 | Golgi SNAP receptor complex member 1 | NaN | 0 | 0 | 0 | 0 | 0 | 0 |
| Q3UDC3;O88746;A0A1D5RM | Tom1 | Target of Myb protein 1 | NaN | 0 | 0 | 0 | 0 | 0 | 0 |
| O88844;A0A087WRM4;D3YV | Idh1 | Isocitrate dehydrogenase [NADP] cytoplasmic | NaN | 0 | 0 | 1844300 | 1721900 | 0 | 0 |
| O89017 | Lgmn | Legumain | NaN | 0 | 0 | 0 | 0 | 0 | 0 |
| O89023 | Tpp1 | Tripeptidyl-peptidase 1 | NaN | 0 | 0 | 0 | 0 | 0 | 0 |
| O89086;Q8BG13;S4R2M6 | Rbm3 | RNA-binding protein 3 | NaN | 0 | 0 | 0 | 0 | 0 | 0 |
| P00520-2;P00520-3;P00520;I | Abl1 | Tyrosine-protein kinase ABL1 | NaN | 0 | 0 | 0 | 0 | 0 | 0 |
| P01101 | Fos | Proto-oncogene c-Fos | NaN | 0 | 0 | 4689200 | 14255000 | 0 | 0 |
| P01790;P01789;P01794;P017 | Ighv7-1 | Ig heavy chain V region HPCG8;Ig heavy chain V region HPCM6;Ig heavy chain V region | NaN | 0 | 0 | 0 | 0 | 0 | 0 |
| P01803 | | Ig heavy chain V region AMPC1 | NaN | 0 | 0 | 0 | 0 | 3336700 | 138130 |
| P01887 | B2m | Beta-2-microglobulin | NaN | 0 | 0 | 0 | 0 | 0 | 0 |
| P03930 | Mtatp8 | ATP synthase protein 8 | NaN | 0 | 0 | 0 | 0 | 0 | 0 |
| P03958 | Ada | Adenosine deaminase | NaN | 4061300 | 1539900 | 0 | 0 | 0 | 0 |
| P05063 | Aldoc | Fructose-bisphosphate aldolase C | NaN | 0 | 0 | 0 | 0 | 0 | 0 |
| P06797 | Ctsl | Cathepsin L1;Cathepsin L1 heavy chain;Cathepsin L1 light chain | NaN | 0 | 0 | 0 | 0 | 0 | 0 |
| Q3TQP6;P06801 | Me1 | Malic enzyme;NADP-dependent malic enzyme | NaN | 0 | 0 | 0 | 0 | 0 | 0 |
| P08030;A0A1D5RLR6 | Aprt | Adenine phosphoribosyltransferase | NaN | 0 | 0 | 0 | 0 | 0 | 0 |
| P08122 | Col4a2 | Collagen alpha-2(IV) chain;Canstatin | NaN | 0 | 0 | 0 | 0 | 0 | 0 |
| P09055;P09055-2 | Itgb1 | Integrin beta-1 | NaN | 0 | 0 | 0 | 0 | 0 | 0 |
| P10493 | Nid1 | Nidogen-1 | NaN | 0 | 0 | 0 | 0 | 0 | 0 |
| P10605 | Ctsb | Cathepsin B;Cathepsin B light chain;Cathepsin B heavy chain | NaN | 0 | 0 | 0 | 0 | 0 | 0 |
| Q8CBB6;Q8CGP2;Q8CGP1;Q( | Hist2h2bb;Hist1h2bh;Hist1h2bb;Hist1h2bm | 2-B;Histone H2B type 1-H;Histone H2B type 1-B;Histone H2B type 1-M;Histone H2B t | NaN | 0 | 0 | 0 | 0 | 0 | 0 |
| P11438 | Lamp1 | Lysosome-associated membrane glycoprotein 1 | NaN | 0 | 0 | 0 | 0 | 0 | 0 |
| P11688 | Itga5 | rin alpha-5;Integrin alpha-5 heavy chain;Integrin alpha-5 light chain | NaN | 0 | 0 | 0 | 0 | 0 | 0 |
| P11881-8;P11881-7;P11881-( | Itpr1 | Inositol 1,4,5-trisphosphate receptor type 1 | NaN | 0 | 0 | 0 | 0 | 0 | 0 |
| P11983;P11983-2;F2Z483 | Tcp1 | T-complex protein 1 subunit alpha | NaN | 0 | 0 | 0 | 0 | 0 | 0 |
| P13597-2;P13597 | Icam1 | Intercellular adhesion molecule 1 | NaN | 0 | 0 | 0 | 0 | 0 | 0 |
| P14211 | Calr | Calreticulin | NaN | 0 | 0 | 0 | 0 | 0 | 0 |
| P14685;F7B7L8 | Psmd3 | 26S proteasome non-ATPase regulatory subunit 3 | NaN | 0 | 0 | 0 | 0 | 0 | 0 |
| P14873 | Map1b | ıbule-associated protein 1B;MAP1B heavy chain;MAP1 light chain LC1 | NaN | 0 | 0 | 0 | 0 | 0 | 0 |
| Q3USD9;P15975-2;P15975 | Usp53 | Inactive ubiquitin carboxyl-terminal hydrolase 53 | NaN | 0 | 0 | 468520 | 541430 | 0 | 0 |
| P17427;A0A140LHG0;P1742( | Ap2a2 | AP-2 complex subunit alpha-2 | NaN | 0 | 0 | 0 | 0 | 0 | 0 |
| Q61696;P17879;P16627 | Hspa1a;Hspa1b;Hspa1l | kDa protein 1A;Heat shock 70 kDa protein 1B;Heat shock 70 kDa protein 1-like | NaN | 0 | 0 | 0 | 0 | 0 | 0 |
| P18406 | Cyr61 | Protein CYR61 | NaN | 0 | 0 | 0 | 0 | 1487900 | 740990 |
| P19324;A0A140LHR4;A0A14( | Serpinh1 | Serpin H1 | NaN | 0 | 0 | 0 | 0 | 0 | 0 |
| P19783;M0QWX7 | Cox4i1 | Cytochrome c oxidase subunit 4 isoform 1, mitochondrial | NaN | 0 | 0 | 0 | 0 | 3835400 | 450630 |

| | | | | | | | | | |
|---|---|---|---|---|---|---|---|---|---|
| P19788 | Mgp | Matrix Gla protein | NaN | 0 | 0 | 0 | 0 | 0 | 0 |
| P21278;A0A494BBL5;P30677 | Gna11;Gna14 | nding protein subunit alpha-11;Guanine nucleotide-binding protein subunit alpha-14 | NaN | 0 | 0 | 0 | 0 | 0 | 0 |
| P21279 | Gnaq | Guanine nucleotide-binding protein G(q) subunit alpha | NaN | 0 | 0 | 0 | 0 | 0 | 0 |
| P21956-2;P21956 | Mfge8 | Lactadherin | NaN | 0 | 0 | 0 | 0 | 0 | 0 |
| P23116 | Eif3a | Eukaryotic translation initiation factor 3 subunit A | NaN | 0 | 0 | 0 | 0 | 0 | 0 |
| P23780 | Glb1 | Beta-galactosidase | NaN | 0 | 0 | 0 | 0 | 0 | 0 |
| P24270;A2AL20 | Cat | Catalase | NaN | 0 | 0 | 0 | 0 | 0 | 0 |
| P24668 | M6pr | Cation-dependent mannose-6-phosphate receptor | NaN | 0 | 0 | 0 | 0 | 0 | 0 |
| P27046 | Man2a1 | Alpha-mannosidase 2 | NaN | 0 | 0 | 0 | 0 | 0 | 0 |
| P28271 | Aco1 | Cytoplasmic aconitate hydratase | NaN | 0 | 0 | 0 | 0 | 0 | 0 |
| P28651 | Ca8 | Carbonic anhydrase-related protein | NaN | 0 | 0 | 0 | 0 | 0 | 0 |
| Q3TF41;Q8BSH9;P28656 | Nap1l1 | Nucleosome assembly protein 1-like 1 | NaN | 0 | 0 | 0 | 0 | 0 | 0 |
| Q3U9N4;P28798;H3BJE0 | Grn | nin;Granulin-1;Granulin-2;Granulin-3;Granulin-4;Granulin-5;Granulin-6;Granulin-7 | NaN | 0 | 0 | 0 | 0 | 0 | 0 |
| Q68FM4;P28828;A0A286YDL | Ptprm | -tyrosine-phosphatase;Receptor-type tyrosine-protein phosphatase mu | NaN | 0 | 0 | 0 | 0 | 4266900 | 218230 |
| Q9CPX4;P29391;A0A1B0GR6 | Ftl1;Ftl2 | Ferritin;Ferritin light chain 1;Ferritin light chain 2 | NaN | 0 | 0 | 0 | 0 | 0 | 0 |
| P29416 | Hexa | Beta-hexosaminidase subunit alpha | NaN | 0 | 0 | 0 | 0 | 0 | 0 |
| P30412 | Ppic | Peptidyl-prolyl cis-trans isomerase C | NaN | 0 | 0 | 0 | 0 | 0 | 0 |
| P30416 | Fkbp4 | s isomerase FKBP4;Peptidyl-prolyl cis-trans isomerase FKBP4, N-terminally processed | NaN | 0 | 0 | 0 | 0 | 0 | 0 |
| P31001 | Des | Desmin | NaN | 0 | 0 | 0 | 0 | 0 | 0 |
| P31324 | Prkar2b | cAMP-dependent protein kinase type II-beta regulatory subunit | NaN | 0 | 0 | 0 | 0 | 0 | 0 |
| P32507-2;P32507 | Pvrl2 | Nectin-2 | NaN | 0 | 0 | 0 | 0 | 0 | 0 |
| P32921-2;P32921 | Wars | Tryptophan--tRNA ligase, cytoplasmic;T1-TrpRS;T2-TrpRS | NaN | 0 | 0 | 0 | 0 | 0 | 0 |
| P35235;P35235-1 | Ptpn11 | Tyrosine-protein phosphatase non-receptor type 11 | NaN | 0 | 0 | 0 | 0 | 0 | 0 |
| Q8C266;P35278 | Rab5c | Ras-related protein Rab-5C | NaN | 0 | 0 | 0 | 0 | 0 | 0 |
| P35282;A0A1W2P6Z4 | Rab21 | Ras-related protein Rab-21 | NaN | 0 | 0 | 0 | 0 | 0 | 0 |
| P35456 | Plaur | Urokinase plasminogen activator surface receptor | NaN | 0 | 0 | 0 | 0 | 0 | 0 |
| P35486 | Pdha1 | ehydrogenase E1 component subunit alpha, somatic form, mitochondrial | NaN | 0 | 0 | 0 | 0 | 0 | 0 |
| Q05DV1;P37040;F6R7H8;E9F | Por | NADPH--cytochrome P450 reductase | NaN | 0 | 0 | 0 | 0 | 0 | 0 |
| P37889-2;P37889 | Fbln2 | Fibulin-2 | NaN | 0 | 0 | 0 | 0 | 0 | 0 |
| P40336;P40336-2 | Vps26a | Vacuolar protein sorting-associated protein 26A | NaN | 0 | 0 | 0 | 0 | 0 | 0 |
| P42669 | Pura | Transcriptional activator protein Pur-alpha | NaN | 0 | 0 | 0 | 0 | 0 | 0 |
| P46061;A0A2R8W753;A0A2F | Rangap1 | Ran GTPase-activating protein 1 | NaN | 0 | 0 | 0 | 0 | 0 | 0 |
| P46414 | Cdkn1b | Cyclin-dependent kinase inhibitor 1B | NaN | 0 | 0 | 0 | 0 | 0 | 0 |
| P46467 | Vps4b | Vacuolar protein sorting-associated protein 4B | NaN | 0 | 0 | 0 | 0 | 0 | 0 |
| P46935;A0A571BEP7;A0A57 | Nedd4 | E3 ubiquitin-protein ligase NEDD4 | NaN | 0 | 0 | 0 | 0 | 0 | 0 |
| P48437 | Prox1 | Prospero homeobox protein 1 | NaN | 0 | 0 | 0 | 0 | 0 | 0 |
| P49070 | Camlg | Calcium signal-modulating cyclophilin ligand | NaN | 0 | 0 | 0 | 0 | 0 | 0 |
| P49717 | Mcm4 | DNA replication licensing factor MCM4 | NaN | 0 | 0 | 0 | 0 | 0 | 0 |
| P50427 | Sts | Steryl-sulfatase | NaN | 0 | 0 | 0 | 0 | 0 | 0 |
| P50580-2;P50580 | Pa2g4 | Proliferation-associated protein 2G4 | NaN | 0 | 0 | 0 | 0 | 0 | 0 |
| P50636 | Rnf19a | E3 ubiquitin-protein ligase RNF19A | NaN | 0 | 0 | 0 | 0 | 0 | 0 |
| P51480;P51480-2 | Cdkn2a | Cyclin-dependent kinase inhibitor 2A | NaN | 0 | 0 | 0 | 0 | 0 | 0 |
| Q8BGZ6;P51569 | Gla | Alpha-galactosidase A | NaN | 0 | 0 | 0 | 0 | 0 | 0 |
| P51660 | Hsd17b4 | nctional enzyme type 2;(3R)-hydroxyacyl-CoA dehydrogenase;Enoyl-CoA hydratase 2 | NaN | 0 | 0 | 0 | 0 | 0 | 0 |
| P52293;A2A600;A2A601;A6P | Kpna2 | Importin subunit alpha-1 | NaN | 0 | 0 | 0 | 0 | 0 | 0 |

| | | | | | | | | | |
|---|---|---|---|---|---|---|---|---|---|
| P52479;P52479-2 | Usp10 | Ubiquitin carboxyl-terminal hydrolase 10 | NaN | 0 | 0 | 0 | 0 | 0 | 0 |
| P52825;A2A8E7;A2A8E8 | Cpt2 | Carnitine O-palmitoyltransferase 2, mitochondrial | NaN | 0 | 0 | 0 | 0 | 0 | 0 |
| P52875 | Tmem165 | Transmembrane protein 165 | NaN | 0 | 0 | 0 | 0 | 0 | 0 |
| P53986 | Slc16a1 | Monocarboxylate transporter 1 | NaN | 0 | 0 | 0 | 0 | 0 | 0 |
| P53994;A0A1D5RMH1;S4R23 | Rab2a;Rab4b;Rab2b | Ras-related protein Rab-2A;Ras-related protein Rab-2B | NaN | 0 | 0 | 0 | 0 | 0 | 0 |
| P54116 | Stom | Erythrocyte band 7 integral membrane protein | NaN | 0 | 0 | 0 | 0 | 0 | 0 |
| P54728 | Rad23b | UV excision repair protein RAD23 homolog B | NaN | 0 | 0 | 0 | 0 | 0 | 0 |
| P54823 | Ddx6 | Probable ATP-dependent RNA helicase DDX6 | NaN | 0 | 0 | 0 | 0 | 0 | 0 |
| P55284 | Cdh5 | Cadherin-5 | NaN | 0 | 0 | 0 | 0 | 0 | 0 |
| P55302 | Lrpap1 | Alpha-2-macroglobulin receptor-associated protein | NaN | 0 | 0 | 0 | 0 | 0 | 0 |
| P56389 | Cda | Cytidine deaminase | NaN | 0 | 0 | 0 | 0 | 0 | 0 |
| Q3U4W8;P56399;D3YYA5;D3 | Usp5 | n carboxyl-terminal hydrolase;Ubiquitin carboxyl-terminal hydrolase 5 | NaN | 0 | 0 | 0 | 0 | 0 | 0 |
| P56812;D3Z7Q5 | Pdcd5 | Programmed cell death protein 5 | NaN | 0 | 0 | 0 | 0 | 0 | 0 |
| P57716 | Ncstn | Nicastrin | NaN | 0 | 0 | 0 | 0 | 0 | 0 |
| V9GXI9;P58404-2;P58404 | Strn4 | Striatin-4 | NaN | 0 | 0 | 0 | 0 | 0 | 0 |
| P58871;Z4YJL4;A0A571BEG9 | Tnks1bp1 | 182 kDa tankyrase-1-binding protein | NaN | 0 | 0 | 0 | 0 | 0 | 0 |
| P58929 | Gmeb2 | Glucocorticoid modulatory element-binding protein 2 | NaN | 1335300 | 22077000 | 0 | 0 | 0 | 0 |
| P59999;Q3TX55;E9PWA7 | Arpc4 | Actin-related protein 2/3 complex subunit 4 | NaN | 0 | 0 | 0 | 0 | 0 | 0 |
| P60122 | Ruvbl1 | RuvB-like 1 | NaN | 0 | 0 | 0 | 0 | 0 | 0 |
| P60670-2;P60670 | Nploc4 | Nuclear protein localization protein 4 homolog | NaN | 0 | 0 | 0 | 0 | 0 | 0 |
| P61089 | Ube2n | Ubiquitin-conjugating enzyme E2 N | NaN | 0 | 0 | 0 | 0 | 0 | 0 |
| P61804 | Dad1 | l-diphosphooligosaccharide--protein glycosyltransferase subunit DAD1 | NaN | 0 | 0 | 0 | 0 | 0 | 0 |
| P61967 | Ap1s1 | AP-1 complex subunit sigma-1A | NaN | 0 | 0 | 0 | 0 | 0 | 0 |
| P62071;A0A1B0GRG1;A0A1E | Rras2 | Ras-related protein R-Ras2 | NaN | 0 | 0 | 0 | 0 | 0 | 0 |
| P62077 | Timm8b | itochondrial import inner membrane translocase subunit Tim8 B | NaN | 467120 | 246010 | 0 | 0 | 0 | 0 |
| P62137 | Ppp1ca | erine/threonine-protein phosphatase PP1-alpha catalytic subunit | NaN | 0 | 0 | 0 | 0 | 0 | 0 |
| Q8K1K2;P62196 | Psmc5 | 26S protease regulatory subunit 8 | NaN | 0 | 0 | 0 | 0 | 0 | 0 |
| P62307 | Snrpf | Small nuclear ribonucleoprotein F | NaN | 0 | 0 | 0 | 0 | 0 | 0 |
| P62311 | Lsm3 | U6 snRNA-associated Sm-like protein LSm3 | NaN | 0 | 0 | 0 | 0 | 0 | 0 |
| P62317 | Snrpd2 | Small nuclear ribonucleoprotein Sm D2 | NaN | 0 | 0 | 0 | 0 | 0 | 0 |
| P62320 | Snrpd3 | Small nuclear ribonucleoprotein Sm D3 | NaN | 0 | 0 | 5681200 | 10919000 | 0 | 0 |
| P62334 | Psmc6 | 26S protease regulatory subunit 10B | NaN | 0 | 0 | 0 | 0 | 0 | 0 |
| P62715;P63330;P97470 | Ppp2cb;Ppp2ca | catalytic subunit beta isoform;Serine/threonine-protein phosphatase 2A catalytic sub | NaN | 0 | 0 | 0 | 0 | 0 | 0 |
| P62869 | Tceb2 | Transcription elongation factor B polypeptide 2 | NaN | 0 | 0 | 0 | 0 | 0 | 0 |
| P63024 | Vamp3 | Vesicle-associated membrane protein 3 | NaN | 0 | 0 | 0 | 0 | 0 | 0 |
| P63037;B1AXY0;B1AXY1 | Dnaja1 | DnaJ homolog subfamily A member 1 | NaN | 0 | 0 | 0 | 0 | 0 | 0 |
| P63213 | Gng2 | anine nucleotide-binding protein G(I)/G(S)/G(O) subunit gamma-2 | NaN | 0 | 0 | 0 | 0 | 0 | 0 |
| P68040 | Gnb2l1 | ubunit beta-2-like 1;Guanine nucleotide-binding protein subunit beta-2-like 1, N-termi | NaN | 0 | 0 | 0 | 0 | 0 | 0 |
| P68369;P05214 | Tuba1a;Tuba3a | Tubulin alpha-1A chain;Tubulin alpha-3 chain | NaN | 0 | 0 | 0 | 0 | 0 | 0 |
| P68373 | Tuba1c | Tubulin alpha-1C chain | NaN | 0 | 0 | 0 | 0 | 0 | 0 |
| P70168 | Kpnb1 | Importin subunit beta-1 | NaN | 0 | 0 | 0 | 0 | 0 | 0 |
| P70195 | Psmb7 | Proteasome subunit beta type-7 | NaN | 0 | 0 | 0 | 0 | 0 | 0 |
| P70227 | Itpr3 | Inositol 1,4,5-trisphosphate receptor type 3 | NaN | 0 | 0 | 0 | 0 | 0 | 0 |
| P70333 | Hnrnph2 | Heterogeneous nuclear ribonucleoprotein H2 | NaN | 0 | 0 | 0 | 0 | 0 | 0 |
| P70445 | Eif4ebp2 | Eukaryotic translation initiation factor 4E-binding protein 2 | NaN | 0 | 0 | 14234000 | 21372000 | 0 | 0 |

| | | | | | | | | | |
|---|---|---|---|---|---|---|---|---|---|
| P70460 | Vasp | Vasodilator-stimulated phosphoprotein | NaN | 0 | 0 | 0 | 0 | 0 | 0 |
| Q60817;P70670 | Naca | mplex subunit alpha;Nascent polypeptide-associated complex subunit alpha, muscle-s | NaN | 0 | 0 | 0 | 0 | 0 | 0 |
| P70697;A0A0A0MQG7 | Urod | Uroporphyrinogen decarboxylase | NaN | 0 | 0 | 0 | 0 | 0 | 0 |
| P70699;F6VEG4;A2AFL5;F6R | Gaa | Lysosomal alpha-glucosidase | NaN | 0 | 0 | 0 | 0 | 0 | 0 |
| Q52KG9;P80317;B1AT05;Q6 | Cct6a;Cct6b | omplex protein 1 subunit zeta;T-complex protein 1 subunit zeta-2 | NaN | 0 | 0 | 0 | 0 | 0 | 0 |
| Q3UKN6;P81117 | Nucb2 | Nucleobindin-2;Nesfatin-1 | NaN | 0 | 0 | 0 | 0 | 0 | 0 |
| P83870 | Phf5a | PHD finger-like domain-containing protein 5A | NaN | 0 | 0 | 0 | 0 | 0 | 0 |
| P84104-2;P84104;A0A3Q4EH | Srsf3 | Serine/arginine-rich splicing factor 3 | NaN | 0 | 0 | 0 | 0 | 0 | 0 |
| P97310 | Mcm2 | DNA replication licensing factor MCM2 | NaN | 0 | 0 | 0 | 0 | 0 | 0 |
| Q3ULG5;P97311 | Mcm6 | DNA helicase;DNA replication licensing factor MCM6 | NaN | 0 | 0 | 0 | 0 | 0 | 0 |
| P97314;A0A1W2P780;A0A1V | Csrp2 | Cysteine and glycine-rich protein 2 | NaN | 0 | 0 | 0 | 0 | 0 | 0 |
| P97346;P97346-2 | Nxn | Nucleoredoxin | NaN | 0 | 0 | 0 | 0 | 0 | 0 |
| P97377-2;P97377 | Cdk2 | Cyclin-dependent kinase 2 | NaN | 0 | 0 | 0 | 0 | 0 | 0 |
| P97384;D3Z7U0 | Anxa11 | Annexin A11;Annexin | NaN | 0 | 0 | 0 | 0 | 0 | 0 |
| Q6P1J1;P97427 | Crmp1 | Dihydropyrimidinase-related protein 1 | NaN | 0 | 0 | 0 | 0 | 0 | 0 |
| P97737 | Gdf10 | Growth/differentiation factor 10 | NaN | 0 | 0 | 1468400 | 966510 | 0 | 0 |
| P97819 | Pla2g6 | 85/88 kDa calcium-independent phospholipase A2 | NaN | 0 | 0 | 0 | 0 | 0 | 0 |
| P97825 | Hn1 | xpressed 1 protein;Hematological and neurological expressed 1 protein, N-terminally | NaN | 0 | 0 | 0 | 0 | 1652600 | 1161500 |
| P97855 | G3bp1 | Ras GTPase-activating protein-binding protein 1 | NaN | 0 | 0 | 0 | 0 | 0 | 0 |
| P97864;A0A5F8MPP6 | Casp7 | Caspase-7;Caspase-7 subunit p20;Caspase-7 subunit p11 | NaN | 0 | 0 | 0 | 0 | 0 | 0 |
| P99028 | Uqcrh | Cytochrome b-c1 complex subunit 6, mitochondrial | NaN | 0 | 0 | 0 | 0 | 0 | 0 |
| V9GX00;Q00560 | Il6st | Interleukin-6 receptor subunit beta | NaN | 0 | 0 | 0 | 0 | 0 | 0 |
| Q00PI9 | Hnrnpul2 | Heterogeneous nuclear ribonucleoprotein U-like protein 2 | NaN | 0 | 0 | 0 | 0 | 0 | 0 |
| Q02013 | Aqp1 | Aquaporin-1 | NaN | 0 | 0 | 0 | 0 | 0 | 0 |
| Q02248;E9Q6A9;F7CRC6;F7E | Ctnnb1 | Catenin beta-1 | NaN | 0 | 0 | 0 | 0 | 0 | 0 |
| Q04447 | Ckb | Creatine kinase B-type | NaN | 0 | 0 | 880470 | 306350 | 0 | 0 |
| Q05186 | Rcn1 | Reticulocalbin-1 | NaN | 0 | 0 | 0 | 0 | 0 | 0 |
| Q05D44 | Eif5b | Eukaryotic translation initiation factor 5B | NaN | 0 | 0 | 0 | 0 | 0 | 0 |
| Q6PAP2;Q06806 | Tie1 | Tyrosine-protein kinase receptor Tie-1 | NaN | 0 | 0 | 0 | 0 | 0 | 0 |
| Q07113 | Igf2r | Cation-independent mannose-6-phosphate receptor | NaN | 0 | 0 | 0 | 0 | 0 | 0 |
| Q07797;E9Q5X5 | Lgals3bp | Galectin-3-binding protein | NaN | 0 | 0 | 0 | 0 | 0 | 0 |
| Q0VBL3-2;Q0VBL3 | Rbm15 | | NaN | 0 | 0 | 0 | 0 | 0 | 0 |
| Q2EG98-7;Q2EG98-6;Q2EG9 | Pkd1l3 | Polycystic kidney disease protein 1-like 3 | NaN | 0 | 0 | 0 | 0 | 0 | 0 |
| Q3TBT3-3;Q3TBT3;Q3TBT3-: | Tmem173 | Stimulator of interferon genes protein | NaN | 0 | 0 | 0 | 0 | 0 | 0 |
| Q3TC93;A0A1W2P6R8 | Hs1bp3 | HCLS1-binding protein 3 | NaN | 0 | 0 | 0 | 0 | 0 | 0 |
| Q3TCN2 | Plbd2 | ase B-like 2 28 kDa form;Putative phospholipase B-like 2 40 kDa form;Putative phosph | NaN | 0 | 0 | 0 | 0 | 0 | 0 |
| Q3TFP0;Q9R0U0-3;Q9R0U0- | Srsf10 | Serine/arginine-rich splicing factor 10 | NaN | 0 | 0 | 4623900 | 616970 | 0 | 0 |
| Q6ZWQ5;Q3TGS7;Q3V2H3;C | Snx12 | Sorting nexin-12 | NaN | 0 | 0 | 0 | 0 | 0 | 0 |
| Q3THK3 | Gtf2f1 | General transcription factor IIF subunit 1 | NaN | 0 | 0 | 0 | 0 | 0 | 0 |
| Q3TIV5-2;Q3TIV5 | Zc3h15 | Zinc finger CCCH domain-containing protein 15 | NaN | 0 | 0 | 0 | 0 | 0 | 0 |
| Q3TQV3;Q8JZR2;Q5ND50;Q6 | Crk | Adapter molecule crk | NaN | 0 | 0 | 0 | 0 | 0 | 0 |
| Q8BPW9;Q3TVK3;Q9Z2W0 | Dnpep | Aspartyl aminopeptidase | NaN | 0 | 0 | 0 | 0 | 0 | 0 |
| Q3TW96;Q3TW96-2 | Uap1l1 | UDP-N-acetylhexosamine pyrophosphorylase-like protein 1 | NaN | 0 | 0 | 1667200 | 617300 | 0 | 0 |
| Q3TXS7;J3QN38 | Psmd1 | 26S proteasome non-ATPase regulatory subunit 1 | NaN | 0 | 0 | 0 | 0 | 0 | 0 |
| Q3TYX2 | Lrrn4cl | LRRN4 C-terminal-like protein | NaN | 0 | 0 | 0 | 0 | 0 | 0 |

| | | | | | | | | | |
|---|---|---|---|---|---|---|---|---|---|
| Q3U0S6 | Rasip1 | Ras-interacting protein 1 | NaN | 0 | 0 | 0 | 0 | 0 | 0 |
| Q3U0V2 | Tradd | or necrosis factor receptor type 1-associated DEATH domain protein | NaN | 0 | 0 | 0 | 0 | 0 | 0 |
| Q3U1Z5-2;Q3U1Z5;G3UYL4 | Gpsm3 | G-protein-signaling modulator 3 | NaN | 0 | 0 | 0 | 0 | 0 | 0 |
| Q3U367;Q9JLJ2 | Aldh9a1 | 4-trimethylaminobutyraldehyde dehydrogenase | NaN | 0 | 0 | 0 | 0 | 0 | 0 |
| Q3U4F0;Q91V61-2;Q91V61;A | Sfxn3 | Sideroflexin-3 | NaN | 0 | 0 | 868620 | 1645800 | 0 | 0 |
| Q3U741;Q501J6;Q3TU25;Q5 | Ddx17 | Probable ATP-dependent RNA helicase DDX17 | NaN | 0 | 0 | 0 | 0 | 0 | 0 |
| Q3U7R1;Q3U7R1-2;A0A1W2 | Esyt1 | Extended synaptotagmin-1 | NaN | 0 | 0 | 0 | 0 | 0 | 0 |
| Q3U9G9;A0A0A6YY12 | Lbr | Lamin-B receptor | NaN | 0 | 0 | 0 | 0 | 0 | 0 |
| Q3UDD3;Q8BG81;F6VR84 | Poldip3 | Polymerase delta-interacting protein 3 | NaN | 0 | 0 | 0 | 0 | 0 | 0 |
| Q3UDE2 | Ttll12 | Tubulin--tyrosine ligase-like protein 12 | NaN | 0 | 0 | 0 | 0 | 0 | 0 |
| Q3UE92;Q6P1B1;A0A494BB( | Xpnpep1 | Xaa-Pro aminopeptidase 1 | NaN | 0 | 0 | 1266500 | 1080200 | 0 | 0 |
| Q3UF75;Q9EPC1 | Parva | Alpha-parvin | NaN | 0 | 0 | 0 | 0 | 0 | 0 |
| Q3UGS4 | Fam195b | Protein FAM195B | NaN | 0 | 0 | 0 | 0 | 0 | 0 |
| Q5SV64;Q3UH59;Q61879 | Myh10 | Myosin-10 | NaN | 0 | 0 | 0 | 0 | 0 | 0 |
| Q3UH93 | Plxnd1 | Plexin-D1 | NaN | 0 | 0 | 0 | 0 | 0 | 0 |
| Q3UHG5;Q62283 | Tspan7 | Tetraspanin;Tetraspanin-7 | NaN | 0 | 0 | 0 | 0 | 0 | 0 |
| Q3UHK1 | Slc2a13 | Proton myo-inositol cotransporter | NaN | 0 | 0 | 0 | 0 | 0 | 0 |
| Q3UHX2 | Pdap1 | 28 kDa heat- and acid-stable phosphoprotein | NaN | 0 | 0 | 0 | 0 | 0 | 0 |
| Q3UJQ9;Q9D0K2 | Oxct1 | oenzyme A transferase;Succinyl-CoA:3-ketoacid coenzyme A transferase 1, mitochond | NaN | 0 | 0 | 0 | 0 | 0 | 0 |
| Q6PD20;Q3ULD5 | Mccc2 | Methylcrotonoyl-CoA carboxylase beta chain, mitochondrial | NaN | 0 | 0 | 0 | 0 | 0 | 0 |
| Q3UM45;A0A087WRA7 | Ppp1r7 | Protein phosphatase 1 regulatory subunit 7 | NaN | 664150 | 9758.1 | 0 | 0 | 0 | 0 |
| Q3UMT1;F6XWD4 | Ppp1r12c | Protein phosphatase 1 regulatory subunit 12C | NaN | 0 | 0 | 0 | 0 | 0 | 0 |
| S4R2A9;Q3UPL0-2;Q3UPL0;S | Sec31a | Protein transport protein Sec31A | NaN | 0 | 0 | 3466700 | 5565100 | 0 | 0 |
| Q3URS9-2;Q3URS9 | Ccdc51 | Coiled-coil domain-containing protein 51 | NaN | 0 | 0 | 0 | 0 | 0 | 0 |
| Q3V038 | Ttc9 | Tetratricopeptide repeat protein 9A | NaN | 0 | 0 | 0 | 0 | 0 | 0 |
| Q3V117;Q91V92;Q3TS02 | Acly | ATP-citrate synthase | NaN | 0 | 0 | 0 | 0 | 0 | 0 |
| Q3V3R1 | Mthfd1l | Monofunctional C1-tetrahydrofolate synthase, mitochondrial | NaN | 0 | 0 | 0 | 0 | 0 | 0 |
| Q4PZA2-3;Q4PZA2-2;Q4PZA | Ece1 | Endothelin-converting enzyme 1 | NaN | 0 | 0 | 0 | 0 | 0 | 0 |
| Q542V3;Q8VE97 | Srsf4 | Serine/arginine-rich splicing factor 4 | NaN | 0 | 0 | 0 | 0 | 0 | 0 |
| Q58A65-5;Q58A65-2;Q58A6 | Spag9 | C-Jun-amino-terminal kinase-interacting protein 4 | NaN | 0 | 0 | 0 | 0 | 0 | 0 |
| Q5DP50 | Tex24 | | NaN | 0 | 0 | 0 | 0 | 0 | 0 |
| Q5F258;Q68FF6 | Git1 | ARF GTPase-activating protein GIT1 | NaN | 0 | 0 | 0 | 0 | 0 | 0 |
| Q5F284;Q9JIZ9 | Plscr3 | Phospholipid scramblase 3 | NaN | 0 | 0 | 0 | 0 | 0 | 0 |
| Q5SQB0;Q61937;Q9DAY9;Q5 | Npm1 | Nucleophosmin | NaN | 0 | 0 | 0 | 0 | 0 | 0 |
| Q5SXA5;Q5SRX1-3;Q5SRX1-4 | Tom1l2 | TOM1-like protein 2 | NaN | 0 | 0 | 0 | 0 | 0 | 0 |
| Q5SUH7;Q5SUH6;Q99KN9-2 | Clint1 | Clathrin interactor 1 | NaN | 0 | 0 | 0 | 0 | 0 | 0 |
| Q5SUR0 | Pfas | Phosphoribosylformylglycinamidine synthase | NaN | 0 | 0 | 0 | 0 | 0 | 0 |
| Q5SUT0;Q5SUS9;Q61545;Q5 | Ewsr1 | RNA-binding protein EWS | NaN | 0 | 0 | 0 | 0 | 0 | 0 |
| Q5SWZ5;F6S5I0;F6RND9;P9 | Mprip | Myosin phosphatase Rho-interacting protein | NaN | 0 | 0 | 1174500 | 3636900 | 0 | 0 |
| Q5SX75;Q60716-2;Q60716;B | P4ha2 | Prolyl 4-hydroxylase subunit alpha-2 | NaN | 0 | 0 | 0 | 0 | 0 | 0 |
| Q5U458;E9Q8B3 | Dnajc11 | DnaJ homolog subfamily C member 11 | NaN | 0 | 0 | 0 | 0 | 0 | 0 |
| Q5U4C5;Q8VI75 | Ipo4 | Importin-4 | NaN | 0 | 0 | 0 | 0 | 0 | 0 |
| Q5U4D9 | Thoc6 | THO complex subunit 6 homolog | NaN | 0 | 0 | 0 | 0 | 0 | 0 |
| Q60876 | Eif4ebp1 | Eukaryotic translation initiation factor 4E-binding protein 1 | NaN | 0 | 0 | 0 | 0 | 0 | 0 |
| Q61001 | Lama5 | Laminin subunit alpha-5 | NaN | 0 | 0 | 0 | 0 | 0 | 0 |

| | | | | | | | | | |
|---|---|---|---|---|---|---|---|---|---|
| Q61033;Q61033-2;Q61029-4 | Tmpo | 2, isoforms alpha/zeta;Lamina-associated polypeptide 2, isoforms beta/delta/epsilon | NaN | 0 | 0 | 0 | 0 | 0 | 0 |
| Q61140-2;Q61140 | Bcar1 | Breast cancer anti-estrogen resistance protein 1 | NaN | 0 | 0 | 0 | 0 | 0 | 0 |
| Q61160 | Fadd | FAS-associated death domain protein | NaN | 0 | 0 | 0 | 0 | 0 | 0 |
| Q61166 | Mapre1 | Microtubule-associated protein RP/EB family member 1 | NaN | 0 | 0 | 0 | 0 | 0 | 0 |
| Q61249 | Igbp1 | Immunoglobulin-binding protein 1 | NaN | 0 | 0 | 0 | 0 | 0 | 0 |
| Q61292 | Lamb2 | Laminin subunit beta-2 | NaN | 0 | 0 | 0 | 0 | 0 | 0 |
| Q61550 | Rad21 | Double-strand-break repair protein rad21 homolog | NaN | 0 | 0 | 0 | 0 | 0 | 0 |
| Q61753;F6ZSB7 | Phgdh | D-3-phosphoglycerate dehydrogenase | NaN | 0 | 0 | 0 | 0 | 0 | 0 |
| Q61768;E9QAK5 | Kif5b | Kinesin-1 heavy chain;Kinesin-like protein | NaN | 0 | 0 | 0 | 0 | 0 | 0 |
| Q61881;D3Z6N3 | Mcm7 | DNA replication licensing factor MCM7 | NaN | 0 | 0 | 0 | 0 | 0 | 0 |
| Q62093 | Srsf2 | Serine/arginine-rich splicing factor 2 | NaN | 0 | 0 | 0 | 0 | 0 | 0 |
| Q62095 | Ddx3y | ATP-dependent RNA helicase DDX3Y | NaN | 0 | 0 | 817250 | 469520 | 0 | 0 |
| Q62159;A0A0G2JEP8;H3BL5( | Rhoc;Rhoa;4930544G11Rik;Rhob | ling protein RhoC;Transforming protein RhoA;Rho-related GTP-binding protein RhoB | NaN | 0 | 0 | 0 | 0 | 0 | 0 |
| Q9D8L3;Q62186 | Ssr4 | Translocon-associated protein subunit delta | NaN | 0 | 0 | 1217200 | 1294600 | 0 | 0 |
| Q62312-2;Q62312 | Tgfbr2 | TGF-beta receptor type-2 | NaN | 0 | 0 | 0 | 0 | 0 | 0 |
| Q8C872;Q62351 | Tfrc | Transferrin receptor protein 1 | NaN | 0 | 0 | 0 | 0 | 0 | 0 |
| Q62376;A0A1B0GSX5;A0A1E | Snrnp70 | U1 small nuclear ribonucleoprotein 70 kDa | NaN | 0 | 0 | 0 | 0 | 0 | 0 |
| Q62433;E9Q514 | Ndrg1 | Protein NDRG1 | NaN | 6981300 | 0 | 0 | 0 | 0 | 0 |
| Q62470;Q62470-2;Q62470-3 | Itga3 | rin alpha-3;Integrin alpha-3 heavy chain;Integrin alpha-3 light chain | NaN | 0 | 0 | 0 | 0 | 0 | 0 |
| Q7TQE2;Q62523;A0A0N4SVI | Zyx | Zyxin | NaN | 0 | 0 | 0 | 0 | 0 | 0 |
| Q63829 | Commd3 | COMM domain-containing protein 3 | NaN | 0 | 0 | 0 | 0 | 0 | 0 |
| Q64152-2;Q64152 | Btf3 | Transcription factor BTF3 | NaN | 0 | 0 | 0 | 0 | 0 | 0 |
| Q64282 | Ifit1 | Interferon-induced protein with tetratricopeptide repeats 1 | NaN | 0 | 0 | 0 | 0 | 0 | 0 |
| Q64387;Q64387-2 | Pnoc | Prepronociceptin;Neuropeptide 1;Nociceptin;Neuropeptide 2 | NaN | 0 | 0 | 0 | 0 | 13336000 | 1109700 |
| Q64669 | Nqo1 | NAD(P)H dehydrogenase [quinone] 1 | NaN | 0 | 0 | 0 | 0 | 0 | 0 |
| Q64727;A0A286YDJ4 | Vcl | Vinculin | NaN | 0 | 0 | 0 | 0 | 0 | 0 |
| Q64735-2;Q64735 | Cr1l | Complement component receptor 1-like protein | NaN | 0 | 0 | 0 | 0 | 0 | 0 |
| Q64737;Q64737-2 | Gart | oribosylamine--glycine ligase;Phosphoribosylformylglycinamidine cyclo-ligase;Phospho | NaN | 0 | 0 | 0 | 0 | 0 | 0 |
| Q6A0A2-2;Q6A0A2 | Larp4b | La-related protein 4B | NaN | 0 | 0 | 0 | 0 | 0 | 0 |
| Q6A0A9 | FAM120A | Constitutive coactivator of PPAR-gamma-like protein 1 | NaN | 0 | 0 | 0 | 0 | 0 | 0 |
| Q6DFV5;Q6DFV5-3 | Helz | Probable helicase with zinc finger domain | NaN | 0 | 0 | 0 | 0 | 1446900 | 1063900 |
| Q6DVA0 | Lemd2 | LEM domain-containing protein 2 | NaN | 0 | 0 | 0 | 0 | 0 | 0 |
| Q6GQT9 | Nomo1 | Nodal modulator 1 | NaN | 0 | 0 | 0 | 0 | 0 | 0 |
| Q6P4T2 | Snrnp200 | U5 small nuclear ribonucleoprotein 200 kDa helicase | NaN | 0 | 0 | 0 | 0 | 0 | 0 |
| Q6P542 | Abcf1 | ATP-binding cassette sub-family F member 1 | NaN | 0 | 0 | 0 | 0 | 0 | 0 |
| Q6P5B5;Q9WVR4 | Fxr2 | Fragile X mental retardation syndrome-related protein 2 | NaN | 0 | 0 | 1846400 | 1001900 | 0 | 0 |
| Q6P5F7-2;Q6P5F7 | Ttyh3 | Protein tweety homolog 3 | NaN | 0 | 0 | 0 | 0 | 0 | 0 |
| Q6P5H2-2;Q6P5H2 | Nes | Nestin | NaN | 0 | 0 | 0 | 0 | 0 | 0 |
| Q6P9J5 | Kank4 | KN motif and ankyrin repeat domain-containing protein 4 | NaN | 0 | 0 | 0 | 0 | 0 | 0 |
| Q6PAM1;A8Y5J8;A2ADZ3;A2 | Txlna | Alpha-taxilin | NaN | 0 | 0 | 1175400 | 1468100 | 0 | 0 |
| Q6PB44-2;Q6PB44 | Ptpn23 | Tyrosine-protein phosphatase non-receptor type 23 | NaN | 0 | 0 | 0 | 0 | 0 | 0 |
| Q6PB66 | Lrpprc | Leucine-rich PPR motif-containing protein, mitochondrial | NaN | 0 | 0 | 0 | 0 | 0 | 0 |
| Q6PD28 | Ppp2r5b | | NaN | 0 | 0 | 0 | 0 | 0 | 0 |
| Q6PF96;Q921G7 | Etfdh | ron transfer flavoprotein-ubiquinone oxidoreductase, mitochondrial | NaN | 0 | 0 | 0 | 0 | 0 | 0 |
| Q6PGH2 | Hn1l | Hematological and neurological expressed 1-like protein | NaN | 0 | 0 | 0 | 0 | 0 | 0 |

| | | | | | | | | | |
|---|---|---|---|---|---|---|---|---|---|
| Q6PGL7-2;Q6PGL7 | Fam21 | WASH complex subunit FAM21 | NaN | 0 | 0 | 0 | 0 | 0 | 0 |
| Q6R891-2;Q6R891 | Ppp1r9b | Neurabin-2 | NaN | 0 | 0 | 0 | 0 | 0 | 0 |
| Z4YJT3;Q6ZQ58;Q6ZQ58-2 | Larp1 | La-related protein 1 | NaN | 0 | 0 | 0 | 0 | 0 | 0 |
| Q6ZQ73 | Cand2 | Cullin-associated NEDD8-dissociated protein 2 | NaN | 0 | 0 | 647670 | 1505100 | 0 | 0 |
| Q6ZWM4;A0A0N4SUZ3 | Lsm8 | U6 snRNA-associated Sm-like protein LSm8 | NaN | 0 | 0 | 0 | 0 | 0 | 0 |
| Q6ZWY8 | Tmsb10 | Thymosin beta-10 | NaN | 4753000 | 866680 | 0 | 0 | 0 | 0 |
| Q6ZWZ4;P47964 | Rpl36 | 60S ribosomal protein L36 | NaN | 0 | 0 | 0 | 0 | 0 | 0 |
| Q70IV5-2;Q70IV5;A0A140LJ7 | Synm | Synemin | NaN | 0 | 0 | 0 | 0 | 0 | 0 |
| Q76MZ3;H3BIV7;H3BLQ0;H3 | Ppp2r1a;Ppp2r1b | ulatory subunit A alpha isoform;Serine/threonine-protein phosphatase 2A 65 kDa regu | NaN | 0 | 0 | 343620 | 293520 | 0 | 0 |
| Q7TQH0-2;Q3TGG2;A0A0U1 | Atxn2l | Ataxin-2-like protein | NaN | 0 | 0 | 0 | 0 | 0 | 0 |
| Q7TRE1;Q7TRE0 | Olfr884;Olfr885 | | NaN | 0 | 0 | 0 | 0 | 0 | 0 |
| Q7TSV4 | Pgm2 | Phosphoglucomutase-2 | NaN | 0 | 0 | 0 | 0 | 0 | 0 |
| Q7TT50;A0A1Y7VM95 | Cdc42bpb | Serine/threonine-protein kinase MRCK beta | NaN | 0 | 0 | 0 | 0 | 1519200 | 1296900 |
| Q80UU9 | Pgrmc2 | Membrane-associated progesterone receptor component 2 | NaN | 0 | 0 | 0 | 0 | 0 | 0 |
| Q80VP0-2;Q80VP0 | Tecpr1 | Tectonin beta-propeller repeat-containing protein 1 | NaN | 0 | 0 | 0 | 0 | 0 | 0 |
| Q80XI4 | Pip4k2b | Phosphatidylinositol 5-phosphate 4-kinase type-2 beta | NaN | 0 | 0 | 0 | 0 | 0 | 0 |
| Q80YX1-2;Q80YX1;Q80YX1-5 | Tnc | Tenascin | NaN | 0 | 0 | 0 | 0 | 0 | 0 |
| Q80ZA0 | Itln1b | Intelectin-1b | NaN | 0 | 0 | 0 | 0 | 0 | 0 |
| Q80ZE3 | Siglecg | | NaN | 0 | 0 | 0 | 0 | 0 | 0 |
| Q80ZS3 | Mrps26 | 28S ribosomal protein S26, mitochondrial | NaN | 0 | 0 | 0 | 0 | 0 | 0 |
| Q80ZX0;F6VJC5 | Sec24b | | NaN | 0 | 0 | 0 | 0 | 0 | 0 |
| Q810B6 | Ankfy1 | Rabankyrin-5 | NaN | 0 | 0 | 0 | 0 | 0 | 0 |
| Q8BFQ8 | Pddc1 | Parkinson disease 7 domain-containing protein 1 | NaN | 0 | 0 | 0 | 0 | 0 | 0 |
| Q8BFR4 | Gns | N-acetylglucosamine-6-sulfatase | NaN | 0 | 0 | 0 | 0 | 0 | 0 |
| Q8BFZ3 | Actbl2 | Beta-actin-like protein 2 | NaN | 0 | 0 | 0 | 0 | 0 | 0 |
| Q8BG32;G3UZ28;G3UYI4;G3 | Psmd11 | 26S proteasome non-ATPase regulatory subunit 11 | NaN | 0 | 0 | 0 | 0 | 0 | 0 |
| Q8BGT1 | Flrt3 | | NaN | 0 | 0 | 0 | 0 | 0 | 0 |
| Q8BH43;B1AUN0 | Wasf2 | Wiskott-Aldrich syndrome protein family member 2 | NaN | 0 | 0 | 0 | 0 | 0 | 0 |
| Q8BHC7 | Rhbdd1 | Rhomboid-related protein 4 | NaN | 0 | 0 | 0 | 0 | 0 | 0 |
| Q8BHL8 | Psmf1 | Proteasome inhibitor PI31 subunit | NaN | 0 | 0 | 0 | 0 | 0 | 0 |
| Q8BHN3;Q8BHN3-2;Q8BHN3 | Ganab | Neutral alpha-glucosidase AB | NaN | 0 | 0 | 0 | 0 | 0 | 0 |
| Q8BII1 | Prox2 | Prospero homeobox protein 2 | NaN | 0 | 769140 | 0 | 0 | 0 | 0 |
| Q8BIJ7 | Rufy1 | RUN and FYVE domain-containing protein 1 | NaN | 0 | 0 | 0 | 0 | 0 | 0 |
| Q8BJ05-2;Q8BJ05-3;Q8BJ05 | Zc3h14 | Zinc finger CCCH domain-containing protein 14 | NaN | 0 | 0 | 0 | 0 | 0 | 0 |
| Q8BJS4-3;Q8BJS4-2;Q8BJS4 | Sun2 | SUN domain-containing protein 2 | NaN | 0 | 0 | 0 | 0 | 0 | 0 |
| Q8BJY1;F7BA91 | Psmd5 | 26S proteasome non-ATPase regulatory subunit 5 | NaN | 0 | 0 | 0 | 0 | 0 | 0 |
| Q8BK64 | Ahsa1 | Activator of 90 kDa heat shock protein ATPase homolog 1 | NaN | 0 | 0 | 0 | 0 | 0 | 0 |
| Q8BL66 | Eea1 | Early endosome antigen 1 | NaN | 0 | 0 | 0 | 0 | 0 | 0 |
| Q8BMF4 | Dlat | due acetyltransferase component of pyruvate dehydrogenase complex, mitochondrial | NaN | 0 | 0 | 0 | 0 | 0 | 0 |
| Q8BN82-3;Q8BN82-2;Q8BN8 | Slc17a5 | Sialin | NaN | 0 | 0 | 0 | 0 | 0 | 0 |
| Q8BPG6 | Sumf2 | Sulfatase-modifying factor 2 | NaN | 0 | 0 | 0 | 0 | 0 | 0 |
| Q8BPU7-3;Q8BPU7 | Elmo1 | Engulfment and cell motility protein 1 | NaN | 0 | 0 | 0 | 0 | 0 | 0 |
| Q8BQ47;D3Z0T5 | Cnpy4 | Protein canopy homolog 4 | NaN | 0 | 0 | 0 | 0 | 0 | 0 |
| Q8BT60;Q9D6C8;A0A0R4J1D | ;Cpne4;Cpne6;Cpne2;Cpne9;Cpne7;Cpne8; | e-3;Copine-2;Copine-9;Copine-6;Copine-4;Copine-7;Copine-8;Copine-5 | NaN | 0 | 0 | 0 | 0 | 0 | 0 |
| Q8BTI8-3;Q8BTI8-2;Q8BTI8;/ | Srrm2 | Serine/arginine repetitive matrix protein 2 | NaN | 0 | 0 | 0 | 0 | 0 | 0 |

| | | | | | | | | | |
|---|---|---|---|---|---|---|---|---|---|
| Q8BTW3 | Exosc6 | Exosome complex component MTR3 | NaN | 0 | 0 | 0 | 0 | 0 | 0 |
| Q8BTZ7 | Gmppb | Mannose-1-phosphate guanyltransferase beta | NaN | 0 | 0 | 0 | 0 | 0 | 0 |
| Q8BU62 | Atp10d | | NaN | 0 | 0 | 0 | 0 | 0 | 0 |
| Q8BUE4;Q8BUE4-2 | Aifm2 | Apoptosis-inducing factor 2 | NaN | 0 | 0 | 1473500 | 1329600 | 0 | 0 |
| Q8BUK6;F6W1A2 | Hook3 | Protein Hook homolog 3 | NaN | 0 | 0 | 0 | 0 | 0 | 0 |
| Q8BUR4 | Dock1 | Dedicator of cytokinesis protein 1 | NaN | 0 | 0 | 0 | 0 | 0 | 0 |
| Q8BVA0;Q99P91 | Gpnmb | Transmembrane glycoprotein NMB | NaN | 0 | 0 | 0 | 0 | 0 | 0 |
| Q99MI6;Q8BWF2 | Gimap3;Gimap5 | GTPase IMAP family member 3;GTPase IMAP family member 5 | NaN | 0 | 0 | 0 | 0 | 0 | 0 |
| Q8BYZ1 | Abi3 | ABI gene family member 3 | NaN | 0 | 0 | 0 | 0 | 0 | 0 |
| Q8BZR9 | | Uncharacterized protein C17orf85 homolog | NaN | 0 | 0 | 0 | 0 | 0 | 0 |
| Q8C0E3;Q8C0E3-2 | Trim47 | Tripartite motif-containing protein 47 | NaN | 0 | 0 | 0 | 0 | 0 | 0 |
| Q8C2Q3;E9QL13;Q8C2Q3-2;J | Rbm14 | RNA-binding protein 14 | NaN | 0 | 0 | 1485500 | 4890600 | 0 | 0 |
| Q8C3W1 | | Uncharacterized protein C1orf198 homolog | NaN | 0 | 0 | 0 | 0 | 0 | 0 |
| Q8C3X8 | Lmf2 | Lipase maturation factor 2 | NaN | 0 | 0 | 0 | 0 | 0 | 0 |
| Q8C522 | Endod1 | Endonuclease domain-containing 1 protein | NaN | 0 | 0 | 0 | 0 | 0 | 0 |
| Q8C7E9;A2AEJ8;F6ZKC7;A2A | Cstf2t;Cstf2 | mulation factor subunit 2 tau variant;Cleavage stimulation factor subunit 2 | NaN | 264860 | 4266000 | 0 | 0 | 0 | 0 |
| Q8C8U0-2;Q8C8U0;Q8C8U0- | Ppfibp1;Ppfibp2 | Liprin-beta-1;Liprin-beta-2 | NaN | 0 | 0 | 0 | 0 | 0 | 0 |
| Q8CBY8-2;Q8CBY8 | Dctn4 | Dynactin subunit 4 | NaN | 0 | 0 | 0 | 0 | 0 | 0 |
| Q8CG76 | Akr7a2 | Aflatoxin B1 aldehyde reductase member 2 | NaN | 0 | 0 | 0 | 0 | 0 | 0 |
| Q8CGA0 | Ppm1f | Protein phosphatase 1F | NaN | 0 | 0 | 0 | 0 | 0 | 0 |
| Q8CGC7;A0A0A6YWH3;A0A0 | Eprs | lutamate/proline--tRNA ligase;Glutamate--tRNA ligase;Proline--tRNA ligase | NaN | 0 | 0 | 0 | 0 | 0 | 0 |
| Q8CGK3 | Lonp1 | Lon protease homolog, mitochondrial | NaN | 0 | 0 | 0 | 0 | 0 | 0 |
| Q8CHT0 | Aldh4a1 | Delta-1-pyrroline-5-carboxylate dehydrogenase, mitochondrial | NaN | 0 | 0 | 0 | 0 | 0 | 0 |
| Q8CHU3;F7CD65;F7CUV7;Q5 | Epn2 | Epsin-2 | NaN | 0 | 0 | 0 | 0 | 0 | 0 |
| Q8CI08;Q8CI08-2 | Slain2 | SLAIN motif-containing protein 2 | NaN | 0 | 0 | 0 | 0 | 0 | 0 |
| Q8JZK9 | Hmgcs1 | Hydroxymethylglutaryl-CoA synthase, cytoplasmic | NaN | 0 | 0 | 0 | 0 | 0 | 0 |
| Q8K078-2;Q8K078 | Slco4a1 | Solute carrier organic anion transporter family member 4A1 | NaN | 0 | 0 | 0 | 0 | 0 | 0 |
| Q8K0B2-3;Q8K0B2-2;Q8K0B2 | Lmbrd1 | Probable lysosomal cobalamin transporter | NaN | 0 | 0 | 0 | 0 | 0 | 0 |
| Q8K0H5 | Taf10 | Transcription initiation factor TFIID subunit 10 | NaN | 0 | 0 | 0 | 0 | 0 | 0 |
| Q8K2C7-2;Q8K2C7 | Os9 | Protein OS-9 | NaN | 0 | 0 | 0 | 0 | 0 | 0 |
| Q8K2Y3 | Eva1b | Protein eva-1 homolog B | NaN | 0 | 0 | 0 | 0 | 0 | 0 |
| Q8K3J1 | Ndufs8 | H dehydrogenase [ubiquinone] iron-sulfur protein 8, mitochondrial | NaN | 0 | 0 | 0 | 0 | 0 | 0 |
| Q8QZT1 | Acat1 | Acetyl-CoA acetyltransferase, mitochondrial | NaN | 0 | 0 | 0 | 0 | 0 | 0 |
| Q8QZY1 | Eif3l | Eukaryotic translation initiation factor 3 subunit L | NaN | 0 | 0 | 0 | 0 | 0 | 0 |
| Q8R010 | Aimp2 | noacyl tRNA synthase complex-interacting multifunctional protein 2 | NaN | 0 | 0 | 0 | 0 | 0 | 0 |
| Q8R059 | Gale | UDP-glucose 4-epimerase | NaN | 0 | 0 | 0 | 0 | 0 | 0 |
| Q8R0G9 | Nup133 | Nuclear pore complex protein Nup133 | NaN | 0 | 0 | 0 | 0 | 0 | 0 |
| Q8R0T6;A0A1D5RLS0 | Gpr97 | Probable G-protein coupled receptor 97 | NaN | 0 | 0 | 0 | 0 | 0 | 0 |
| Q8R1V4;Q99KF1 | Tmed4;Tmed9 | 24 domain-containing protein 4;Transmembrane emp24 domain-containing protein 9 | NaN | 0 | 0 | 0 | 0 | 0 | 0 |
| Q8R2Q8 | Bst2 | Bone marrow stromal antigen 2 | NaN | 0 | 0 | 0 | 0 | 0 | 0 |
| Q8R2U6 | Nudt4 | Diphosphoinositol polyphosphate phosphohydrolase 2 | NaN | 0 | 0 | 0 | 0 | 0 | 0 |
| Q8R307 | Vps18 | Vacuolar protein sorting-associated protein 18 homolog | NaN | 0 | 0 | 0 | 0 | 0 | 0 |
| Q8R317;Q8R317-2 | Ubqln1 | Ubiquilin-1 | NaN | 0 | 0 | 0 | 0 | 0 | 0 |
| Q8R4R6;A2ATJ2 | Nup35 | Nucleoporin NUP53 | NaN | 0 | 0 | 0 | 0 | 0 | 0 |
| Q8R570 | Snap47 | Synaptosomal-associated protein 47 | NaN | 0 | 0 | 0 | 0 | 0 | 0 |

| | | | | | | | | | |
|---|---|---|---|---|---|---|---|---|---|
| Q8VBV7;A0A087WPM5 | Cops8 | COP9 signalosome complex subunit 8 | NaN | 0 | 0 | 0 | 0 | 0 | 0 |
| Q8VC30 | Dak | one kinase/FAD-AMP lyase (cyclizing);ATP-dependent dihydroxyacetone kinase;FAD-A | NaN | 0 | 0 | 0 | 0 | 0 | 0 |
| Q8VCF0 | Mavs | Mitochondrial antiviral-signaling protein | NaN | 0 | 0 | 0 | 0 | 0 | 0 |
| Q9CQ43;Q8VCG1 | Dut | | NaN | 0 | 0 | 0 | 0 | 0 | 0 |
| Q8VCR4;Q9DCG9 | Trmt112 | Multifunctional methyltransferase subunit TRM112-like protein | NaN | 0 | 0 | 0 | 0 | 0 | 0 |
| Q8VCV2;Q9QYF9 | Ndrg3 | Protein NDRG3 | NaN | 0 | 0 | 0 | 0 | 0 | 0 |
| Q8VD75;A0A0J9YV08;A0A1D | Hip1 | Huntingtin-interacting protein 1 | NaN | 0 | 0 | 0 | 0 | 0 | 0 |
| Q8VDM4 | Psmd2 | 26S proteasome non-ATPase regulatory subunit 2 | NaN | 4565700 | 395720 | 0 | 0 | 0 | 0 |
| Q8VDP3-2;Q8VDP3-3;Q8VDP | Mical1 | Protein-methionine sulfoxide oxidase MICAL1 | NaN | 0 | 0 | 0 | 0 | 0 | 0 |
| Q8VE99 | Ccdc115 | Coiled-coil domain-containing protein 115 | NaN | 0 | 0 | 0 | 0 | 0 | 0 |
| Q8VEE1 | Lmcd1 | LIM and cysteine-rich domains protein 1 | NaN | 0 | 0 | 0 | 0 | 0 | 0 |
| Q91V12-2;Q91V12-4;E9PYH2 | Acot7 | Cytosolic acyl coenzyme A thioester hydrolase | NaN | 0 | 0 | 0 | 0 | 0 | 0 |
| Q91VD9 | Ndufs1 | NADH-ubiquinone oxidoreductase 75 kDa subunit, mitochondrial | NaN | 0 | 0 | 1268600 | 1530800 | 0 | 0 |
| Q91VH6;A0A3B2WCC1;A0A3 | Memo1 | Protein MEMO1 | NaN | 0 | 0 | 0 | 0 | 0 | 0 |
| Q91VR5;A0A1Y7VM48 | Ddx1 | ATP-dependent RNA helicase DDX1 | NaN | 696940 | 1019700 | 0 | 0 | 0 | 0 |
| Q91WC0-3;Q91WC0-2;Q91W | Setd3 | Histone-lysine N-methyltransferase setd3 | NaN | 0 | 0 | 0 | 0 | 0 | 0 |
| Q91WG2-2;Q91WG2-1;Q91\ | Rabep2 | Rab GTPase-binding effector protein 2 | NaN | 0 | 0 | 0 | 0 | 0 | 0 |
| Q91YP2 | Nln | Neurolysin, mitochondrial | NaN | 0 | 0 | 790550 | 325820 | 0 | 0 |
| Q91YQ5;A0A0N4SUJ8 | Rpn1 | nyl-diphosphooligosaccharide--protein glycosyltransferase subunit 1 | NaN | 0 | 0 | 0 | 0 | 0 | 0 |
| Q91YR1 | Twf1 | Twinfilin-1 | NaN | 0 | 0 | 0 | 0 | 0 | 0 |
| Q91YT8 | Tmem63a | CSC1-like protein 1 | NaN | 0 | 0 | 0 | 0 | 0 | 0 |
| Q91YW3 | Dnajc3 | DnaJ homolog subfamily C member 3 | NaN | 0 | 0 | 0 | 0 | 0 | 0 |
| Q91Z96-2;Q91Z96 | Bmp2k | BMP-2-inducible protein kinase | NaN | 609870 | 1025900 | 0 | 0 | 0 | 0 |
| Q91ZR1 | Rab4b | Ras-related protein Rab-4B | NaN | 0 | 0 | 0 | 0 | 0 | 0 |
| Q921I1 | Tf | Serotransferrin | NaN | 0 | 0 | 0 | 0 | 0 | 0 |
| Q921M3-2;Q921M3 | Sf3b3 | Splicing factor 3B subunit 3 | NaN | 0 | 0 | 0 | 0 | 0 | 0 |
| Q922F4 | Tubb6 | Tubulin beta-6 chain | NaN | 0 | 0 | 0 | 0 | 0 | 0 |
| Q922Q1 | 02-Mar | Mitochondrial amidoxime reducing component 2 | NaN | 0 | 0 | 0 | 0 | 0 | 0 |
| Q922Q4 | Pycr2 | Pyrroline-5-carboxylate reductase 2 | NaN | 0 | 0 | 0 | 0 | 0 | 0 |
| Q922Q8 | Lrrc59 | Leucine-rich repeat-containing protein 59 | NaN | 0 | 0 | 0 | 0 | 0 | 0 |
| Q923D5 | Wbp11 | WW domain-binding protein 11 | NaN | 0 | 0 | 0 | 0 | 0 | 0 |
| Q925B0-2;Q925B0 | Pawr | PRKC apoptosis WT1 regulator protein | NaN | 0 | 0 | 15850000 | 0 | 114600000 | 0 |
| Q925F2;D3Z5Y0 | Esam | Endothelial cell-selective adhesion molecule | NaN | 0 | 0 | 0 | 0 | 0 | 0 |
| Q99J36 | Thumpd1 | THUMP domain-containing protein 1 | NaN | 0 | 0 | 0 | 0 | 0 | 0 |
| Q99JB2;A2AG41;A2AG39;F6' | Stoml2 | Stomatin-like protein 2, mitochondrial | NaN | 1872300 | 585780 | 0 | 0 | 0 | 0 |
| Q99JP6;Q99JP6-2;J3QQ00 | Homer3 | Homer protein homolog 3 | NaN | 0 | 0 | 0 | 0 | 0 | 0 |
| Q99JR1 | Sfxn1 | Sideroflexin-1 | NaN | 0 | 0 | 0 | 0 | 0 | 0 |
| Q99JX4 | Eif3m | Eukaryotic translation initiation factor 3 subunit M | NaN | 0 | 0 | 0 | 0 | 0 | 0 |
| V9GX96;V9GXF3;Q99JX7 | Nxf1 | Nuclear RNA export factor 1 | NaN | 0 | 0 | 0 | 0 | 0 | 0 |
| Q99JY9;A0A087WRA1;A0A0 | Actr3 | Actin-related protein 3 | NaN | 0 | 0 | 0 | 0 | 0 | 0 |
| Q99K41 | Emilin1 | EMILIN-1 | NaN | 0 | 0 | 0 | 0 | 0 | 0 |
| Q99K48;Q99K48-2;B1AXT0 | Nono | Non-POU domain-containing octamer-binding protein | NaN | 0 | 0 | 0 | 0 | 0 | 0 |
| Q99KC8 | Vwa5a | von Willebrand factor A domain-containing protein 5A | NaN | 0 | 0 | 0 | 0 | 0 | 0 |
| Q99KE1 | Me2 | NAD-dependent malic enzyme, mitochondrial | NaN | 0 | 0 | 0 | 0 | 0 | 0 |
| Q99KI0;A0A2R8VJW0 | Aco2 | Aconitate hydratase, mitochondrial | NaN | 0 | 0 | 3828100 | 1474900 | 0 | 0 |

| | | | | | | | | | |
|---|---|---|---|---|---|---|---|---|---|
| Q99KJ8 | Dctn2 | Dynactin subunit 2 | NaN | 0 | 0 | 0 | 0 | 0 | 0 |
| Q99KK7;A0A494BBC1;A0A49 | Dpp3 | Dipeptidyl peptidase 3 | NaN | 0 | 0 | 0 | 0 | 0 | 0 |
| Q99KQ4 | Nampt | Nicotinamide phosphoribosyltransferase | NaN | 0 | 0 | 0 | 0 | 0 | 0 |
| Q99KV1 | Dnajb11 | DnaJ homolog subfamily B member 11 | NaN | 0 | 0 | 0 | 0 | 0 | 0 |
| Q99L45;E0CXJ3 | Eif2s2 | Eukaryotic translation initiation factor 2 subunit 2 | NaN | 0 | 0 | 831350 | 343090 | 0 | 0 |
| Q99LB4;P24452;D3YZN3;D3Y | Capg | Macrophage-capping protein | NaN | 0 | 0 | 1419900 | 1670100 | 0 | 0 |
| Q99LJ0 | Cttnbp2nl | CTTNBP2 N-terminal-like protein | NaN | 0 | 0 | 0 | 0 | 0 | 0 |
| Q99LN9;D3Z7J7;D3Z6Y9;Q99 | Dohh | Deoxyhypusine hydroxylase | NaN | 0 | 0 | 0 | 0 | 0 | 0 |
| Q99M08 | | Uncharacterized protein C4orf3 homolog | NaN | 0 | 0 | 0 | 0 | 0 | 0 |
| Q99M71-2;Q99M71 | Epdr1 | Mammalian ependymin-related protein 1 | NaN | 0 | 0 | 0 | 0 | 0 | 0 |
| Q99M87-3;Q99M87-2;Q99M | Dnaja3 | DnaJ homolog subfamily A member 3, mitochondrial | NaN | 0 | 0 | 0 | 0 | 0 | 0 |
| Q99MR3 | Slc12a9 | Solute carrier family 12 member 9 | NaN | 0 | 0 | 0 | 0 | 0 | 0 |
| Q99MR6-3;Q99MR6-4;Q99M | Srrt | Serrate RNA effector molecule homolog | NaN | 0 | 0 | 0 | 0 | 0 | 0 |
| Q99NB8 | Ubqln4 | Ubiquilin-4 | NaN | 0 | 0 | 0 | 0 | 0 | 0 |
| Q99P72-5;Q99P72-4;Q99P72 | Rtn4 | Reticulon-4 | NaN | 0 | 0 | 0 | 0 | 0 | 0 |
| Q99PV0;B7ZC27 | Prpf8 | Pre-mRNA-processing-splicing factor 8 | NaN | 0 | 0 | 0 | 0 | 0 | 0 |
| Q9CPP0 | Npm3 | Nucleoplasmin-3 | NaN | 0 | 0 | 0 | 0 | 0 | 0 |
| Q9CPQ3;A0A2R8VHM4 | Tomm22 | Mitochondrial import receptor subunit TOM22 homolog | NaN | 0 | 0 | 0 | 0 | 0 | 0 |
| Q9CPS7 | Pno1 | RNA-binding protein PNO1 | NaN | 0 | 0 | 0 | 0 | 0 | 0 |
| Q9CPY7-2;Q9CPY7;A0A0G2JE | Lap3 | Cytosol aminopeptidase | NaN | 0 | 0 | 0 | 0 | 0 | 0 |
| Q9CPZ6 | Ormdl3 | ORM1-like protein 3 | NaN | 0 | 0 | 0 | 0 | 0 | 0 |
| Q9CQ65 | Mtap | S-methyl-5-thioadenosine phosphorylase | NaN | 0 | 0 | 1983100 | 417110 | 0 | 0 |
| Q9CQ69 | Uqcrq | Cytochrome b-c1 complex subunit 8 | NaN | 0 | 0 | 0 | 0 | 0 | 0 |
| Q9CQ71 | Rpa3 | Replication protein A 14 kDa subunit | NaN | 0 | 0 | 0 | 0 | 0 | 0 |
| Q9CQE8 | | UPF0568 protein C14orf166 homolog | NaN | 0 | 0 | 0 | 0 | 3869800 | 505170 |
| Q9CQI6 | Cotl1 | Coactosin-like protein | NaN | 0 | 0 | 1017100 | 721460 | 0 | 0 |
| Q9CQI7 | Snrpb2 | U2 small nuclear ribonucleoprotein B | NaN | 0 | 0 | 0 | 0 | 0 | 0 |
| Q9CQN1 | Trap1 | Heat shock protein 75 kDa, mitochondrial | NaN | 0 | 0 | 0 | 0 | 0 | 0 |
| Q9CQQ4 | Gemin2 | Gem-associated protein 2 | NaN | 0 | 0 | 0 | 0 | 0 | 0 |
| Q9CQV4-2;Q9CQV4 | Fam134c | Protein FAM134C | NaN | 0 | 0 | 0 | 0 | 0 | 0 |
| Q9CR00;A0A0G2JGN6 | Psmd9 | 26S proteasome non-ATPase regulatory subunit 9 | NaN | 0 | 0 | 0 | 0 | 0 | 0 |
| Q9CR41 | Hypk | Huntingtin-interacting protein K | NaN | 0 | 0 | 0 | 0 | 0 | 0 |
| Q9CR57;A0A1L1SUF6 | Rpl14 | 60S ribosomal protein L14 | NaN | 0 | 0 | 0 | 0 | 0 | 0 |
| Q9CR86 | Carhsp1 | Calcium-regulated heat stable protein 1 | NaN | 0 | 0 | 0 | 0 | 0 | 0 |
| Q9CRD2 | Emc2 | ER membrane protein complex subunit 2 | NaN | 0 | 0 | 0 | 0 | 0 | 0 |
| Q9CU62 | Smc1a | Structural maintenance of chromosomes protein 1A | NaN | 0 | 0 | 0 | 0 | 3175400 | 0 |
| Q9CW03 | Smc3 | Structural maintenance of chromosomes protein 3 | NaN | 0 | 0 | 0 | 0 | 0 | 0 |
| Q9CWK3;A0A0U1RP07;A0A0 | Cd2bp2 | CD2 antigen cytoplasmic tail-binding protein 2 | NaN | 0 | 0 | 0 | 0 | 0 | 0 |
| Q9CWL8 | Ctnnbl1 | Beta-catenin-like protein 1 | NaN | 0 | 0 | 0 | 0 | 0 | 0 |
| Q9CX00 | Ist1 | IST1 homolog | NaN | 0 | 0 | 0 | 0 | 0 | 0 |
| Q9CX34 | Sugt1 | Suppressor of G2 allele of SKP1 homolog | NaN | 0 | 0 | 0 | 0 | 0 | 0 |
| Q9CX60;F6UVG6 | Lbh | Protein LBH | NaN | 0 | 0 | 0 | 0 | 0 | 0 |
| Q9CX86 | Hnrnpa0 | Heterogeneous nuclear ribonucleoprotein A0 | NaN | 0 | 0 | 0 | 0 | 0 | 0 |
| Q9CY97-2;Q9CY97 | Ssu72 | A polymerase II subunit A C-terminal domain phosphatase SSU72 | NaN | 0 | 0 | 0 | 0 | 0 | 0 |
| Q9CYG7;Q9CYG7-2 | Tomm34 | Mitochondrial import receptor subunit TOM34 | NaN | 0 | 0 | 0 | 0 | 0 | 0 |

| | | | | | | | | | |
|---|---|---|---|---|---|---|---|---|---|
| Q9CYH6 | Rrs1 | Ribosome biogenesis regulatory protein homolog | NaN | 0 | 0 | 0 | 0 | 0 | 0 |
| Q9CYL5 | Glipr2 | Golgi-associated plant pathogenesis-related protein 1 | NaN | 0 | 0 | 0 | 0 | 0 | 0 |
| Q9CYN9 | Atp6ap2 | Renin receptor | NaN | 0 | 0 | 0 | 0 | 0 | 0 |
| Q9CZ04;Q9CZ04-2 | Cops7a | COP9 signalosome complex subunit 7a | NaN | 0 | 0 | 0 | 0 | 0 | 0 |
| Q9CZ13;A0A0A6YWX6;A0A0. | Uqcrc1 | Cytochrome b-c1 complex subunit 1, mitochondrial | NaN | 996990 | 1142500 | 0 | 0 | 0 | 0 |
| Q9CZ44-2;Q9CZ44;Q9CZ44-3 | Nsfl1c | NSFL1 cofactor p47 | NaN | 0 | 0 | 0 | 0 | 0 | 0 |
| Q9ERR1-2;Q9CZA6-3;Q9CZA( | Ndel1;Nde1 | :tribution protein nudE-like 1;Nuclear distribution protein nudE homolog 1 | NaN | 0 | 0 | 0 | 0 | 0 | 0 |
| Q9CZC8;A0A0N4SWD0 | Scrn1 | Secernin-1 | NaN | 0 | 0 | 0 | 0 | 0 | 0 |
| Q9CZE3 | Rab32 | Ras-related protein Rab-32 | NaN | 0 | 0 | 0 | 0 | 0 | 0 |
| Q9CZJ2;H7BX84;Q8K0U4 | Hspa12b | Heat shock 70 kDa protein 12B | NaN | 0 | 0 | 0 | 0 | 0 | 0 |
| Q9CZR8 | Tsfm | Elongation factor Ts, mitochondrial | NaN | 0 | 0 | 0 | 0 | 0 | 0 |
| Q9CZW5 | Tomm70a | Mitochondrial import receptor subunit TOM70 | NaN | 0 | 0 | 0 | 0 | 0 | 0 |
| Q9D051 | Pdhb | ·uvate dehydrogenase E1 component subunit beta, mitochondrial | NaN | 0 | 0 | 0 | 0 | 0 | 0 |
| Q9D0F3 | Lman1 | Protein ERGIC-53 | NaN | 0 | 0 | 0 | 0 | 0 | 0 |
| Q9D0I9 | Rars | Arginine--tRNA ligase, cytoplasmic | NaN | 0 | 0 | 849870 | 2228000 | 0 | 0 |
| Q9D0J4 | Arl2 | ADP-ribosylation factor-like protein 2 | NaN | 0 | 0 | 0 | 0 | 0 | 0 |
| Q9D0L8-2;Q9D0L8 | Rnmt | mRNA cap guanine-N7 methyltransferase | NaN | 0 | 0 | 0 | 0 | 0 | 0 |
| Q9D0M3-2;Q9D0M3;A0A2R8 | Cyc1 | Cytochrome c1, heme protein, mitochondrial | NaN | 0 | 0 | 0 | 0 | 0 | 0 |
| Q9D0R2;A0A2I3BPK8 | Tars | Threonine--tRNA ligase, cytoplasmic | NaN | 0 | 0 | 1054000 | 4800500 | 0 | 0 |
| Q9D0S9 | Hint2 | Histidine triad nucleotide-binding protein 2, mitochondrial | NaN | 0 | 0 | 0 | 0 | 0 | 0 |
| Q9D1I2 | | Bcl10-interacting CARD protein | NaN | 0 | 0 | 0 | 0 | 0 | 0 |
| Q9D1M0 | Sec13 | Protein SEC13 homolog | NaN | 0 | 0 | 0 | 0 | 0 | 0 |
| Q9D1Q4 | Dpm3 | Dolichol-phosphate mannosyltransferase subunit 3 | NaN | 0 | 0 | 0 | 0 | 0 | 0 |
| Q9D1Q6 | Erp44 | Endoplasmic reticulum resident protein 44 | NaN | 0 | 0 | 0 | 0 | 0 | 0 |
| Q9D2G2-2;Q9D2G2 | Dlst | succinyltransferase component of 2-oxoglutarate dehydrogenase complex, mitochonc | NaN | 0 | 0 | 0 | 0 | 0 | 0 |
| Q9D2R0 | Aacs | Acetoacetyl-CoA synthetase | NaN | 0 | 0 | 0 | 0 | 0 | 0 |
| Q9D554 | Sf3a3 | Splicing factor 3A subunit 3 | NaN | 0 | 0 | 0 | 0 | 0 | 0 |
| Q9D662;A2ANA0;A2AN97;A2 | Sec23b | Protein transport protein Sec23B | NaN | 0 | 0 | 0 | 0 | 0 | 0 |
| Q9D6V8 | Paip2 | Polyadenylate-binding protein-interacting protein 2 | NaN | 0 | 0 | 0 | 0 | 0 | 0 |
| Q9D6Z1;F6USW7;F6U250;F6 | Nop56 | Nucleolar protein 56 | NaN | 0 | 0 | 1001300 | 1326900 | 0 | 0 |
| Q9D7E4 | | UPF0449 protein C19orf25 homolog | NaN | 0 | 0 | 0 | 0 | 0 | 0 |
| Q9D7S7-2;Q9D7S7 | Rpl22l1 | 60S ribosomal protein L22-like 1 | NaN | 0 | 0 | 0 | 0 | 0 | 0 |
| Q9D7S9 | Chmp5 | Charged multivesicular body protein 5 | NaN | 0 | 0 | 0 | 0 | 0 | 0 |
| Q9D819 | Ppa1 | Inorganic pyrophosphatase | NaN | 0 | 0 | 0 | 0 | 908640 | 875110 |
| Q9D880 | Timm50 | litochondrial import inner membrane translocase subunit TIM50 | NaN | 0 | 0 | 0 | 0 | 0 | 0 |
| Q9D883;A0A494B947;G3UW | U2af1 | Splicing factor U2AF 35 kDa subunit | NaN | 0 | 0 | 0 | 0 | 0 | 0 |
| Q9D8U8 | Snx5 | Sorting nexin-5 | NaN | 0 | 0 | 0 | 0 | 0 | 0 |
| Q9D8X2 | Ccdc124 | Coiled-coil domain-containing protein 124 | NaN | 0 | 0 | 0 | 0 | 0 | 0 |
| Q9D967 | Mdp1 | Magnesium-dependent phosphatase 1 | NaN | 0 | 0 | 0 | 0 | 0 | 0 |
| Q9DAK9 | Phpt1 | 14 kDa phosphohistidine phosphatase | NaN | 0 | 0 | 0 | 0 | 0 | 0 |
| Q9DB73 | Cyb5r1 | NADH-cytochrome b5 reductase 1 | NaN | 0 | 0 | 0 | 0 | 0 | 0 |
| Q9DB77;A0A140LI98 | Uqcrc2 | Cytochrome b-c1 complex subunit 2, mitochondrial | NaN | 0 | 0 | 0 | 0 | 0 | 0 |
| Q9DBC7;D3Z0V6;P12849 | Prkar1a;Prkar1b | ıdent protein kinase type I-alpha regulatory subunit, N-terminally processed;cAMP-dep | NaN | 0 | 0 | 0 | 0 | 0 | 0 |
| Q9DBG5;A0A3B2WCW2 | Plin3 | Perilipin-3 | NaN | 0 | 0 | 3910600 | 886150 | 0 | 0 |
| Q9DBH5 | Lman2 | Vesicular integral-membrane protein VIP36 | NaN | 0 | 0 | 0 | 0 | 0 | 0 |

| | | | | | | | | | |
|---|---|---|---|---|---|---|---|---|---|
| Q9DBL2;Q9DBL2-2 | Gdap2 | Ganglioside-induced differentiation-associated protein 2 | NaN | 0 | 0 | 0 | 1435000 | 0 | 1467600 |
| Q9DBR0 | Akap8 | A-kinase anchor protein 8 | NaN | 0 | 0 | 0 | 0 | 0 | 0 |
| Q9DBR1-2;Q9DBR1 | Xrn2 | 5-3 exoribonuclease 2 | NaN | 0 | 0 | 0 | 0 | 0 | 0 |
| Q9DBS5 | Klc4 | Kinesin light chain 4 | NaN | 0 | 0 | 0 | 0 | 0 | 0 |
| Q9DC51 | Gnai3 | Guanine nucleotide-binding protein G(k) subunit alpha | NaN | 0 | 0 | 0 | 0 | 0 | 0 |
| Q9DCC4 | Pycrl | Pyrroline-5-carboxylate reductase 3 | NaN | 0 | 0 | 0 | 0 | 0 | 0 |
| Q9DCD0 | Pgd | 6-phosphogluconate dehydrogenase, decarboxylating | NaN | 0 | 0 | 0 | 0 | 0 | 0 |
| Q9DCH4 | Eif3f | Eukaryotic translation initiation factor 3 subunit F | NaN | 0 | 0 | 0 | 0 | 0 | 0 |
| Q9DCS9;D3YUK4 | Ndufb10 | ADH dehydrogenase [ubiquinone] 1 beta subcomplex subunit 10 | NaN | 0 | 0 | 0 | 0 | 0 | 0 |
| Q9DCT2 | Ndufs3 | )H dehydrogenase [ubiquinone] iron-sulfur protein 3, mitochondrial | NaN | 0 | 0 | 0 | 0 | 0 | 0 |
| Q9DCW4;A0A0U1RNR3;A0A( | Etfb | Electron transfer flavoprotein subunit beta | NaN | 0 | 0 | 0 | 0 | 0 | 0 |
| Q9DD18 | Dtd1 | D-tyrosyl-tRNA(Tyr) deacylase 1 | NaN | 0 | 0 | 0 | 0 | 0 | 0 |
| Q9DD24 | Wbp5 | WW domain-binding protein 5 | NaN | 0 | 0 | 0 | 0 | 0 | 0 |
| Q9EPE9 | Atp13a1 | Manganese-transporting ATPase 13A1 | NaN | 0 | 0 | 0 | 0 | 0 | 0 |
| Q9EPL8 | Ipo7 | Importin-7 | NaN | 0 | 0 | 0 | 0 | 0 | 0 |
| Q9EQH3 | Vps35 | Vacuolar protein sorting-associated protein 35 | NaN | 0 | 0 | 0 | 0 | 0 | 0 |
| Q9EQP2 | Ehd4 | EH domain-containing protein 4 | NaN | 0 | 0 | 0 | 0 | 0 | 0 |
| Q9QX15;Q9EQR4;A0A0G2JG | Clca3a1;Clca3a2;Clca3b | | NaN | 0 | 0 | 0 | 0 | 0 | 0 |
| Q9ER00 | Stx12 | Syntaxin-12 | NaN | 598680 | 200340 | 0 | 0 | 0 | 0 |
| Q9ER05 | Ctrl | | NaN | 0 | 0 | 0 | 0 | 0 | 0 |
| Q9ER72-2;Q9ER72;A0A140LI | Cars | Cysteine--tRNA ligase, cytoplasmic | NaN | 0 | 0 | 0 | 0 | 0 | 0 |
| Q9ER73 | Elp4 | Elongator complex protein 4 | NaN | 0 | 0 | 0 | 0 | 0 | 0 |
| Q9ER81 | Tor1aip2 | Torsin-1A-interacting protein 2, isoform IFRG15 | NaN | 0 | 0 | 0 | 0 | 0 | 0 |
| Q9ERD7 | Tubb3 | Tubulin beta-3 chain | NaN | 0 | 0 | 0 | 0 | 0 | 0 |
| Q9ERG0-2;Q9ERG0 | Lima1 | LIM domain and actin-binding protein 1 | NaN | 0 | 0 | 0 | 0 | 0 | 0 |
| Q9ERN0 | Scamp2 | Secretory carrier-associated membrane protein 2 | NaN | 0 | 0 | 0 | 0 | 0 | 0 |
| Q9ET22 | Dpp7 | Dipeptidyl peptidase 2 | NaN | 0 | 0 | 0 | 0 | 0 | 0 |
| Q9JHF5;F6ZFB8 | Tcirg1 | V-type proton ATPase subunit a | NaN | 0 | 0 | 0 | 0 | 0 | 0 |
| Q9JHP7-3;Q9JHP7-2;Q9JHP7 | Kdelc1 | KDEL motif-containing protein 1 | NaN | 0 | 0 | 0 | 0 | 0 | 0 |
| Q9JHU9;A0A1B0GS86 | Isyna1 | Inositol-3-phosphate synthase 1 | NaN | 0 | 0 | 0 | 0 | 0 | 0 |
| Q9JIG7 | Ccdc22 | Coiled-coil domain-containing protein 22 | NaN | 0 | 0 | 0 | 0 | 0 | 0 |
| Q9JII6;B1AXW3 | Akr1a1 | Alcohol dehydrogenase [NADP(+)] | NaN | 0 | 0 | 0 | 0 | 0 | 0 |
| Q9JIK5 | Ddx21 | Nucleolar RNA helicase 2 | NaN | 0 | 0 | 0 | 0 | 0 | 0 |
| Q9JJ28 | Flii | Protein flightless-1 homolog | NaN | 0 | 0 | 0 | 0 | 0 | 0 |
| Q9JKF1 | Iqgap1 | Ras GTPase-activating-like protein IQGAP1 | NaN | 0 | 0 | 0 | 0 | 0 | 0 |
| Q9JKR6;F6TRP3;A0A1L1SQ3( | Hyou1 | Hypoxia up-regulated protein 1 | NaN | 0 | 0 | 167420 | 1363000 | 0 | 0 |
| Q9JKY0 | Rqcd1 | Cell differentiation protein RCD1 homolog | NaN | 0 | 0 | 0 | 0 | 0 | 0 |
| Q9JMG1 | Edf1 | Endothelial differentiation-related factor 1 | NaN | 0 | 0 | 0 | 0 | 0 | 0 |
| Q9JMH6-2;Q9JMH6;A0A1W2 | Txnrd1 | Thioredoxin reductase 1, cytoplasmic | NaN | 0 | 0 | 0 | 0 | 0 | 0 |
| Q9QUR6 | Prep | Prolyl endopeptidase | NaN | 0 | 0 | 0 | 0 | 0 | 0 |
| Q9QUR7 | Pin1 | Peptidyl-prolyl cis-trans isomerase NIMA-interacting 1 | NaN | 0 | 0 | 0 | 0 | 0 | 0 |
| Q9QZE5;Q7TNQ1 | Copg1 | Coatomer subunit gamma-1 | NaN | 0 | 0 | 0 | 0 | 0 | 0 |
| Q9QZM0 | Ubqln2 | Ubiquilin-2 | NaN | 0 | 0 | 0 | 0 | 0 | 0 |
| Q9QZS8-2;Q9QZS8 | Sh2d3c | SH2 domain-containing protein 3C | NaN | 0 | 0 | 0 | 0 | 0 | 0 |
| Q9QZZ4-3;Q9QZZ4-2;Q9QZZ | Myo15a;Myo15 | Unconventional myosin-XV | NaN | 0 | 0 | 0 | 0 | 0 | 0 |

| | | | | | | | | | |
|---|---|---|---|---|---|---|---|---|---|
| Q9R0E1 | Plod3 | Procollagen-lysine,2-oxoglutarate 5-dioxygenase 3 | NaN | 0 | 0 | 3206100 | 714420 | 0 | 0 |
| Q9R0H0-2;Q9R0H0 | Acox1 | Peroxisomal acyl-coenzyme A oxidase 1 | NaN | 0 | 0 | 0 | 0 | 0 | 0 |
| Q9R0X4 | Acot9 | Acyl-coenzyme A thioesterase 9, mitochondrial | NaN | 0 | 0 | 0 | 0 | 0 | 0 |
| Q9R190 | Mta2 | Metastasis-associated protein MTA2 | NaN | 0 | 0 | 0 | 0 | 0 | 0 |
| Q9R1E0 | Foxo1 | Forkhead box protein O1 | NaN | 0 | 0 | 0 | 0 | 0 | 0 |
| Q9R1J0 | Nsdhl | Sterol-4-alpha-carboxylate 3-dehydrogenase, decarboxylating | NaN | 0 | 0 | 0 | 0 | 0 | 0 |
| Q9R1P3 | Psmb2 | Proteasome subunit beta type-2 | NaN | 0 | 0 | 0 | 0 | 0 | 0 |
| Q9R1P4 | Psma1 | Proteasome subunit alpha type-1 | NaN | 0 | 0 | 0 | 0 | 0 | 0 |
| Q9R1Q6 | Tmem176b | Transmembrane protein 176B | NaN | 0 | 0 | 0 | 0 | 0 | 0 |
| Q9R1Q7 | Plp2 | Proteolipid protein 2 | NaN | 0 | 0 | 0 | 0 | 0 | 0 |
| Q9WTL7 | Lypla2 | Acyl-protein thioesterase 2 | NaN | 0 | 0 | 0 | 0 | 2609000 | 912800 |
| Q9WTP6-2;Q9WTP6;F7BP55 | Ak2 | se 2, mitochondrial;Adenylate kinase 2, mitochondrial, N-terminally processed | NaN | 0 | 0 | 0 | 0 | 0 | 0 |
| Q9WTR5 | Cdh13 | Cadherin-13 | NaN | 0 | 0 | 0 | 0 | 0 | 0 |
| Q9WTX5 | Skp1 | S-phase kinase-associated protein 1 | NaN | 0 | 0 | 0 | 0 | 0 | 0 |
| Q9WU78;Q9WU78-3;Q9WU | Pdcd6ip | Programmed cell death 6-interacting protein | NaN | 0 | 0 | 12016000 | 2785300 | 0 | 0 |
| Q9WUM4 | Coro1c | Coronin-1C | NaN | 0 | 0 | 0 | 0 | 0 | 0 |
| Q9WUU7 | Ctsz | Cathepsin Z | NaN | 0 | 0 | 0 | 0 | 0 | 0 |
| Q9WV66 | 07-Mar | E3 ubiquitin-protein ligase MARCH7 | NaN | 0 | 0 | 0 | 0 | 0 | 0 |
| Q9WVA2;Q4FZG7 | Timm8a1;Timm8a2 | anslocase subunit Tim8 A;Putative mitochondrial import inner membrane translocase | NaN | 0 | 0 | 0 | 0 | 0 | 0 |
| T1ECW4;Q9WVB0;Q9WVB0- | Rbpms | RNA-binding protein with multiple splicing | NaN | 0 | 0 | 0 | 0 | 0 | 0 |
| Q9Z0F7 | Sncg | Gamma-synuclein | NaN | 0 | 0 | 0 | 0 | 0 | 0 |
| Q9Z0J7 | Gdf15 | Growth/differentiation factor 15 | NaN | 0 | 0 | 0 | 0 | 0 | 0 |
| Q9Z0P4;Q9Z0P4-2;A0A1W2F | Palm | Paralemmin-1 | NaN | 0 | 0 | 0 | 0 | 0 | 0 |
| Q9Z110-2;Q9Z110;D3Z0B4 | Aldh18a1 | -carboxylate synthase;Glutamate 5-kinase;Gamma-glutamyl phosphate reductase | NaN | 0 | 0 | 0 | 0 | 2135900 | 2102600 |
| Q9Z131-3;Q9Z131-2;Q9Z131 | Sh3bp5 | SH3 domain-binding protein 5 | NaN | 0 | 0 | 0 | 0 | 0 | 0 |
| Q9Z1A1;F6QJV5;B8JJG9;B8JJ | Tfg | | NaN | 0 | 0 | 0 | 0 | 0 | 0 |
| Q9Z1G3 | Atp6v1c1 | V-type proton ATPase subunit C 1 | NaN | 0 | 0 | 0 | 0 | 0 | 0 |
| Q9Z1M8;A0A494BAW7 | Ik | Protein Red | NaN | 0 | 0 | 0 | 0 | 0 | 0 |
| Q9Z1N5 | Ddx39b | Spliceosome RNA helicase Ddx39b | NaN | 0 | 0 | 0 | 0 | 0 | 0 |
| Q9Z1Q5 | Clic1 | Chloride intracellular channel protein 1 | NaN | 0 | 0 | 0 | 0 | 0 | 0 |
| Q9Z1Z2 | Strap | Serine-threonine kinase receptor-associated protein | NaN | 0 | 0 | 0 | 0 | 0 | 0 |
| V9GXI0;V9GXT7;Q9Z266 | Snapin | SNARE-associated protein Snapin | NaN | 0 | 0 | 0 | 0 | 0 | 0 |
| Q9Z277-2;Q9Z277 | Baz1b | Tyrosine-protein kinase BAZ1B | NaN | 0 | 0 | 0 | 0 | 0 | 0 |
| Q9Z2A7 | Dgat1 | Diacylglycerol O-acyltransferase 1 | NaN | 0 | 0 | 0 | 0 | 0 | 0 |
| Q9Z2M7 | Pmm2 | Phosphomannomutase 2 | NaN | 0 | 0 | 0 | 0 | 0 | 0 |
| Q9Z2N8;A0A0A6YWR1;A0A0 | Actl6a | Actin-like protein 6A | NaN | 0 | 0 | 0 | 0 | 0 | 0 |
| Q9Z2Z6 | Slc25a20 | Mitochondrial carnitine/acylcarnitine carrier protein | NaN | 0 | 0 | 0 | 0 | 0 | 0 |
| Q9Z315 | Sart1 | U4/U6.U5 tri-snRNP-associated protein 1 | NaN | 0 | 0 | 0 | 0 | 0 | 0 |
