## Supplementary Table 2 for "The role of aging on endothelial cell-cell junctions and pulmonary microvascular permeability"

**Supplementary Table 2 |** Differentially enriched proteins from young mice associated with actin cytoskeleton.

| Gene name | Protein name |
|---|---|
| Stmn1 | stathmin 1 |
| Dync1h1 | dynein cytoplasmic 1 heavy chain 1 |
| Plec | plectin |
| Dbnl | drebrin-like |
| Hsp90ab1 | heat shock protein 90 alpha (cytosolic), class B member 1 |
| Lmnb1 | lamin B1 |
| Tln1 | talin 1 |
| Arpc3 | actin related protein 2/3 complex, subunit 3 |
| Tmsb4x | thymosin, beta 4, X chromosome |
| Pacsin2 | protein kinase C and casein kinase substrate in neurons 2 |
| Ezr | ezrin |
| Capza2 | capping actin protein of muscle Z-line subunit alpha 2 |
| Csrp1 | cysteine and glycine-rich protein 1 |
| Actb | actin, beta |
| Actr2 | actin related protein 2 |
| Capzb | capping actin protein of muscle Z-line subunit beta |
| Sptbn1 | spectrin beta, non-erythrocytic 1 |
| Hspg2 | perlecan (heparan sulfate proteoglycan 2) |
| Gsn | gelsolin |
| Marcksl1 | MARCKS-like 1 |
| Sntb2 | syntrophin, basic 2 |
| Vim | vimentin |
| Tmod3 | tropomodulin 3 |
