## Supplementary Table 3 for "The role of aging on endothelial cell-cell junctions and pulmonary microvascular permeability"

**Supplementary Table 3 |** Differentially enriched proteins from aged mice associated with actin cytoskeleton.

| Gene name | Protein name |
|---|---|
| Itga4 | integrin alpha 4 |
| Add1 | adducin 1 |
| Anxa2 | annexin A2 |
| Lmna | lamin A |
| Cnn2 | calponin 2 |
| Tpx2 | TPX2, microtubule-associated |
| Dpysl3 | dihydropyrimidinase-like 3 |
| Myo1c | myosin IC |
| Flnc | filamin C, gamma |
| Cav1 | caveolin 1, caveolae protein |
| Krt10 | keratin 10 |
| Tpm3 | tropomyosin 3, gamma |
| Tagln2 | transgelin 2 |
| Fn1 | fibronectin 1 |
| Tpm4 | tropomyosin 4 |
| Cald1 | caldesmon 1 |
| Actn4 | actinin alpha 4 |
| Pdlim7 | PDZ and LIM domain 7 |
| Pdlim1 | PDZ and LIM domain 1 (elfin) |
| Fus | fused in sarcoma |
| Gmfb | glia maturation factor, beta |
| Pfn1 | profilin 1 |
| Dstn | destrin |
| Ran | RAN, member RAS oncogene family |
| Ehd2 | EH-domain containing 2 |
| Cfl1 | cofilin 1, non-muscle |
| Flnb | filamin, beta |
| Syne1 | spectrin repeat containing, nuclear envelope 1 |
